# Large future genetic diversity losses are predicted even with habitat protection

**DOI:** 10.1101/2024.10.21.619096

**Authors:** Kristy S. Mualim, Jeffrey P. Spence, Clemens Weiß, Oliver Selmoni, Meixi Lin, Moises Exposito-Alonso

**Author notes:** Authors contributed equally (Kristy S. Mualim, Jeffrey P. Spence, Clemens Weiß).

## Abstract

Genetic diversity within species is the basis for evolutionary adaptive capacity and has recently been included as a target for protection in the United Nations’ Global Biodiversity Framework (GBF). However, we lack large-scale mathematical frameworks to quantify how much genetic diversity has already been lost, let alone to predict future losses under 21st century conservation scenarios. To fill this gap, we developed an area-based spatio-temporal predictive framework of genetic diversity calibrated with population-scale genomic data of 29 plant and animal species. To estimate present genetic diversity loss with our framework, we used species’ habitat area and population sizes losses reported in the Living Planet Index, the Red List, and new GBF indicators across 13,808 species for the last 5 decades. Applying our evolutionary framework across these species, we estimate genetic diversity loss lags behind population and habitat area declines, with an estimated current 13–22% π genetic diversity loss. However, we forecast future genetic diversity losses will reach 41–76% even if populations are not further contracted. These results highlight that safeguarding existing habitats is insufficient to maintain the genetic health of species and relying solely on continuous genetic monitoring underestimates lagging long term impacts.

**Significance statement:** Genetic diversity is crucial for both species adaptation and survival. Recently, it has been included as a target for protection in the United Nations’ Global Biodiversity Framework. However, we lack large-scale predictive methods to quantify current and future losses of genetic diversity across species. Here, we develop an area-based spatio-temporal predictive framework trained with high-quality genome-wide data from 29 plant and animal species to enable quantitative predictions of genetic biodiversity at global scales. We infer global genetic diversity losses are beyond the preliminary UN targets to protect 90% of genetic diversity of species, evolutionary models dramatic genetic losses will occur in the future even with habitat protection if populations across species are not recovered.

## Main Text

Genetic diversity dictates a species’ ability to adapt to new environmental conditions (1) and thus was coined as one of the three key dimensions of biodiversity at the 1992 United Nations’ Convention of Biological Diversity (CBD) (2); along with species and ecosystem level biodiversity. After being overlooked in global biodiversity framework texts for decades (3), the 2022 Kunming-Montreal UN CBD convention finally ratified a Global Biodiversity Framework (GBF, <u>CBD/COP/DEC/15/4</u>, <u>CBD/COP/DEC/15/5</u>) that explicitly recognizes to “*maintain and restore the genetic diversity within and between populations of native, wild and domesticated species to maintain their adaptive potential”* (4).

Conservation genetic approaches have focused on species at high extinction risk—e.g. species where the last populations and individuals remain—by managing populations to prevent complete depauperation of genetic diversity and avoid inbreeding, which leads to further extinction risk. For instance, helpful prescriptions include ensuring species have >500 effective individuals (*N_e_*) to safeguard genetic diversity over generations from stochastic genetic drift or *ex situ* breeding programs that mate individuals with different genetic variation (5). One may then conclude that species (or their component populations) with *N_e_* > 500 may not be at risk of losing genetic diversity. To the contrary, many non-threatened species have suffered major geographic area range and population losses (6, 7), which we expect must have major consequences of genetic diversity losses (8–10). The new post-2020 global biodiversity framework aims to protect all species (not only threatened ones) from genetic diversity loss and to maintain this key genetic dimension of biodiversity for present and future generations. However, while new genomic tools are applied to an ambitiously growing numbers of species (11–13), global predictive frameworks and estimates of genetic diversity loss across species are still limited, let alone methods to project temporal trajectories of genetic diversity in the 21^st^ century.

To monitor and protect genetic diversity across large numbers of species and over time, global conservation efforts have focused on proxy or qualitative *indicators* based on present species’ population sizes and habitat areas. Such proxies are based on the biological insight that the number of individuals within a species and its geographic distribution correlates with the genetic diversity of a species (14, 15). For example, species that are categorized as threatened, with rapid declines in their habitat or population sizes, often have lower genetic diversity—at least in vertebrates and mammals which are mostly studied (14, 16, 17). A proxy headline indicator adopted in the Global Biodiversity Framework proposed by the GEO-BON genetics working group is the proportion of populations with effective population size (*N_e_*) below 500, following the legacy of conservation genetics of threatened species. A second complementary indicator includes the total fraction of populations within a species that has been lost with respect to a recent past baseline. A third indicator of monitoring efforts of a country is the number of species with any genetic data. These three indicators were showcased recently in an effort to evaluate 100+ species for 9 countries (18). It has also been proposed that existing large population monitoring of species could serve as proxies: One such coordinated effort is the Red List from the International Union for Conservation of Nature (IUCN), which has classified ∼80,000 animal, fungi, and plant species into non-threatened, vulnerable, endangered, and critically endangered, using several thresholds of rapid species decline (i.e. few individuals, contracting geographic area, etc.). A second coordinated effort is the Living Planet Index (19), also a GBF indicator, which monitors 41,986 populations of 5,579 terrestrial animal [vertebrate] species.

These proxy indicators for genetic diversity are based on demographic rather than genetic DNA data, and thus deemed feasible, cheap, and scalable. However, by design they do not provide a quantitative metric of DNA diversity nor inform future losses. To calculate quantitative genetic metrics requires DNA sequencing of at least partial or entire genomes for multiple individuals or populations of a species. Such metrics include the number of DNA mutations within a species (i.e. allelic richness or segregating sites [*S*]), or average nucleotide genetic diversity (i.e. average number of mutation differences between two individuals of a species or nucleotide diversity [π]).

To bridge the gap that conservation policy is often proxy-based through area- or population metrics, we recently described a mutations-area relationship (MAR) power law: *M = cA^zmar^* . This links allelic richness or segregating sites or number of mutations (*M*) and its geographic range area (*A*) through a *z_MAR_* parameter characterizing spatial structure (8, 20) (note we avoid *S* notation for segregating sites to avoid confusion with the species area equation, SAR). Re-arranged, the equation *(A_present_ / A_past_)^zmar^* quantitatively predicts the percentage loss of genetic diversity with a percentage habitat area reduction within a species (*A_present_ / A_past_*), and could in principle translate proxy indicators into quantitative genetic diversity metrics. However, it is still unclear how different genetic diversity metrics behave in continuous geographic ranges with complex area losses (e.g. fragmentation) nor what are the long-term consequences that area losses and genetic drift have on the genetics of a species.

To develop predictions of genetic diversity losses in complex geographic landscapes for the 21st century, we build new spatio-temporal population genetic theory and continuous space forward-in-time simulations informed and validated by a large genome-wide DNA sequencing database of 9,804 geo-referenced individuals from 29 plant and animal species. Although classic population genetics has described genetic diversity at equilibrium through frameworks such as mutation-drift balance (21, 22), migration-drift equilibrium in spatial contexts (23, 24), and the effects of population bottlenecks (25), these models fall short in capturing the spatio-temporal non-equilibrium dynamics of genetic diversity, especially in complex geographic landscapes of species threatened with extinction. We then build a standard Wright-Fisher population genetic model with an arbitrary number of subpopulations in a two dimensional geographic grid connected by migration, i.e. a meta-population. To calculate genetic diversity, we use the fact that the average differences between two sampled individuals can be derived from allele frequencies of a subpopulation or group of populations. The key insight is that the dynamics of genetic diversity depend only on the first and second moments of allele frequencies across subpopulations over time, which obey a linear system of ordinary differential equations, even when we alter landscapes and remove subpopulations due to fragmentation (hereon termed *WFmoments*, see details in **Mathematical Appendix**). Solving these we can thus compute expected genetic diversity over time across a landscape. In parallel, we validate these insights through a simulation-based framework using SLiM v4.1 (26), under which we model non-Wright Fisher dynamics in a two dimensional geographic continuous space incorporating both age and spatial structure during mating (see **Materials and Methods, Text S1**, **Fig. 1-3**). In contrast to other theory and simulation approaches in conservation genetics (27, 28), we design our framework to not only be applicable to a threatened species with one or very few small populations, but it incorporates spatial structure and is amenable to large population sizes representing a broadly distributed species (e.g. simulation landscape runs are N_e_=5000-20000 individuals, and theory landscapes can be arbitrarily large, e.g. θ=4N_e_µ=0.01; N_e_ ≈ 2×10^6^). With this in hand, we study and simulate species with different population sizes and geographic landscapes, and evaluate the consequences to genetic diversity with different habitat and population loss scenarios (**Fig. 1A, Fig. S1-2, S7**).

**Fig. 1.**
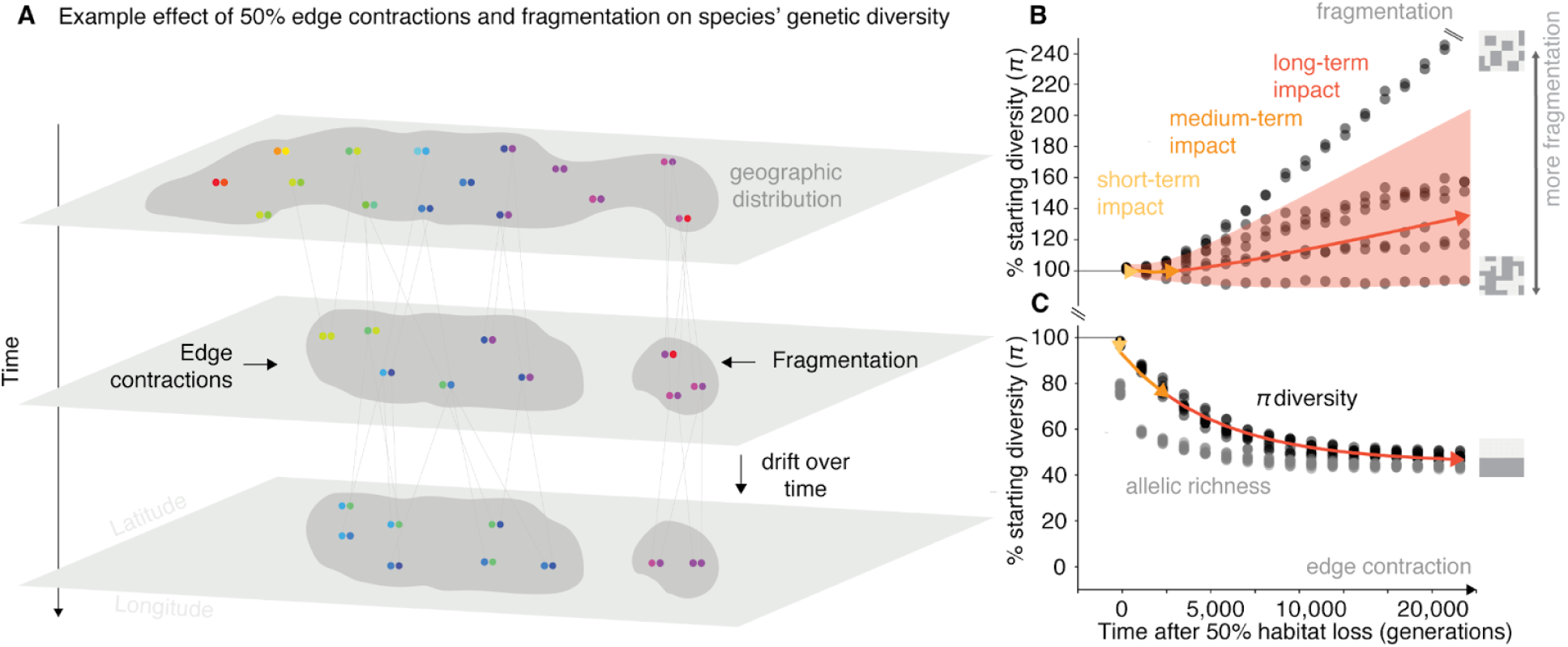
Genetic diversity metrics after habitat destruction. (**A**) Cartoon depicting a geographic distribution of a diploid species with spatial genetic structure (colors), the genealogy of genotypes, and how genetic diversity changes after area or population losses through edge contraction and fragmentation. (**B**) Genetic diversity as average pairwise nucleotide distances (π) and richness of alleles or mutations (*M*), trajectories after geographic range loss driven by habitat fragmentation over time. Each trajectory represents a replicate for 50% habitat loss over time, where each replicate is a different habitat fragmentation map. Habitat fragmentation maps vary according to level of connectivity between patches of landscape. Each black dot represents a sampled time point of π for that trajectory. Trajectories were tracked across 20,000 generations. Red arrow represents average long-term genetic diversity projection based on our theoretical framework, with shaded areas showing 95% confidence intervals. Orange and yellow arrows represent average medium-term and short-term genetic diversity projections based on our theoretical framework respectively (Species parameters: F_ST_ = 0.3 and θ = 10^−4^). (**C)** Genetic diversity, π, trajectories after geographic range loss driven by edge contraction over time.

We first ask how increasing spatial habitat destruction immediately impacts genetic diversity on a short-term timescale. A simple approach to simulate habitat destruction is removing area from one extinction edge, hereon termed edge contraction (**Fig. 2**). We found short-term loss of genetic diversity lags behind area loss in a predictable manner (simulation results predicted by *WFmoments*: R^2^ = 0.98, *P* < 1 × 10^−16^, n =81 **Fig. 2A**). The extent of this lag is determined by the migration and gene flow rates of the species in the landscape, i.e. the spatial population genetic structure (**Fig. S2**). For instance, a panmictic species with infinite gene flow and thus no population structure (F_ST_ ≈ 0) suffers only a 4.7% (95% CI = 4.5–4.9%) instantaneous nucleotide genetic diversity loss at 50% of habitat loss (**Fig. 2B**), while a species with high population structure (e.g. F_ST_ = 0.9) would suffer an instantaneous 9% (95% CI = 7.9–10.0%) nucleotide genetic diversity loss (**Fig. 2B**). These results are robust to the dimensions of the landscape (e.g. a 1 dimension landscape such as a river system, **Text S2, Fig. S22**) or to step-wise loss such as a 1% habitat loss per generation rather than instantaneous habitat loss (**Fig. S1**), suggesting short-term genetic diversity under straightforward edge contraction remains robust to the rate and shape of habitat loss. This recapitulates our power law findings of the mutations-area relationship (MAR) originally developed for allelic richness (8, 20) (**Text S4**). As suspected, in a species landscape of moderate population structure (F_ST_ = 0.3) a power law with a scaling *z_GDAR_* ∼ 0.03 also fits a nucleotide genetic-diversity-area relationship (GDAR), with high accuracy (R^2^=0.96) and a MAR with a scaling of *z_MAR_=* 0.3 (R^2^=0.98) (**Fig. S2, Tables S1-4**). Simulating extinction of sampled populations of empirical datasets of 29 plant and animal species also confirms MAR and GDAR have predictability in real world species (mean population structure of F_ST_ = 0.26, mean accuracy R^2^ = 0.677, range 0.10-0.98, **Table S18**). Our simulations and the power law intuitively explain why there is a lag of genetic diversity loss, as a power law that approaches *z→0* (i.e. low population structure) indicates genetic diversity should be relatively insensitive to habitat area losses: *(A_present_ / A_past_)^z→0^* ≈ 1 (**Fig. 2A, Fig S2-3**). Together, the MAR and GDAR power laws could thus be readily applicable to predict short-term losses of allelic richness and nucleotide diversity, respectively, across large numbers of species whose habitat is monitored.

**Fig. 2.**
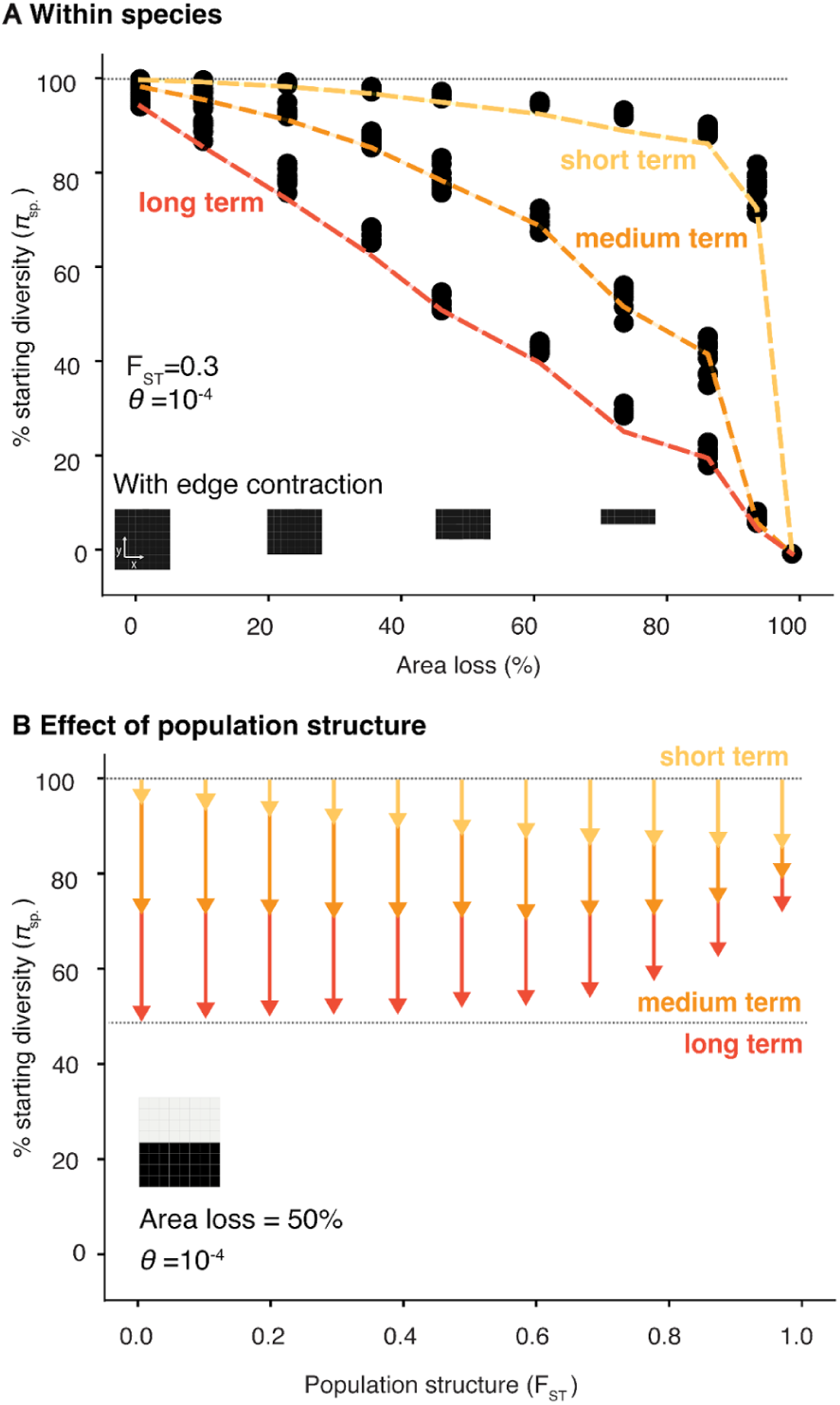
Predictable short- and long-term genetic diversity losses from edge contraction. (**A**) Landscape simulation (n=9 for each 10 area losses) of genetic diversity and theoretical expectations (lines) immediately after habitat loss (short-term, yellow), mid-way through habitat loss (medium-term, orange) and at final equilibrium (long-term, red). (**B**) The dependence of short- (yellow), medium- (orange) and long-term (red) losses with geographic population structure defined by average pairwise F_ST_ across 100 subpopulations (10 × 10 grid).

**Fig. 3.**
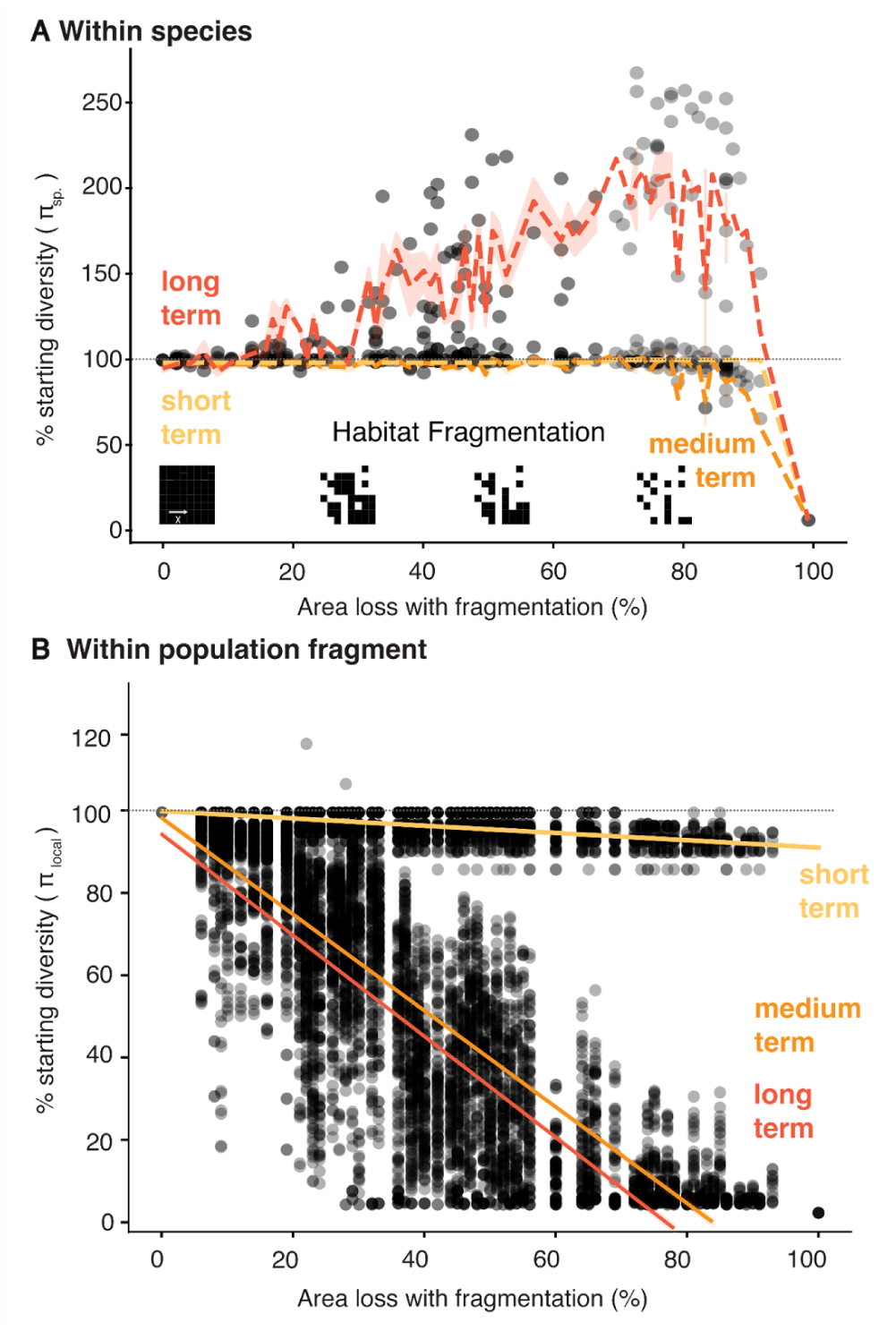
Impacts from habitat destruction by fragmentation are only detectable in within-population genetic diversity. (**A**) Overlay of theoretical and simulation-based projections of short-, medium- and long-term genetic diversity loss (π) due to habitat loss with fragmentation (121 random fragmentation maps). Each dot represents an average measure of π across a landscape per simulation and overlaid the walking average and 95% interval of short (yellow), medium (orange) and long-term (red) genetic diversity. (**B**) Within-population genetic diversity, π_local_, for the same simulations as (A). Dotted lines represent a linear model.

While it is expected that habitat loss has long-term impacts on genetic diversity (29), its magnitude in complex landscapes remains poorly understood. A classic prediction from population genetics assuming a single panmictic population (i.e. no spatial structure) is that the expected nucleotide diversity for diploids in equilibrium follows (30, 31): *E[π] = 4N_e_ μ.* If area or population abundance represents *N*_e_, a reduction in population size would thus proportionally reduce *π* towards a long-term equilibrium. By utilizing WFmoments to measure nucleotide diversity over time and tracking simulated populations for ∼20,000 generations, we investigated the non-equilibrium dynamics of genetic diversity loss just after habitat loss stops, monitoring its progression until a final equilibrium is reached (**Fig. 1-2, Fig. S1, S4**). We found that under edge contraction and in a species with no population structure (F_ST_ ≈ 0), the long-term equilibrium roughly follows and surpasses the classic expectation: that a ∼50% area loss roughly results in a ∼50% long-term genetic diversity π loss (**Fig. 2B**), with similar trends for allelic richness (**Fig. S15**). Counterintuitively, while increasing population structure led to larger short-term genetic diversity loss **(Fig. 2B**; yellow arrows), we found the opposite is true for both medium- and long-term loss (**Fig. 2B**; orange and red arrows, see theoretical interpretation **Text S3**); likely because structured landscapes can generally hold larger amounts of genetic diversity. However, both medium- and long-term genetic diversity loss equilibrium is unlikely to be reached in the near-term timeline of conservation policies for 2050. For instance, even for relatively fast growing organisms, from fruit flies (∼10-20 generations per year), to annual plants (∼1 generation per year), to fast growing trees (∼ 15 years per generation), the remaining ∼25 years would correspond to ∼400, ∼25, and ∼2 generations respectively (see scales in **Fig. 1B**). Nonetheless, species with moderate population sizes and fast generation times are being impacted within this century, while others will continue to lose diversity in longer timescales even if conservation practices succeed in stopping habitat destruction.

To avoid dramatic long-term losses, we need to restore habitat area and population sizes within species. The buildup of genetic diversity through new mutations is incredibly slow. Using our SLiM framework, we simulated two scenarios: a habitat recovery (letting populations naturally colonize it) and population restoration (individuals across the intact landscape are moved to the restored habitat) (**Fig. S23**, **Text S5**). With this, we were able to see a significant lag in genetic diversity recovery (**Fig. S24-25**). For short-lived species, recovery is faster. For slow growing species, we must reduce initial demographic impacts and restore populations as soon as possible since recovery will be unlikely in the next centuries.

Habitat destruction and geographic area losses within a species may not occur from an edge. Rather, habitat may become fragmented, which poses unique risks for species (32). Fragmentation alters gene flow, and increases isolation and exposure of habitat edges to genetic drift. Using our SLiM and *WFmoments* frameworks, we stochastically remove habitat grids, sized at 1/100^th^ of the landscape, allowing for isolated populations that have no gene flow with neighbors (**Fig. 1B, 3A, Fig. S7**). Simulations showed species-wide nucleotide genetic diversity is highly insensitive to fragmentation in the short-term and in fact dramatically increases in the long-term: for instance, with ∼90% habitat loss, we observed an increase of 263% of nucleotide genetic diversity (range: 100-263%; also observed in allelic richness, **Fig. S14**). While seemingly counter-intuitive, this is a longstanding observation in population genetics coined the “Wahlund effect”, where lack of gene flow between two previously connected populations leads to genetic drift that fixes independent alleles, causing populations to become more distinct. Thus, increased diversity is observed if populations are pooled (33, 34) (**Fig. S14, S15**). This genetic diversity inflation could consequently be predictable, and in fact, the *WFmoments* expectations in fragmented 2D landscapes recapitulate diversity trends in simulations (R^2^ = 0.46, *P* = 4.46 × 10^−244^, n = 121 **Fig. 2A, Fig. S22**). This inflation effect is repeatable with smaller (1/400^th^ of the landscape) and larger (1/12^th^) fragmentation, and is explained by landscape fragmentation metrics of different simulations, including total core area (R^2^ = 0.76, *P* = 1.62× 10^−25^), spatial connectedness (R^2^ = 0.87, *P* = 1.11× 10^−35^), the general perimeter of patches (R^2^ = 0.53, *P* = 2.49× 10^−14^), and patch and edge density (R^2^ = 0.63, *P* = 2.12× 10^−18^) **(Fig. S10-11)**.

Although fragmentation and isolation of populations in a species may cause them to diverge and inflate species-wide genetic diversity, this is hardly indicative of good genetic health. In fact, by quantifying genetic diversity within population fragments in our simulations, we detect a small but significant reduction in the short term (linear regression *b* = -0.09, *P* = 2.57 × 10^−236^, R^2^ = 0.30, **Fig. 3B**), a large medium-term reduction (linear regression *b* = -1.20, *P* < 1.0 × 10^−16^, R^2^ = 0.69, **Fig. 3B)** and an even larger long-term reduction (linear regression *b* = -1.26, *P* < 1.0 × 10^−16^, R^2^ = 0.72, **Fig. 3B, Fig. S14**), with more dramatic results for allelic richness (**Fig. S14, S15**). The implications of our study of species area reductions with severe isolations are twofold. First, genetic diversity protection goals and indicators within the Global Biodiversity Framework must include explicit guidelines on how genetic diversity metrics are calculated, and these metrics have to be spatially-aware (i.e. genetic diversity computed in individuals from different isolated populations may result in misleading values). Second, the combination of area loss and fragmentation may challenge the application of easy-to-use power law equations, MAR and GDAR, which use only area loss. In contrast, the *WFmoments* framework can achieve accurate predictions of within- and across-population diversity if census monitoring frameworks not only focus on population and area sizes but also provide landscape connectivity metrics.

Finally, we use our power law functions GDAR/MAR and *WFmoments* approximation tables (see **Supplementary Data**) to estimate global genetic diversity loss due to land use change and habitat destruction to date. Although accurate species-specific geographic area reduction data in the past centuries are scarce for many species, we leverage existing demography indicators to serve as proxies for habitat transformations and showcase how we can bridge this information with population genetic theory to achieve estimates of genetic diversity loss.

The first prediction uses species assessment documents from the IUCN Red List, which contains habitat area and population size decline information within at least 10 years / 3 generations, especially under classification criteria A1-A4c and C1 (8, 35, 36) (while other criteria are also relevant for extinction risk due to small ranges and populations, such as B1-2 and D, they rely on fixed thresholds, such as fewer than 1,000 individuals, without considering trends over time.). We must emphasize that we do not expect genetic diversity losses to only apply to threatened species where population sizes are close to the typical N_e_ 500 limit used in conservation genetics (a minority of species 1,800 out of 82,798 have less than N<1000 census individuals; Red List criterion D, **Table S20**, while the majority of genetic biodiversity losses is attributed to many thousands of non-threatened yet declining species, ∼65% of Red List species are least concerned or near threatened, but 65% of those have a demographic decline). Hence, we attempt to translate assessments of all 82,798 plant, animal, and fungi species of estimated threat categories to genetic diversity loss (**Fig. S18**, **Table S20)**. For instance, focusing on A2-4c criteria only: there were 2,240 *vulnerable* species which suffered ≥30% area loss (*1-A_present_/A_past_*), 1,621 *endangered* species that lost ≥50% of area, 916 *critically* endangered species that lost ≥80% of area, and 1,688 that did not pass the ≥25% area loss threshold that have thus been classified as least concern (other criteria that contain area information can also be used, **Fig. S18**, **Table S20)**. Using MAR/GDAR, we transform these area loss ranges into expected genetic diversity loss. In cases where genetic data (F_ST_, *z_MAR_, z_GDAR_*) is not available, we utilize ranges from species with empirical data: *F_ST_=*0.01–0.6, *z_MAR_* =0.01–0.8, and *z_GDAR_* =0.01–0.8 (**Fig. 4A**, **Fig. 4B-E, Table S20**). One such species of least concern, *Eucalyptus melliodora*, is a common tree of Australia for which we have genomic data (**Fig. 4A**, **Fig. S17, Table S19**). It was assessed as vulnerable under criterion A2c, meaning its area loss must exceed 30% but remain below 50%; otherwise, it would be classified as endangered. With this, its minimum expected allelic genetic diversity loss using MAR is equivalent to: *1-(0.3)^0.3^ = 10%;* and a maximum of *1-(0.5)^0.3^ = 19%;* (midpoint reported in **Fig. 4A**). The GDAR approach can be applied to predict nucleotide genetic diversity loss (π) by substituting *z_GDAR_* values in the same MAR equation (**Table S19**). Equivalently, we can use the species’s average *F_ST_* along with our *WFmoments* pre-computed loss tables. For *E. melliodora*, a *F_ST_ =0.01* and area loss of 30-50%, would incur in a short-term nucleotide diversity loss of 0.15% (IQR=0.11–0.18%) and a long-term loss of 39.5% (IQR=34.5–44.5%). Applying these approaches across 7,263 species evaluated in the Red List with A1-4 criteria, we find a GDAR/*WFmoments*-based average short-term nucleotide genetic diversity loss of 12.8% (IQR=0–17.3%), and MAR-based allelic richness loss of 28.8% (IQR=1.4–44.1%) (**Fig. 4B**). Assuming no further habitat losses in the future we predict using *WFmoments* that the lagged dynamics of genetic diversity will continue increasing losses to a medium-term nucleotide diversity average of 17.8% (IQR=2.3–24.5%) (**Fig. 4C**) before reaching a final long-term equilibrium of 41.3% (IQR=12.8–64.1%).

**Fig. 4.**
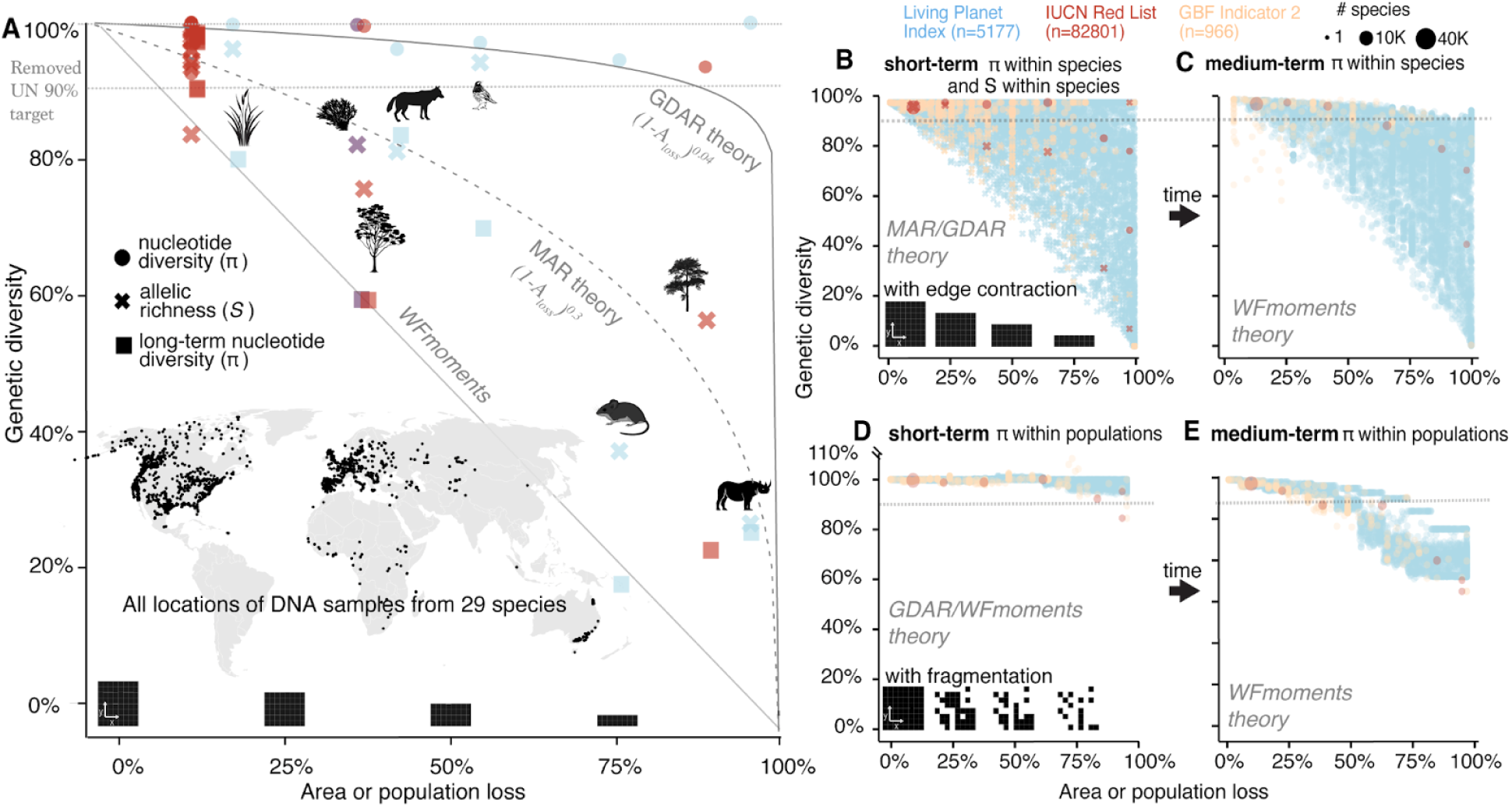
Estimates of genetic diversity loss for empirical, LPI and Red List species over both the short and long-term. (**A**) Estimates of genetic diversity loss for 29 plant and animal species for which genomic information is also available to calibrate MAR/GDAR/*WFmoments* predictions. Using per species *z_MAR_* parameter and the MAR equation, we predict short-term allelic richness (crosses) losses. Using per species *F_ST_* or *z_GDAR_* and GDAR and *WFmoments*, we predict short-term nucleotide genetic diversity losses (π, dots). Finally, using *WFmoments* we also predict long-term nucleotide genetic diversity losses (π, squares). Area or population losses per species are extracted from Red List category (red), Living Planet Index (blue), or additional expert range (purple). Predictions follow the edge contraction expectations. Dotted line at 90% represents the now-removed preliminary UN genetic diversity target of 90%. (**B**) Genetic diversity loss projections for 13,808 species for which recent area or population loss information is available from Red List (red), LPI (blue), or new GBF indicators (orange). Predictions of allelic richness (crosses) with MAR use randomly sampled *z_MAR_* parameters for 13,808 species following distribution of known species in (A), and predictions of nucleotide diversity π (dots) with *WFmoments* follow the same approach sampling *F_ST_*. Sizes of dots represent numbers of species, as in Red List many species have the same threatened category and there are no species-specific area losses. (**C**) Long term projections of genetic diversity loss following the short-term reduction in (B) assuming area and population losses stop. (**D-E**) Genetic diversity projections for the same 13,808 species as in (B) assuming area loss with high fragmentation and isolation using *WFmoments* in the short-term (D) and long-term (E).

For the second prediction, we use the comprehensive demographic survey data across 5,579 threatened and threatened and non-threatened vertebrate species from the Living Planet Index (LPI) (37), to compute population decline losses per species, with a specific focus on within-species average census losses: 64% (IQR=40–93%) (**Fig. S16**, see others in **Materials and Methods**). As before, using GDAR/MAR, we estimate that across 4 decades of data there is an average population size decline per species that corresponds to a nucleotide genetic diversity loss (π) on average 15% (IQR=0–14.5%) **(Fig. 4B)** while average allelic richness loss is 36.5% (IQR=7.5–60%) (**Fig. 4B**). Medium-term projections with *WFmoments* estimate a 16.2% nucleotide diversity loss (IQR= 10.5–20.4%) **(Fig. 4C)** and equilibration to long-term estimates of a 45% nucleotide diversity loss (IQR = 20–68%).

Third, the GEOBON Genetic Composition group showcased the feasibility of 2022’s GBF genetic proxy indicators by carefully annotating 966 species for 9 countries (38) (**Fig. S19**). The fraction of wild populations lost (Indicator 2) per species averaged 18.6% (IQR=0–35%); of which 76% (IQR=60–100%) of the remaining populations had over 500 N_e_ (Indicator 1) (note that N_e_ is assumed (39) to be 10% of individual census size N_c_, although there are discrepancies on how variable N_e_/N_c_ ratios can vary several orders of magnitude across vertebrate species (17), (40)). As before, using approximated population loss from indicator 2 across species in combination with the MAR/GDAR/WFmoments framework, we estimate an average nucleotide diversity loss of 1.5% (IQR=0%) and allelic richness loss of 7.2% (IQR=0–7.5%) in the short-term, 2.4% (IQR=0–4.3%) nucleotide diversity loss in the medium term, and 6% (IQR–1-2%) nucleotide diversity loss in the long term. For now, it is unclear how indicator 1 may be used for population genetic predictions, as N_e_ within a population not only depends on N_c_ but also on gene flow, migration and connectivity between populations (e.g. at equal N_c_, increased gene flow increases local N_e_). In the absence of a complementary quantitative demographic value, this qualitative indicator is helpful in prioritizing efforts and detecting imminent population and genetic diversity losses (39). For instance, combining indicator 1 and 2, we anticipate that the 26.7% of populations which are below N_e_ < 500 will likely become extinct soon. Adding this to the already lost 17.6% of populations per species would result in a potential 39% of total populations lost, a potential mean short-term nucleotide diversity loss of 41% (IQR=0–100%) and an allelic richness loss of 63% (IQR=9–100%).

All above predictions are based on the robust edge contraction patterns (**Fig. 2**), but we expect fragmentation to limit genetic diversity loss predictability in certain scenarios. To explore these challenges, we also conduct predictions assuming that monitored species lose populations with high fragmentation. Because we lack quantitative connectivity metrics of populations evaluated in LPI, Red List, and GEOBON-driven GBF indicators, we utilize *WFmoments* and assume high random fragmentation and isolation levels of our simulations (**Fig. 1, 3**). Since we observe that in fragmented landscapes within-population or local genetic diversity may be most informative (**Fig. 3**), we summarize predictions of within-population genetic diversity (**Fig. 4D-E**). These show a 2% nucleotide diversity loss (IQR=0–3.9%) in the short term, a 16% nucleotide diversity loss (IQR=2.4–31%) in the medium term and a 71% nucleotide diversity loss (IQR=49–95%) in the long term. All in all, even under complex fragmentation scenarios, across prediction approaches and diversity metrics, we expect to have already lost moderate levels of genetic diversity. These genetic diversity losses will become more dramatic over the next decades even if no more populations or habitats are lost.

These first estimates of present and future global genetic diversity lost percentages across thousands of species solidify our previous warnings of major genetic biodiversity declines in the Anthropocene (8). On average, many species have likely already started losing genetic diversity, even those with populations of *N_e_ >* 500. Such losses may be hard to measure empirically in shorter time-scales (10, 29, 41) but become much larger over time. To achieve the new broad GBF genetic diversity targets, we urgently need data-driven and theoretically-informed approaches that recognize several biological principles of spatiotemporal dynamics of genetic diversity. Our new insights signal an urgent need for action and provide both a tool and a window of opportunity to ambitiously recover and reconnect populations, ensuring species’ long-term genetic protection.

## ADDITIONAL INFORMATION

### Author contribution

K.S.M., J.P.S., C.L.W., and M.E.-A. designed the project. K.S.M., J.P.S., C.L.W., O.S., M.L., and M.E.-A.conducted analyses and research. J.P.S. derived Wright Fisher moments theory. C.L.W. and K.S.M. built and run the simulation engine. O.S., M.L. K.S.M., M.E.-A. analyzed species data. K.S.M. prepared the first manuscript with M.E.-A.. All authors interpreted the results and wrote the paper.

### Data availability

Scripts will be deposited publicly at https://github.com/kmualim/spatial_extinction_sim WFmoments is available at https://github.com/jeffspence/wfmoments.

### Competing interest

The authors declare no competing financial interests. The funders had no role in study design, data collection and analysis, decision to publish, or preparation of the manuscript.

## Acknowledgements

We are grateful to the genomics community for making datasets publicly available and global conservation organizations for sharing publicly species monitoring data. We thank members of the Moi lab for comments, discussion, and references. We thank Sean Hoban for references and sharing new indicator data. J.P.S. is supported by an NIH training grant (5T32HG000044-23), C.L.W. is supported by Stanford’s Center for Computational, Evolutionary, and Human Genomics. M.L. is supported by the Smiths Fellowship from the Society of Conservation Biology. M.E.-A. and this research is supported by the Office of the Director of the National Institutes of Health’s Early Investigator Award with award number: 1DP5OD029506-01; by the U.S. Department of Energy, Office of Biological and Environmental Research, grant number: DE-SC0021286; and by the Carnegie Institution for Science, by the University of California Berkeley, and by the Howard Hughes Medical Institute.

## Methods summary

### Continuous-space population genomic simulations with SLiM

To study genetic diversity under different scenarios of species ranges and extinctions, we set up forward simulations in continuous 2D space using SLiM v. 4.0.1 (26). We simulated diploid genomes using a single, 10^8^ long chromosome, with recombination rate and mutation rate set at 10^−8^. Utilizing SLiM’s non-Wright-Fisher mode, we simulate a single population, the size of which is maintained through density dependent fitness effects given a carrying capacity K, which limits the census of individuals in the landscape, N_c_. Spatial structure is established by associating each individual with a continuous 2D coordinate (i.e. latitude and longitude), and by using these coordinates to govern three demographic processes: mate choice, dispersal, and competition. For mate choice and reproduction, if the focal individual is of fertile age, a mate is chosen randomly and weighted by spatial proximity, to generate a number of offspring sampled from a poisson distribution parameterized to control average fertility. A newly generated offspring’s position is drawn from a Gaussian distribution centered at the location of the maternal individual with a standard deviation of dispersal rate. In addition to this juvenile dispersal, all individuals alive at the end of a simulation tick disperse randomly, using twice the juvenile dispersal rate. The effect of spatial competition is based on the local population density felt by each individual. This is established using a Gaussian interaction to govern the strength of spatial competition, which is then rescaled based on the carrying capacity to keep the population at a size parameterized by the carrying capacity. During habitat reduction, we scale the carrying capacity by a function of habitat size to ensure that a reduction of habitat by 50% also leads to a 50% reduction in carrying capacity of the entire habitat. In addition to the density-dependent fitness effect, we also scale an individual’s fitness non-linearly with its age, up to a maximum age of 10 ticks. Together with a fertile age of 3 ticks, and a possion fertility rate of 0.5, these parameters set up the age structure of our population. Finally, note that the effective population size in population genetics, N_e_, is not a set parameter but an emergent parameter that may differ from census size N_c_ in non-panmictic, spatially-structured populations with complex age structures, reproduction, and overlapping generations (all complexities expected in nature). Unless otherwise specified, our simulations utilize a K = 5000 (i.e. carrying capacity N_c_) and a dispersal rate of 0.05.

To allow time for spatial population structure to develop, we allow this population to evolve forward in time for 1,000,000 SLiM ticks (∼200,000 generations) (**Fig. S20**). As predicted by the isolation by distance pattern, individuals sampled closely together in 2D space are now more genetically related than individuals sampled far apart. Rather than simulating every mutation, which is a major computational burden, we use tree sequence recording (42) to track the full genealogy of all individuals in the simulation which are either alive at the end of the simulation or sampled through time using the treeSeqRememberIndividuals function of SLiM. For the purposes of our simulation, we simulate 1,000,000 SLiM ticks (∼200,000 generations) to establish spatial structure and ensure that all sampled individuals fully coalesce (in which case the value of genetic diversity resulting from these simulations would be at equilibrium) (**Fig. S20**). To be able to start simulations with a large population in space that is at equilibrium without wasting computational resources, we simulated coalescence backwards in time with msprime (42). This process has been referred to as “recapitation” (43), where an incomplete genealogy of a large population with multiple roots (from SLiM) is “recapitated” using coalescent simulation backwards in time. This is made possible by using the tree sequence data structure to record and simulate genealogies in both SLiM and msprime. Since our simulation is only concerned with how processes such as dispersal affect neutral variation across space and through time, we can use the “recapitated” tree sequence to overlay mutations onto the full genealogy of all sampled individuals. The rationale being that under neutrality, mutations will not affect the structure of the genealogy, so we can simulate the genealogy without mutations first, before overlaying neutral mutations to reduce computational burden. We then extracted the resulting genotypes of all individuals from the tree sequence for downstream analysis. We partitioned the continuous-space simulation landscape in SLiM into a 10×10 grid for subpopulation sampling. For each grid, we calculated the pairwise genetic diversity, both π and number of segregating sites, between individuals in all 100 spatial grids. In addition, we tracked each individual’s location within spatial grids and the total number of segregating sites at each sampling time point. In further downstream analysis, we approximate that each generation corresponds to 4.5 SLiM time points as we simulate realistic age-structured populations with overlapping generations.

### Genetic diversity metrics

Genetic diversity is measured as π, the average pairwise difference between all possible pairs of individuals, and S, allelic richness (or the number of segregating sites). In addition, we proceed to calculate both within-population (one grid cell) and species-wide metrics of both π and S. We measured genetic diversity using pairwise genetic diversity with the equation below, where n is the number of remaining individuals, L is the total number of SNPs selected, and p_i_ is the allele frequency at genomic location i. The percentage of area loss was calculated by dividing the total number of map cells removed with the number of total map cells constructed from all geo-referenced individuals at each iteration. Genetic diversity was estimated as: *π = (n/n-1) (1/L) Σ ^L^ 2p_i_(1-p_i_)*.

#### Within-population genetic diversity metrics

Within-population genetic diversity metrics calculate the average pairwise difference between all possible pairs of individuals (π_local_) within each grid cell across all 100 grid cells. In total, we obtain 100 values of π or S, and proceed to calculate the average of π or allelic richness across all 100 grid cells. This ensures that individuals that belong in the same population are compared with each other.

#### Species-wide genetic diversity metrics

Species-wide metrics (π_species_) are calculated by getting the average pairwise difference between all possible pairs of individuals within all grids, so individuals–possibly in separate grids–are compared to each other to obtain an average value for all 100 grid cells.

### Simulating different scenarios of human impacts on species habitats

Our simulations included two ways of habitat destruction, habitat loss from one leading edge, and habitat loss with fragmentation.

#### Instantaneous range contraction simulations

To assess the dynamics of genetic diversity in populations undergoing habitat loss, we performed range contraction simulations starting from 10% habitat loss to 90% habitat loss at 10% increments, with 9 replicates at each progression of habitat loss. Each simulation was run for 1,000,000 SLiM time points before instantaneous habitat loss occurred at 1,000,001 SLiM time points. After which, we tracked the dynamics of both π and allelic richness in the short-term (at 1,000,001 SLiM time point after habitat loss), in the medium-term (at 1,010,025 SLiM time points or approximately 2,200 generations after habitat loss) and in the long-term (at 1,062,001 SLiM time points or approximately 13,800 generations after habitat loss). Unless otherwise specified, we utilized a census population size of N_c_ = 5,000, a dispersal rate of 0.05, mutation rate of 10^−8^ and recombination rate of 10^−8^.

Under habitat loss with fragmentation, we performed habitat loss simulations from 6% to 93%. We utilized a random integer generator to select which grids in our 10×10 simulation map to “extinct”, hence causing the range of habitat loss percentages to be variable. Each simulation run generates a unique habitat fragmentation map. In total, we simulated 121 different habitat loss maps with fragmentation. Unless otherwise noted, we utilize a 10×10 simulation map for our simulations.

In order to briefly explore other habitat fragmentation scenarios, we performed habitat loss simulations on a larger geographical grid of 20×20. Utilizing a random integer generator to select grids in our 20×20 simulation map to “extinct”, this simulation setup ensures that habitat loss is distributed more evenly across the geographical grid and at smaller areas than previously implemented. We term this as “mini” fragmentation to understand how genetic diversity changes with time if fragmentation occurs at much smaller scales across a wider geographical range.

#### Altering parameters in spatial population dynamics

Given the expectation that key population genetic parameters affect the dynamics of how genetic diversity changes with habitat loss, we wanted to explore how population size and migration rate might affect π diversity estimates under various habitat loss scenarios. We utilized population sizes ranging from 500 to 10,000 (exceptionally 20,000 individuals for computational efficiency) and varied dispersal rate respectively to test our variable Fst values to mimic low, medium and high population structure. Specifics of combinations of population sizes and dispersal rates can be found listed in the legend of **Fig S1**.

#### Gradual habitat loss simulations

To explore alternative habitat loss scenarios, we also looked at gradual habitat loss dynamics where habitat loss would decrease by 1% every 11 generations. This simulation was run for 2,000 SLiM time points before gradual habitat extinction of 50% habitat loss which occurred from 2,001 SLiM timepoints (approximately 450 generations) to 2,250 SLiM timepoints (500 generations).

### Fitting mathematical theory to simulations

Our modeling (see **Mathematical Appendix**) has three free parameters: effective population size, mutation rate, and migration rate. Under standard Wright-Fisher dynamics these may all be derived from real-world observable data, namely the census population size, rate of new mutations per generation, and number of migrants per generation. Yet, our simulations contain a number of features that strongly deviate from the standard Wright-Fisher model. There are overlapping generations, individuals live in continuous space instead of discrete demes, and there is density-dependent selection and spatially controlled mate choice. The Wright-Fisher model is surprisingly robust to deviations from its assumptions with the caveat that parameters must now be interpreted in terms of an “effective population size” that will depend on the actual model in complex ways (44). As a result, we expect our mathematical model to be able to recapitulate key features of the simulation, albeit with modified parameters.

To find the parameters that provided a good fit between our theory and our simulations we used least-squares fitting. We approximated the continuous habitat in the simulations by a 10 by 10 grid of square demes, with migration between adjacent demes. We then fixed the effective population size and optimized the least-squares fit between the theoretically predicted species-wide π from simulations and theory over three parameters: the mutation rate, the migration rate between adjacent demes, and a time-scaling for converting units of time in our theoretical model to SLiM timepoints in the simulations. Optimization was performed using the “Powell” method in scipy.optimize (45–47).

### Reanalyses of population genomic datasets of 29 species

#### Generating F_st_ values across diverse species

We utilized datasets across diverse species that were collected for (8) and (48). All datasets were transformed into PLINK files using PLINK v1.9 (49). For additional information on data processing, refer to the Supplementary materials of (8). For computational efficiency, we generated F_st_ values using admixture v1.3.0 (50), specifically utilizing the command ‘admixture –cv sample.bed <K or number-of-clusters>’ where we tested a range of 1 to 15 K and picked the K with the lowest CV error. We reported the average and maximum F_st_ values across K populations for each species (**Table S19**).

#### Simulating short-term extinction in empirical datasets

To confirm the genetic extinction patterns we observed in population genomic simulations and mathematical models, we also simulated short-term extinctions in empirical sequencing datasets of nineteen wild plant and animal species (detailed description of datasets available in (8)). Briefly, for each species, the empirical dataset contains geo-referenced individuals broadly sampled across its geographical distribution and naturally occurring mutations were discovered through various DNA sequencing methods. We conducted the analyses with up to 10,000 randomly selected biallelic SNPs for each species sampled genome-wide, or in the largest chromosome for those species with large genomes. We sought to use unfiltered SNP datasets to avoid ascertainment biases.

For each species, the *in silico* extinction simulations were conducted by iteratively removing map cells in the sample map, and geo-referenced individuals falling within the removed cells are considered extinct. Genetic diversity was estimated from the genotype matrix of remaining individuals using the R package MAR v0.0.3. We measured genetic diversity using pairwise genetic diversity with the equation below, where n is the number of remaining individuals, L is the total number of SNPs selected, and p_i_ is the allele frequency at genomic location i. The percentage of area loss was calculated by dividing the total number of map cells removed with the number of total map cells constructed from all geo-referenced individuals at each iteration. Genetic diversity was estimated as: *π = (n/n-1) (1/L) Σ_i_^L^ 2p_i_(1-p_i_)*.

We implemented two hypothesized patterns of extinction to match the population genomic simulations presented in the main text **(Table S5, Table S6)**. The random scheme involved extinctions occurring randomly across the range as would be expected by the habitat fragmentation scenario, while the south-north scheme had extinctions starting in the north and moving southward simulating the impacts of climate change. Each species underwent 20 replicates for both the random and south-north extinction schemes.

### Estimating global genetic diversity loss based on global population and area loss census

#### Living Planet Index

In order to estimate global genetic diversity loss, we extracted population sizes of 5,230 species that have been tracked for over 3 decades via the Living Planet Index 2024 (37). For each species, we summarized the population data in a few ways. To obtain the fraction of populations lost, we obtained the arithmetic mean of populations over 3 decades. For more information on the distribution of LPI data used, see **Fig. S16**. Given complexities in measuring population declines using time-series data (51), we opted to calculate the average arithmetic abundance decline from earliest to latest census (100 *× N_present_ / N_past_* +1) across populations within a species. Finally, we subsetted the LPI data to only estimate genetic diversity loss for species that experienced a decline in average arithmetic abundance across populations. This left us with 3,417 species. We then utilize these values as a proxy for habitat loss.

We assumed that F_st_ across LPI species would follow a normal distribution with mean 0.270 and standard deviation 0.211 (as determined by the F_st_ values we obtained across diverse species, see **Table S19**). Utilizing these F_st_ and habitat loss approximations, we estimated the amount of short-, medium- and long-term genetic diversity utilizing our theoretical framework (see **Supplementary data**). To obtain an average global genetic diversity loss estimate, we summed the total genetic diversity loss to obtain the total amount of genetic diversity loss divided by all populations considered.

#### Red list

We obtained criteria of area loss from the Red List database (www.iucnredlist.org). Counts of each category and criteria used in Red List classification are summarized (see **Supplementary data**). Genetic diversity loss values were calculated for all Red List categories: Extinct, Likely Extinct, Critically endangered, Endangered, Vulnerable, Near Threatened and Least Concern. We approximated area loss by using population size loss criteria for each of the 6 categories by taking the arithmetic mean of the minimum and maximum of the range. For example, a species is categorized as critically endangered when population size loss is between 80 to 95%. Utilizing these approximated area loss values for each category, we then used our theoretical and simulation based framework to make predictions of short, medium- and long-term genetic diversity π (see **Supplementary data**).

#### New KM GBF proxy indicators for genetic diversity

We obtained Indicator 2 calculations from (52). We calculated Indicator 2 using data from the Living Planet Index (see **Supplementary data** for details, **Fig. S19**). To obtain the fraction of populations lost in the LPI, we utilized the mean of populations that have reached 0 over 3 decades.

## Supplemental Materials

### Supplementary Text

#### Text S1 Simulation model assumptions and limitations

Our simulation-based predictions make a series of assumptions such as: (1) species are at equilibrium prior to any habitat loss; and (2) no further habitat loss occurs after the initial reduction. These projections will likely change when considering fluctuating population sizes, further changes in habitat, or changing migration patterns **(Fig. S3, Fig. S24, Fig. S25)**. Prior to altering the habitat of a species, we ensured that the species was at equilibrium by allowing the simulations to run for a defined burn-in rate of 1,000,000 SLiM timepoints. This was done to ensure that π was at a stable equilibrium before range contraction occurred **(Fig. S23)**.

##### Species at equilibrium

Initially, we performed simulations of edge contraction without sufficient burn-in (species are not at equilibrium), which caused genetic diversity trajectories across different percentages of habitat loss to become highly variable in the long-term **(Fig. S20, S21)**. This is likely going to be more reflective of how existing populations are in the wild and hence, further experiments should be performed to calculate how the dynamics of genetic diversity changes for a species not at equilibrium under habitat loss.

##### Constant population size

To address the second assumption of our model, we only induce just one bottleneck event (habitat range loss) before allowing the population to reach equilibrium over time. We assume that long-term constant population size from the time of habitat loss to the next equilibrium. This is designed for us to understand how habitat loss impacts genetic diversity immediately after habitat loss as well as across generations. However, it is likely that species in the wild will encounter frequent bottleneck events which would cause these genetic diversity trajectories to vary from those predicted in this work. We recommend further simulations in understanding how repeated bottleneck events may impact genetic diversity trajectories in the wild. It’s likely that these projections will become increasingly complicated, such as those seen in under habitat fragmentation **(Fig 2)**.

Neutral genetic diversity is heavily dependent on both mutation rate and population size and under the assumption of an infinite sites model in a Wright-Fisher (WF) model, it can be expressed as:

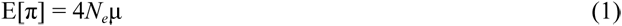

In theory, *N_e_* can be understood as a key population genetic parameter that determines the number of breeding individuals in an idealized population that shows the same level of genetic drift or inbreeding as the observed population. *N_e_* affects the rate of loss of genetic diversity and inbreeding and is an emergent property in our simulations. Accordingly, across generations, we expect that contemporary N_e_ will fluctuate according to the number and variance of offspring per generation, as well as the number of potential parents that can pass on genetic material to the next generation.

##### Neutral genetic diversity

The field of conservation genetics often aims at preserving genetic diversity to prevent inbreeding and promote evolutionary adaptation—however this is practically achieved through presumably-neutral genetic diversity as this represents the bulk of genetic diversity and serves as a proxy for adaptive/deleterious variation (1). Our current theoretical and simulation projections also do not incorporate selection on adaptive and deleterious mutations.

In addition, our framework also assumes that population sizes are large enough to be modeled across time. It is likely that once we consider extremely small populations, which has typically been the focus on conservation genetics geared towards endangered species, processes like mutational meltdown or inbreeding effects may take hold sooner than equilibrium is achieved. This is because smaller populations may suffer from increasing mutation load which in turn, causes the step-wise successive loss of individuals with high fitness due to both mutation accumulation and genetic drift (termed Muller’s ratchet).

More work needs to be done to understand how genetic diversity projections will change with selection and adaptation under habitat loss scenarios and how these processes may impact modeling for species with small population sizes. There are many possibilities in modeling these and are thus beyond the scope of this manuscript, where our focus is understanding neutral genetic diversity loss in spatiotemporal non-equilibrium dynamics.

#### Text S2 Genetic diversity dynamics for geographic habitats of 1D and 2D

In most of the main text, we only considered two-dimensional square landscapes. Yet, alternative habitat geometries are common across species. Utilizing our theoretical framework, we explore the implications of genetic diversity across 1-D habitats and compare it to the 2-D habitat that we study in the main text.

We find that short-term genetic diversity loss for 1-D habitats is more severe than 2-D habitats with increasing habitat area loss **(Fig. S25)**. This relationship is exacerbated when the migration rate is low **(Fig. S25)** and expectedly, becomes more dramatic with increasing habitat loss. This means that species that have low migration and a 1-D habitat range, or a habitat range that can be characterized as long and narrow, lose much more genetic diversity during habitat loss. This loss can be up to 3 orders of magnitude worse than that of species that also have low migration but a 2-D habitat range, or a habitat range that can be characterized as both long and wide. In the long-term, genetic diversity loss for 1-D habitats is also more severe than 2-D habitats with increasing habitat area but this behavior is more exaggerated under low migration regimes **(Fig. S25)**. Under high migration regimes, habitat geometry matters less and genetic diversity loss for both 1-D and 2-D habitats are similar **(Fig. S25)**.

To explain the differences between the 1-D and 2-D results, we consider a decomposition of species-wide π as a combination of the average π within each deme (π_within_), and the variance of allele frequencies across demes (d_between_):

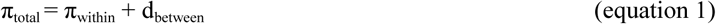

The first term measures diversity within each deme and refers to the average genetic diversity observed within individual subpopulations while the second term measures divergence across demes and measures how different allele frequencies are from one deme to another. This second term will be larger the less migration there is between pairs of demes.

Habitat loss affects each of these two terms differently. Intuitively, π_within_ can be thought of as being related to the effective number of ancestors a randomly chosen individual has. If an individual is in a deme that has many neighbors that are connected by high migration rates, then that individual could have ancestors from any of these nearby demes, resulting in a higher diversity within that deme. Conversely, if an individual is from an isolated deme, their ancestors must all come from just that deme, resulting in lower within-deme diversity. As a result, higher migration rates result in larger within-deme diversity. On the other hand, d_between_ can be thought of as how difficult it is to migrate from one deme to another on average. The more connected demes are by migration, the more similar their allele frequencies will be and the smaller d_between_ will be. As a result increasing migration results in smaller d_between_.

The geometry of the habitat affects how habitat loss changes π_within_ and d_between_. For example, in a 1-D habitat, losing 90% of the habitable area results in the remaining demes being much closer together, resulting in a much smaller d_between_. In contrast, in a 2-D habitat, even after losing 90% of the habitable area, some pairs of demes remain extremely distant and poorly connected by migration, having only a modest impact on d_between_. As a result, when migration rates are low enough to make d_between_ the dominant contributor to π, habitat loss affects π much more strongly in a 1-D habitat than a 2-D habitat.

#### Text S3 A coalescent interpretation of habitat loss

These incredibly large long-term genetic diversity losses are worrisome, but have an intuitive population genetic explanation **(Fig. S13)**. Diversity in population genetics is related to the number of potential ancestors an individual might have (2–4). If a population of present day individuals has a large pool of potential ancestors, then the population is more diverse than a population with a smaller pool of potential ancestors. Immediately following habitat loss, individuals can still have ancestors from across the entire species range, and hence habitat loss has little immediate effect on π. As time goes on, however, all of an individual’s ancestors that lived after the loss of habitat must come from a smaller pool of ancestors living in the habitable area. After enough time has passed, all of the individuals in a population will have a most-recent common ancestor that survived during the habitat loss, and therefore all of their relevant ancestors will have come from the smaller pool of individuals that could live in the reduced range. At this point, π equilibrates to its new value and no further changes should occur.

#### Text S4 Genetic diversity-area relationships and power laws

##### Background on biodiversity SAR and population genetics diversity MAR power laws

Classic population genetic theory has described that as individuals of a species move over generations and accumulate mutations, an isolation by distance pattern emerges (5), whereby the genetic distance between two individuals increases with the geographic distance between them.

From this principle, it is expected that larger areas of populations should harbor larger numbers of mutations. It was not until recently that we formally proposed and described a mutations-area relationship (6) inspired by the well-known ecological relationship species-area relationship (7–9). Apart from the isolation by distance pattern, a suspicion that the mutations-area relationship could exist specifically as a power law equation came from the observations that species relative abundances in an ecosystem are very uneven, with the majority of species being at low frequencies called the “commonness of rarity”. This is identical to the concept of the site or mutation frequency spectrum, whereby the majority of mutations remain at low frequency. It has also been pointed out that a number of ecological and evolutionary principles are analogous: speciation is equivalent to mutation, ecological drift is equivalent to genetic drift, environmental species filtering is equivalent to natural selection (10). From the assumption of “commonness of rarity” of species abundances following a log-Normal distribution, Preston analytically derived a power law of species-area relationship with the rationale behind that as area of ecosystems or islands increase one would encounter more new, rare, species.

The species-area relationship (SAR) power law then follows: *S=cA^z^* ; where species is *S* and area is *A*, and the scaling coefficient *z* describes the spatial structure of species in geographic space with a constant *c*. Preston derived theoretically the scaling should be z=0.27, under a number of assumptions (Fig. S3). This has been empirically shown to be close to reality (11), although there is some variation across ecosystems and spatial scales with causes and implications widely discussed although still debated (8).

The equivalent of species numbers of richness in genetics is the allelic richness, or segregating sites, or plainly number of mutations. We hence proposed a mutations-area relationship (MAR) of the same form: *M=cA^z^* ; with *z_MAR_* (to distinguish from the *z_SAR_*). The first empirical tests of such relationships over 10 thousand genomes of 20 plant and animal species showed an intriguing average scaling per species of *z_MAR_* = 0.3 (6).

We are yet to analytically derive an exact expectation of a *z* value given species traits, but the boundaries of the *z_MAR_* ⊂ (0 - 1] parameters are straightforward . From classic population genetics, we can derive the expectation under no structure, a panmictic population grows its number of segregation sites proportional to the number of individuals as: *M ∼ log(N) ∼ N ^z→0^.* In the opposite scenario, a species whose populations accumulate totally independent new mutations, under an infinite genomic sites assumption, would grow its number of segregating sites exactly proportional to area: *M ∼ A^1^*.

##### Power law to predict diversity extinction fractions with area

An appeal of defining within-species genetic variation with a MAR power law is for the widespread use of SAR power law for conservation of species richness (12) including Intergovernmental Panel for Biodiversity and Ecosystem Services (IPBES) (13). Habitat area loss is the number one threat of species to extinction and much conservation management is area based. For instance, IPBES reports that across countries we have already destroyed or altered ∼50% of Earth’s terrestrial habitats. The rationale of the power law approach for predictions is that under the original intact conditions (past) the diversity (alleles for MAR, species for SAR) is given by the past area: *cA^z^* ; and with a contemporary altered landscape with a reduction (*a*) in area: *A_present_ = A_past_ - a*; the new genetic diversity would be: *cA^z^ .* Taking the ratio removes the constant *c* and gives us the proportion of diversity remaining: (*A_present_ / A_past_*)*^z^*. If we just have the area lost: *A_loss_ = 1-(A_present_ / A_past_)*; which is how often conservation organisms report threat on ecosystems or species, we can just rearrange: *(1-A_loss_)^z^* ; and if we want to express the diversity loss instead of diversity remaining fraction, we can use: *1-(1-A_loss_)^z^ .* All these slightly rearranged versions of the MAR/SAR equations are equivalent and are very easily deployed by conservation practitioners.

The MAR equation captures spatial genetic structure of species and thus predicts much more substantial loses of genetic diversity compared to classic population genetic expectations of population bottlenecks because in population genetics most often we assume population panmixia (i.e. free gene flow and no population structure). We can come up with an equation on the immediate effect of a bottleneck. Before we show that with no structure, *M ∼ log(N).* Similarly as MAR, the loss of genetic diversity after a population reduction of *N_x_* individuals would be:

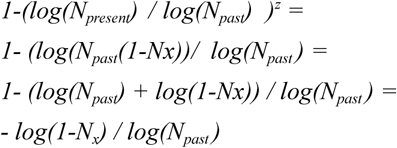

This derivation shows that the loss of allelic richness or mutations is in the scale of *log(1-x)*; which is very slow, as we expected from having derived the trend under panmixia *z_MAR_ ≈ 0*.

In the most extreme spatial structure scenario, a MAR with parameter *z_MAR_ ≈ 1*, predicts the fraction loss of geographic area directly translates to the same fraction loss of genetic diversity. Luckily most species studied have a moderate *z_MAR_* with an average of 0.3 (**Table S20**).

##### MAR in the long-term

Our MAR framework was developed for short-term genetic diversity loss. That is, if an immediate area reduction of a geographic range of a species happens, how many unique genetic variants were found in that area and are thus gone. However, with time, the original area reduction causes an increased stochasticity within a species population dynamics that incurs in further loss of genetic variants from increased genetic drift.

We could not derive analytically the *z_MAR_* of long-term genetic diversity but the current manuscript aimed to address this through SLiM simulations. Comparing increasing area loss % we impose on a simulated species in the landscape with the long-term genetic diversity after running the simulations for hundreds to thousands of generations, we could phenomenologically fit a power law (**Fig S15)** which discovered a worrisome long-term *z_MAR_ ∼ 1*.

##### From mutations-area relationship (MAR) to genetic diversity-area relationship (GDAR)

We proposed MAR to model allelic richness with area, analogous to species richness in SAR. There are of course multiple metrics to measure genetic diversity. The most prevalent one together with allelic richness in nucleotide genetic diversity or average pairwise distance (π), defined as: 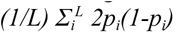; where *L* is the total number of genetic variants assessed and *p_i_* is their frequency in the population.

From a theoretical standpoint, it is unclear why nucleotide diversity π would follow a power law with area.

In fact, the appeal of π for population genetics is that because its value is more affected by frequency than the number of loci, it is a more robust metric to the number of sampled individuals in a population (and thus also robust to area sampled).

This is also the case for species diversity metrics such as Shannon’s diversity or Simpsons’ diversity indices. In fact, Simpson’s index or evenness: 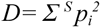; for *S* species in the ecosystem and *p*_i_ for their relative frequency, which is remarkably similar to the inverse of average genetic diversity π. Despite the biogeography and community ecology field has empirically fitted power laws to other diversity metrics such as Simpson’s species diversity (8, 14). Although not theoretically derived, we therefore nevertheless try to fit different area relationships to π, as this would provide a powerful empirical equation for conservation.

#### Text S5 Initial results on habitat restoration and assisted population recovery

Motivated by the plethora of restoration projects occurring globally, we wanted to understand and explore recovery dynamics of genetic diversity after habitat loss. We caveat these results by restating that our theoretical and simulation-based predictions only consider neutral evolution.

Utilizing our existing habitat loss from one leading edge and habitat loss with fragmentation scenarios, we “restored” habitats by restoring carrying capacity of habitats with carrying capacity zero to one **(Fig. S23)**. For example, for a 50% habitat loss, this means that 50 grids within the 10×10 simulation map have a carrying capacity of 0. During restoration, these 50 grids now have a carrying capacity 1. Thereby, allowing individuals in nearby habitable areas to disperse and eventually repopulate these newly rehabitable areas. Here, we wanted to examine if time of restoration mattered and performed habitat restoration at varying numbers of generations after habitat loss (termed 2000 (early), 10000 (medium) and 20000 (late) generations). We then tracked species-wide genetic diversity metrics over time and across 10, 50, 90% habitat loss to understand if there was a potential tipping point in which genetic diversity cannot be restored **(Fig. S24)**.

Overall, we found that genetic diversity eventually always bounces back to 100% given sufficient time. This behavior is more pronounced in the habitat loss from one leading edge scenario, where a substantial short-term reduction in genetic diversity is reported at 50% habitat loss before a gradual increase in genetic diversity metrics with time at the point of restoration **(Fig. S24)**.

Given the complicated genetic diversity trajectories of habitat fragmentation scenarios, we observed less reduction in genetic diversity at 50% habitat loss and a slight increase in genetic diversity metrics at the point of restoration **(Fig. S24).** These observations are similar even when we look at within-species metrics **(Fig. S25)**.

Overall, these simulations seem to show an overall optimistic conclusion that as long as habitat restoration occurs, genetic diversity will eventually be restored. However, given that these long-term trajectories span tens of thousands of generations, these optimistic conclusions might prove to be unrealistic, and are likely species specific. It is likely that species with faster generation times will reach long-term genetic diversity loss quicker than species with longer generation times but genetic diversity loss for species with higher generation times may be more easily recoverable, given that genetic diversity restoration is slow. For species with long generation times, it may be impossible to recover their genetic diversity within timescales relevant for conservation policy. In addition, these results will likely change as one considers the addition of adaptive and deleterious mutations. Hence, more work needs to be done to develop the theory and predictions that would follow habitat destruction and corresponding habitat restoration projects.

## Supplemental Figures

**Fig. S1.**
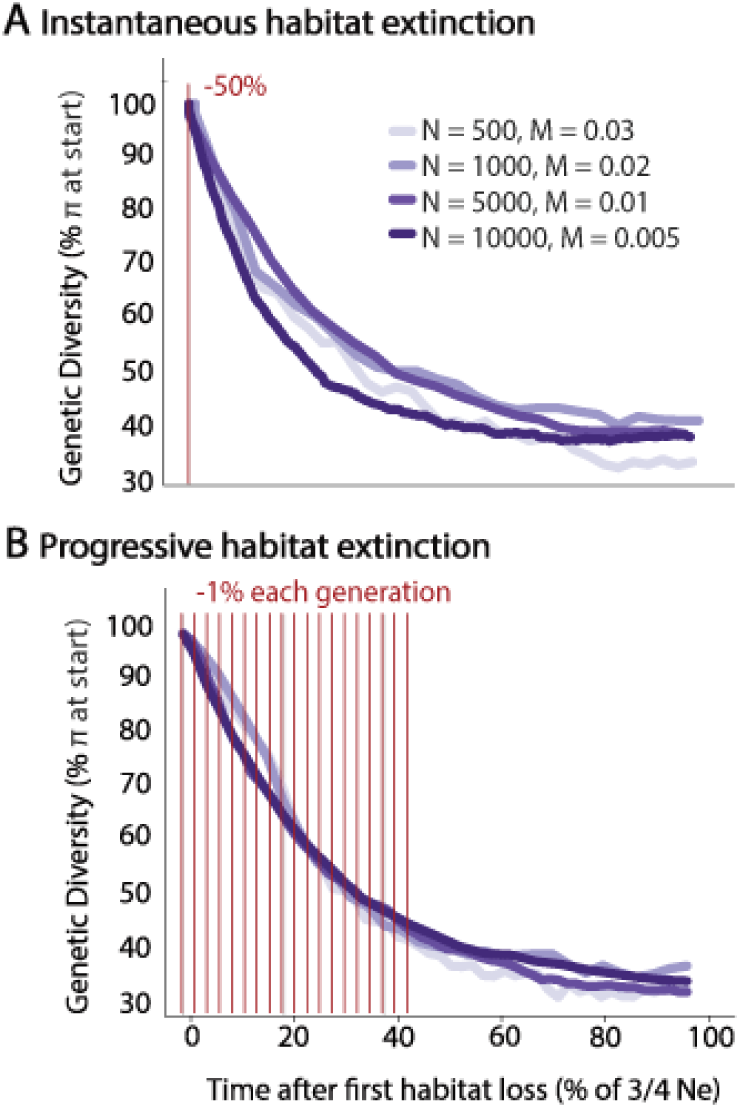
Temporal dynamics of genetic diversity. (A) Trajectories of genetic diversity loss after 50% instantaneous extinction of habitat has halted. Parameters that alter these dynamics include population size (N) and migration rate (M). (B) Trajectories of genetic diversity loss with gradual extinction of 50% habitat. Gradual habitat loss was kept at 1% of habitat loss per ∼11 generations. Colors represent different values of population size and migration rate used, and are consistent between A and B.

**Fig. S2.**
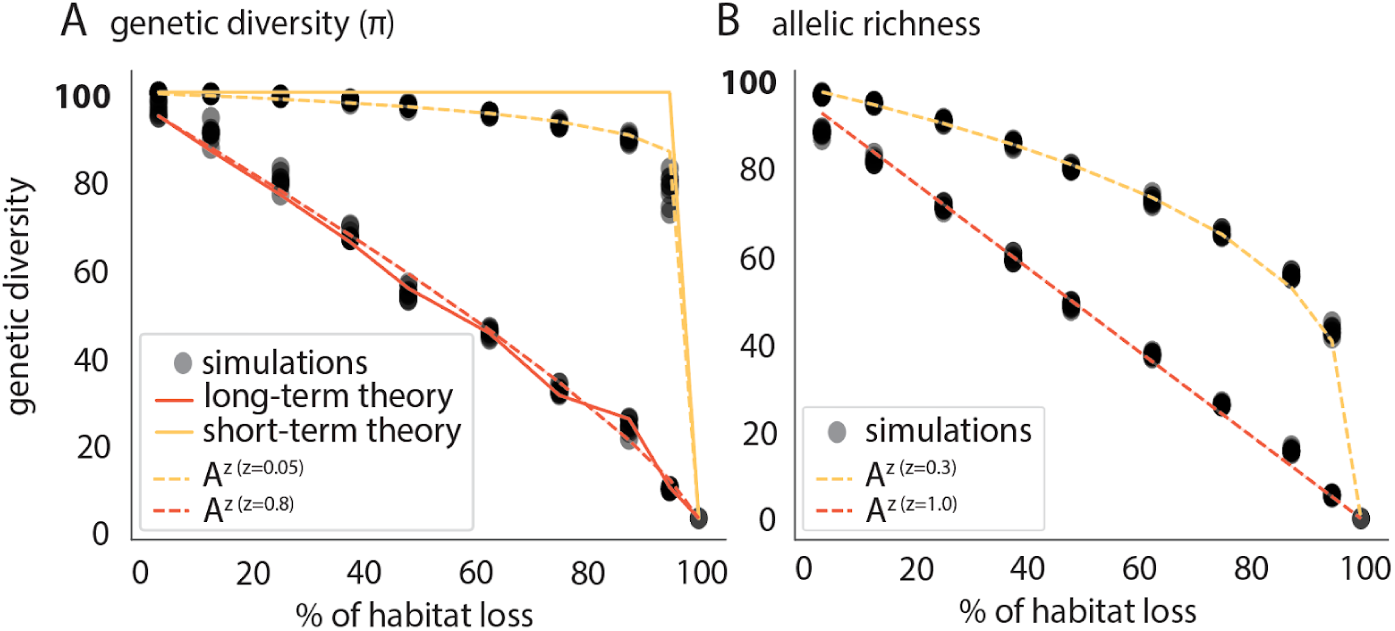
Relationship between genetic diversity and habitable area. In log space, genetic diversity and habitable area follow a power law relationship, genetic diversity = A^z^ for both short and long-term trajectories of **(A)** π and **(B)** allelic richness. Solid lines illustrate the genetic diversity trajectories seen using our theoretical and simulation-based framework. Dotted lines indicate the power law relationship using the parameters corresponding to both short and long-term respectively. Black dots represent genetic diversity trajectories using our simulations. In red are short-term estimates while in orange are long-term estimates. **(A)** In the short-term, z=0.05 while in the long-term, z=0.8 . **(B)** In the short-term, z=0.3 while in the long-term, z=1.0.

**Fig. S3.**
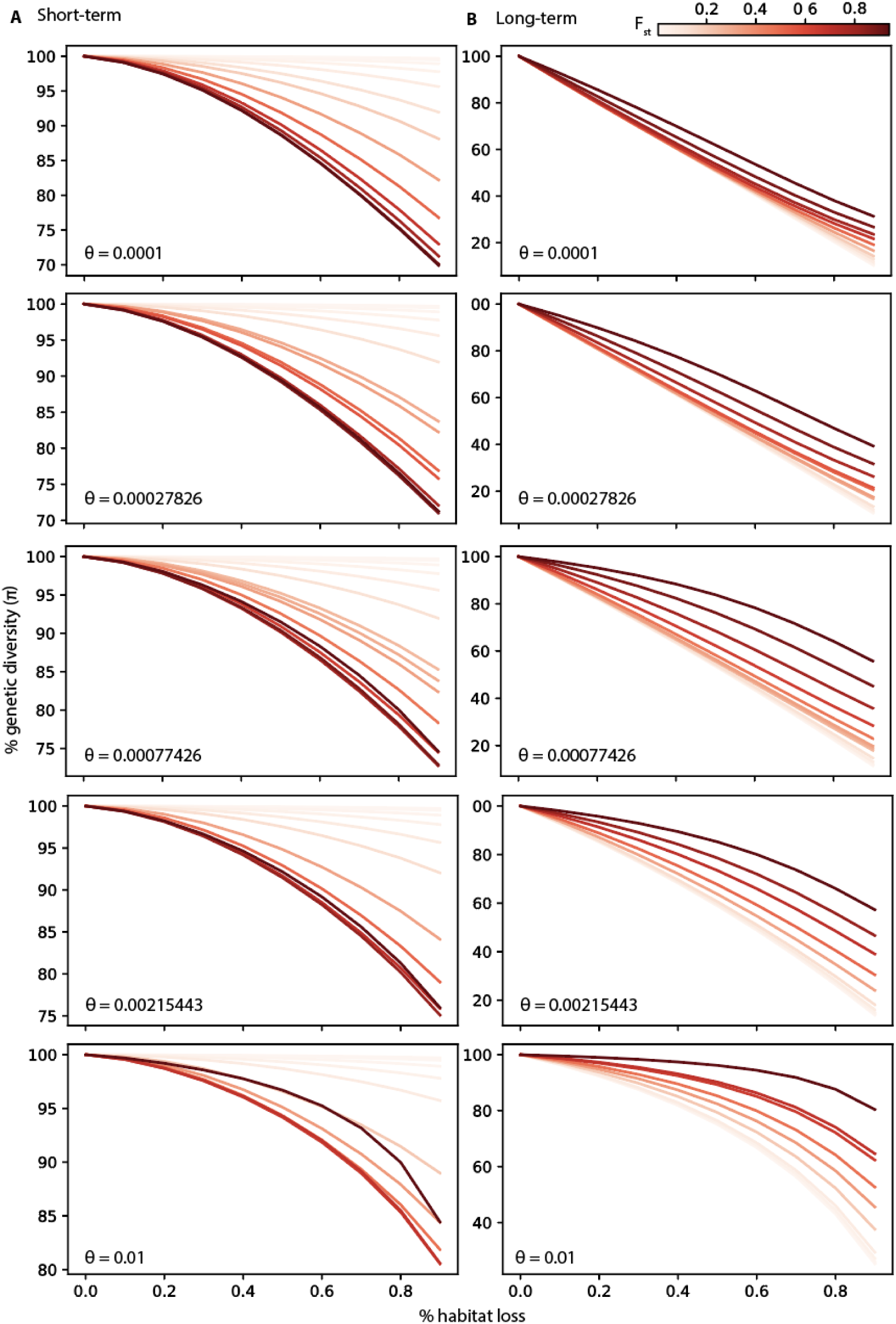
Relationship between genetic diversity and habitable area across F_ST_ and θ values for edge contraction scenarios. Theoretical projections of genetic diversity across different F_ST_ and θ values using wfmoments. Different hues of red represent different F_ST_values, as indicated via the colour bar. Each row represents a different θ value tested. Each column represents genetic diversity trajectories seen in the short and long-term.

**Fig. S4.**
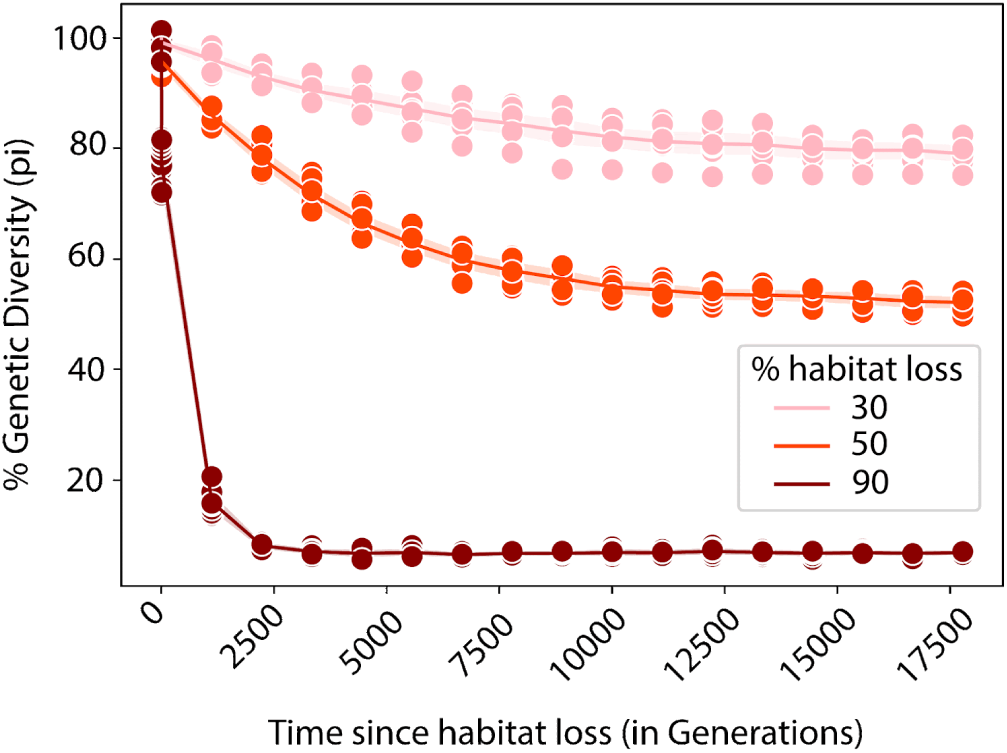
Genetic diversity extinction with species range contraction over time. Loss of genetic diversity (π) from edge range contraction (30%, 50%, 90%) over time (in Generations). Each dot represents an estimate of genetic diversity for that specific % of habitat loss at that specific time point. A total of 9 replicates were run for 30, 50, 90% habitat loss at every specific time point.

**Fig. S5.**
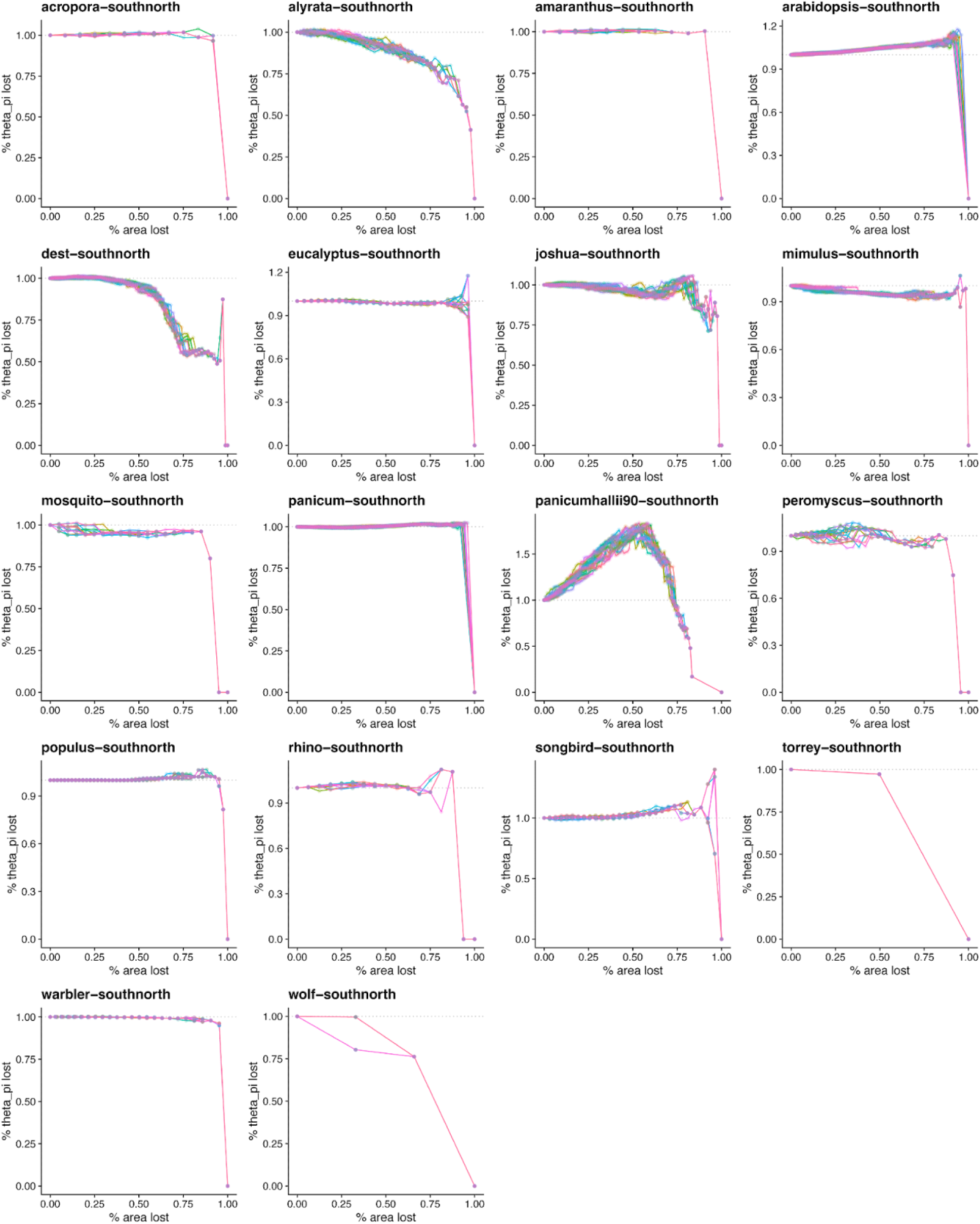
Genetic diversity trajectories across species. For each species, we plot short-term trajectories for north-south extinction. Different colors represent different replicates. The x-axis represents the percentage of area lost and the y-axis shows the percentage of pairwise differences (π) left in a species.

**Fig. S6.**
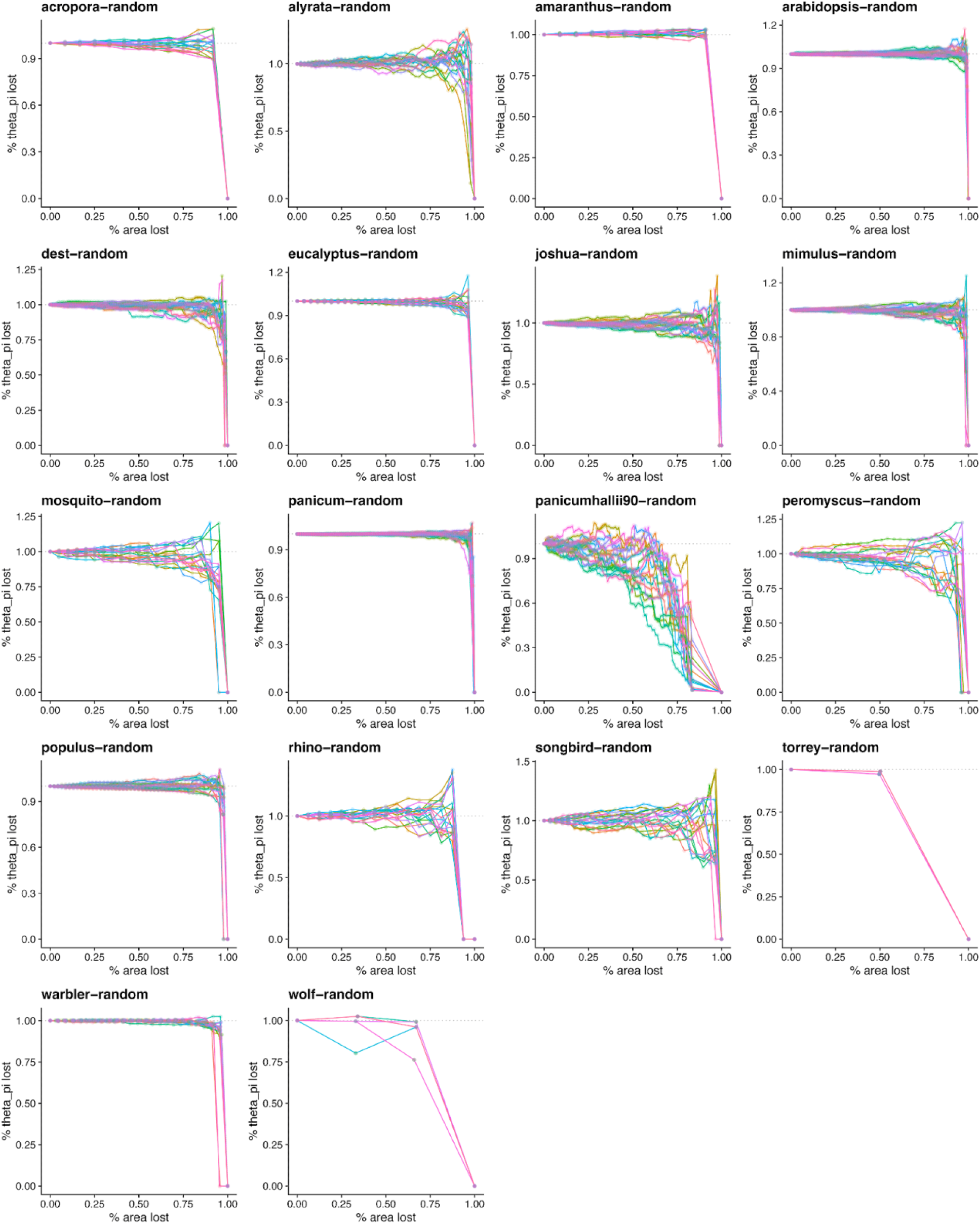
Genetic diversity trajectories across species. For each species, we plot short-term trajectories for random extinction. Different colors represent different replicates. The x-axis represents the percentage of area lost and the y-axis shows the percentage of pairwise differences (π) left in a species.

**Fig. S7.**
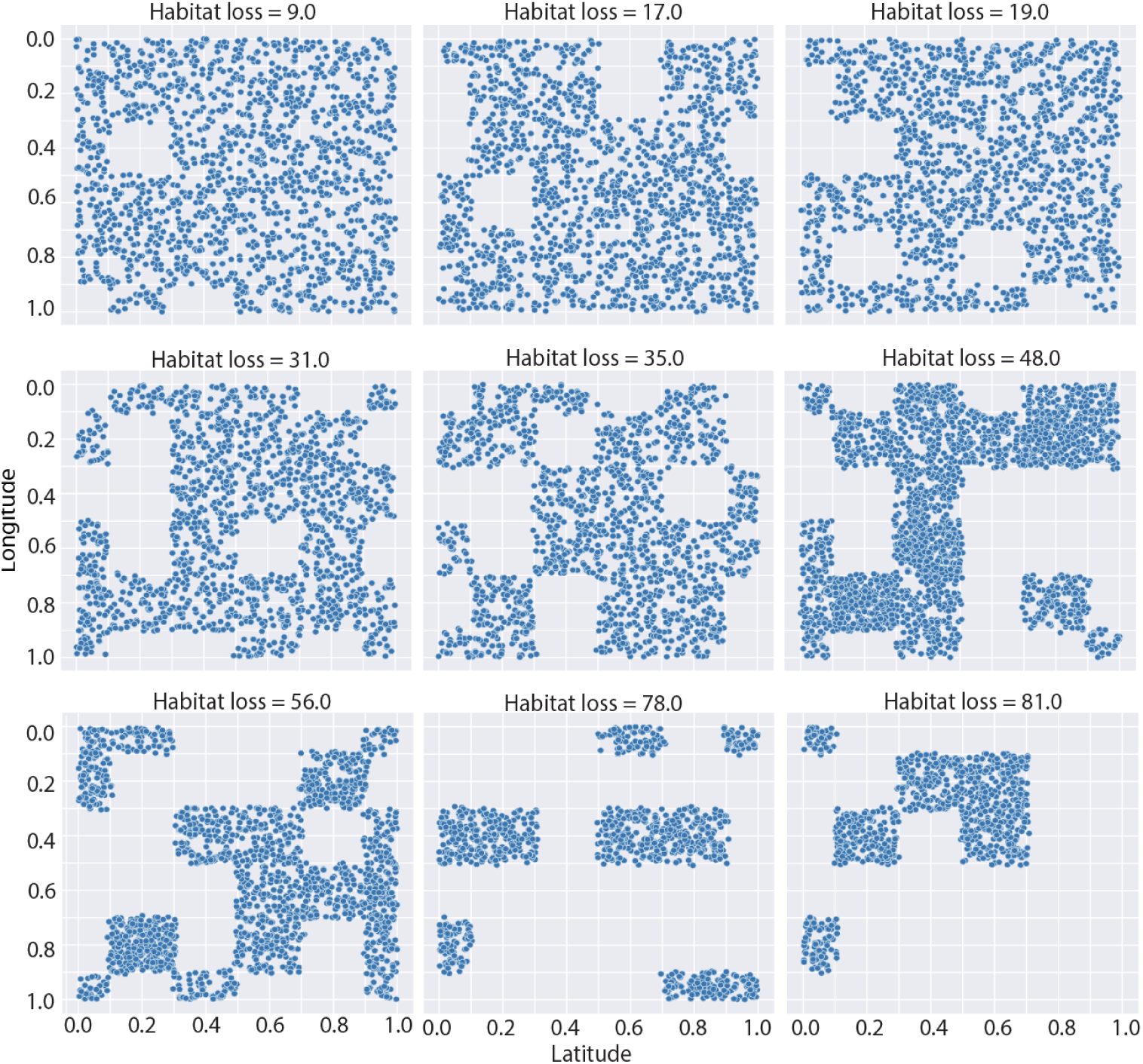
Habitat fragmentation maps. Habitat fragmentation maps shown across a range of habitat loss scenarios within the simulation. Each map is a 10×10 grid with each dot representing an individual along a 2-D coordinate system (latitude, longitude). Here we show how habitat fragmentation occurs within simulation space, with empty boxes representing the “extincted” habitat.

**Fig. S8.**
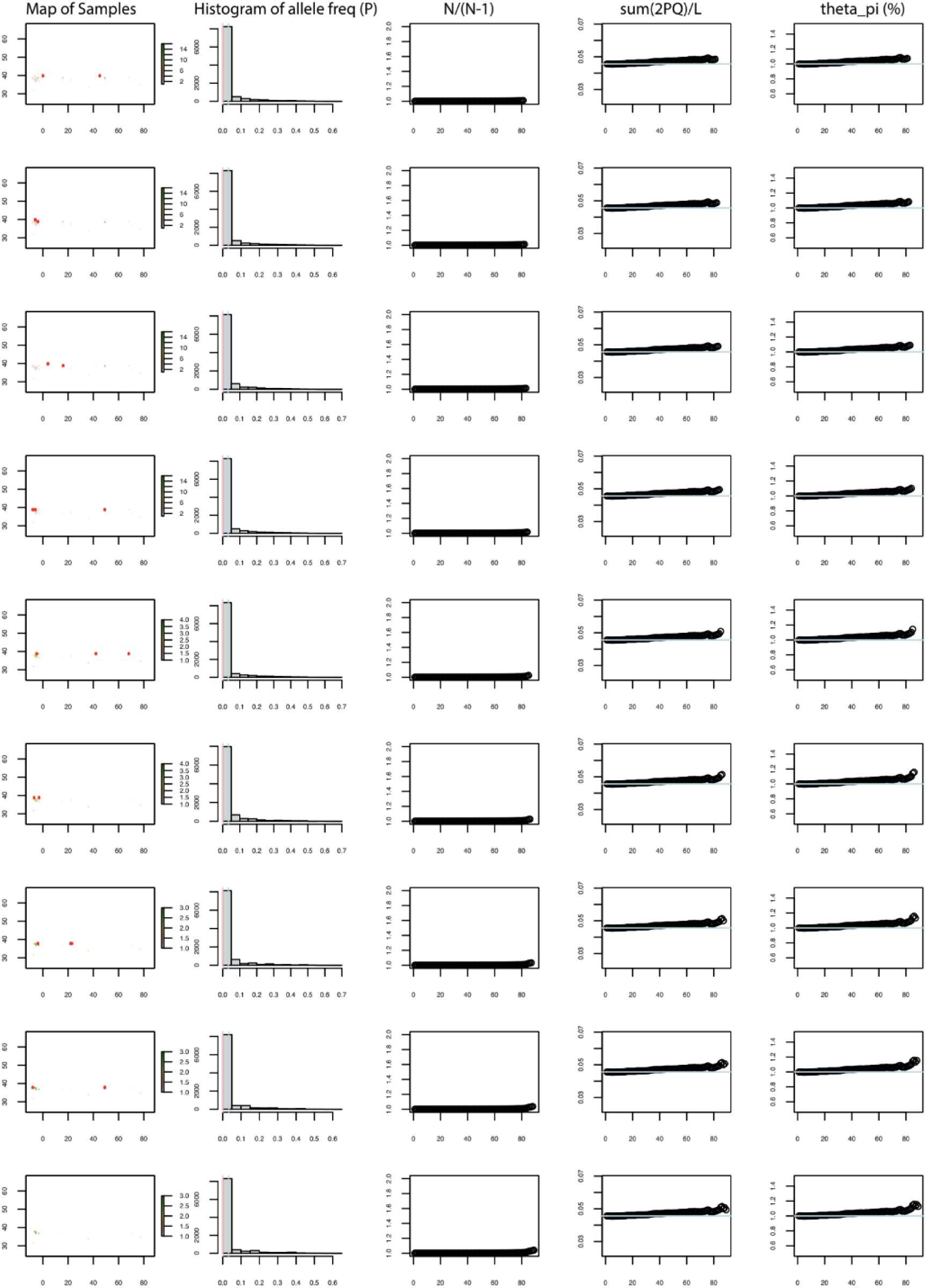
Empirical extinction simulations for *Arabidopsis thaliana*. Tracking the extinction process for Arabidopsis thaliana south-north empirical extinction simulations. Each row represents a specific time point in the simulation extinction process.

**Fig. S9.**
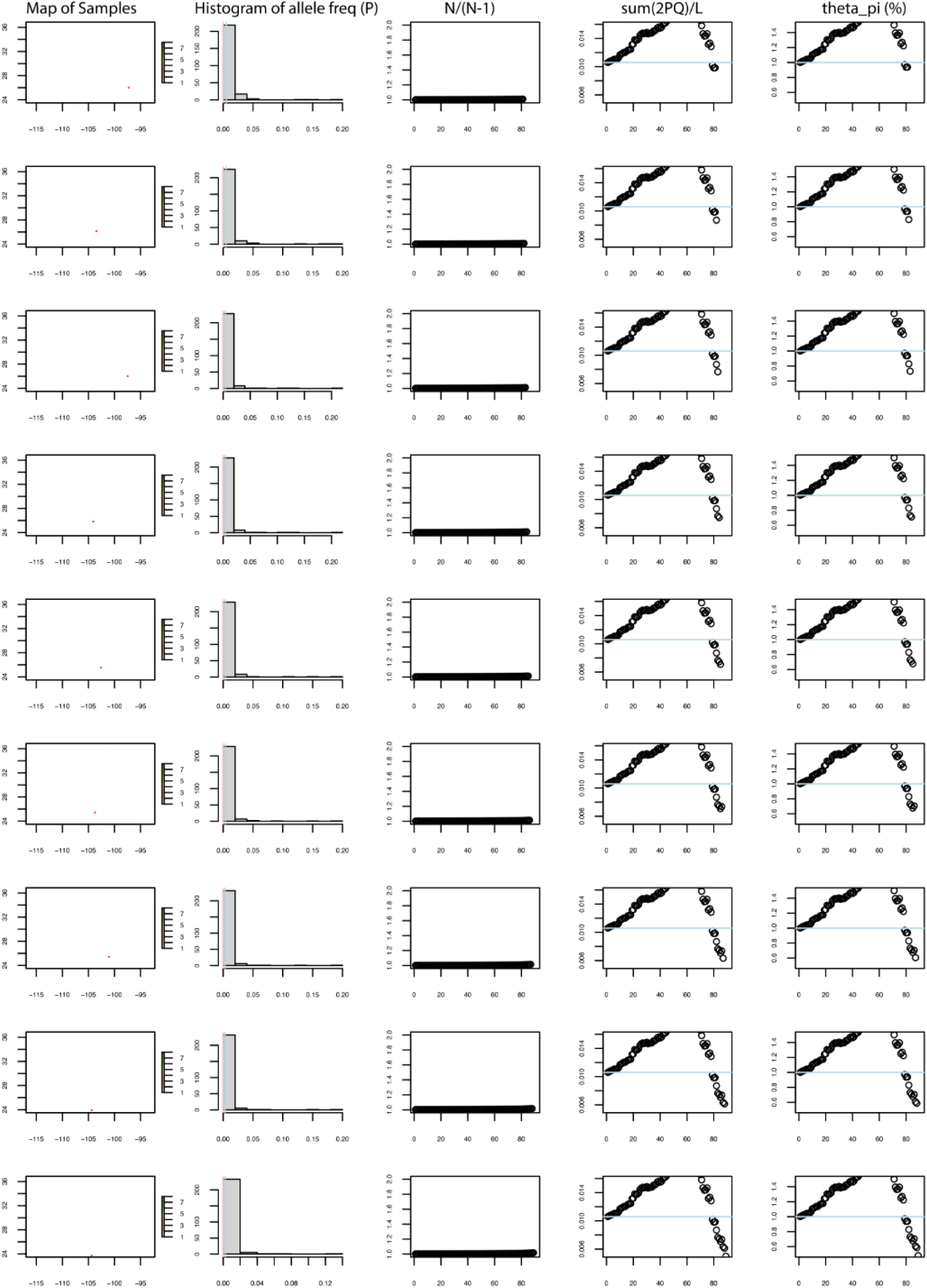
Empirical extinction simulations for *Panicum hallii*. Tracking the extinction process for *Panicum hallii* south-north empirical simulations. Each row represents a specific time point in the simulation extinction process.

**Fig. S10.**
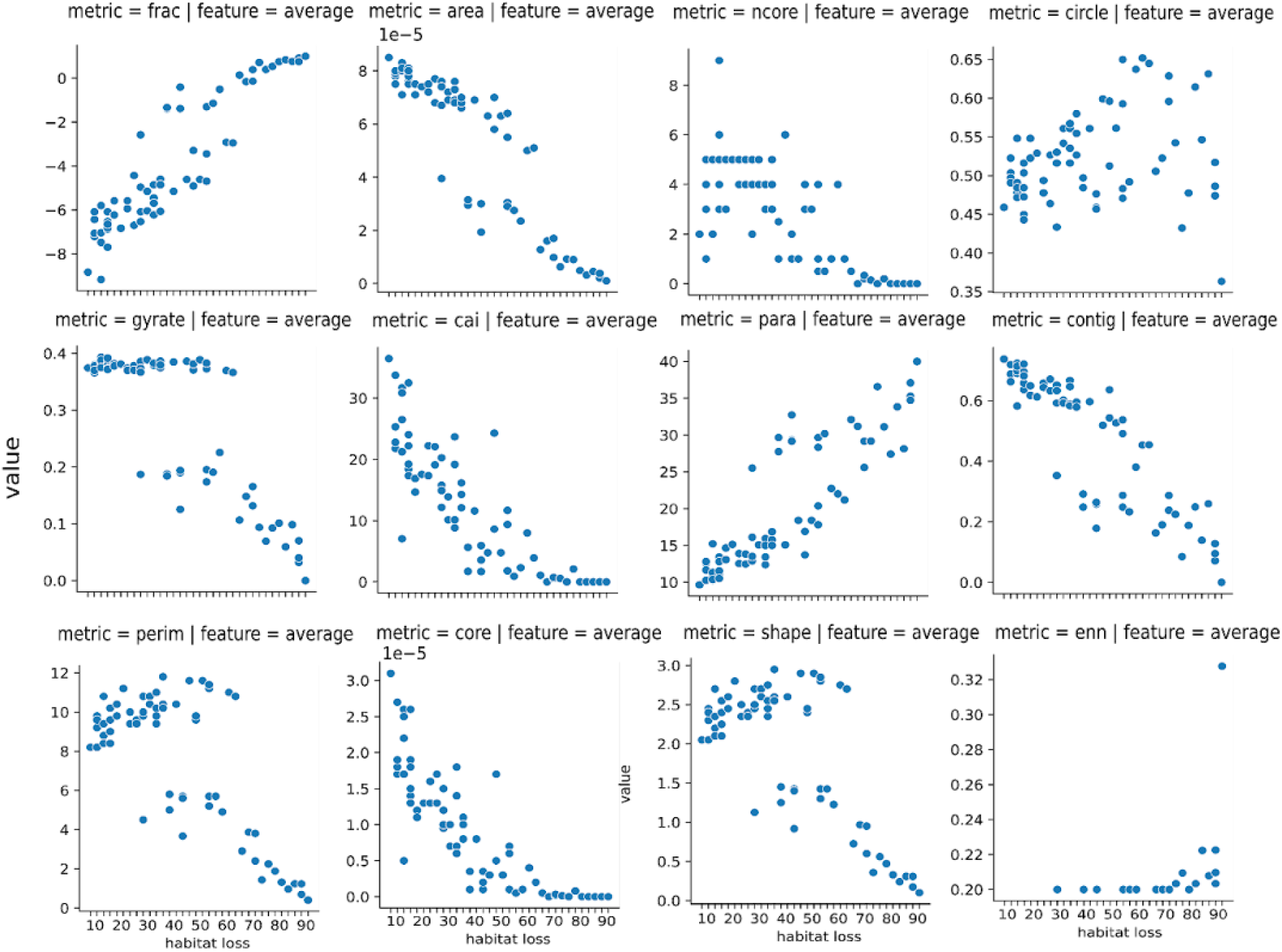
Connectivity metrics across habitat fragmentation maps (6×6) in SLiM simulations. Each plot represents a different plotted connectivity metric and the corresponding aggregation function used to report each metric. Blue dots represent different replicates for that % habitat loss. In total, 90 different simulation runs with varying habitat fragmentation maps are displayed in this figure. We considered % habitat loss ranging from 10% to 90%, at 10% increments. Metrics are selected by utilizing the landscape metrics R package.

**Fig. S11.**
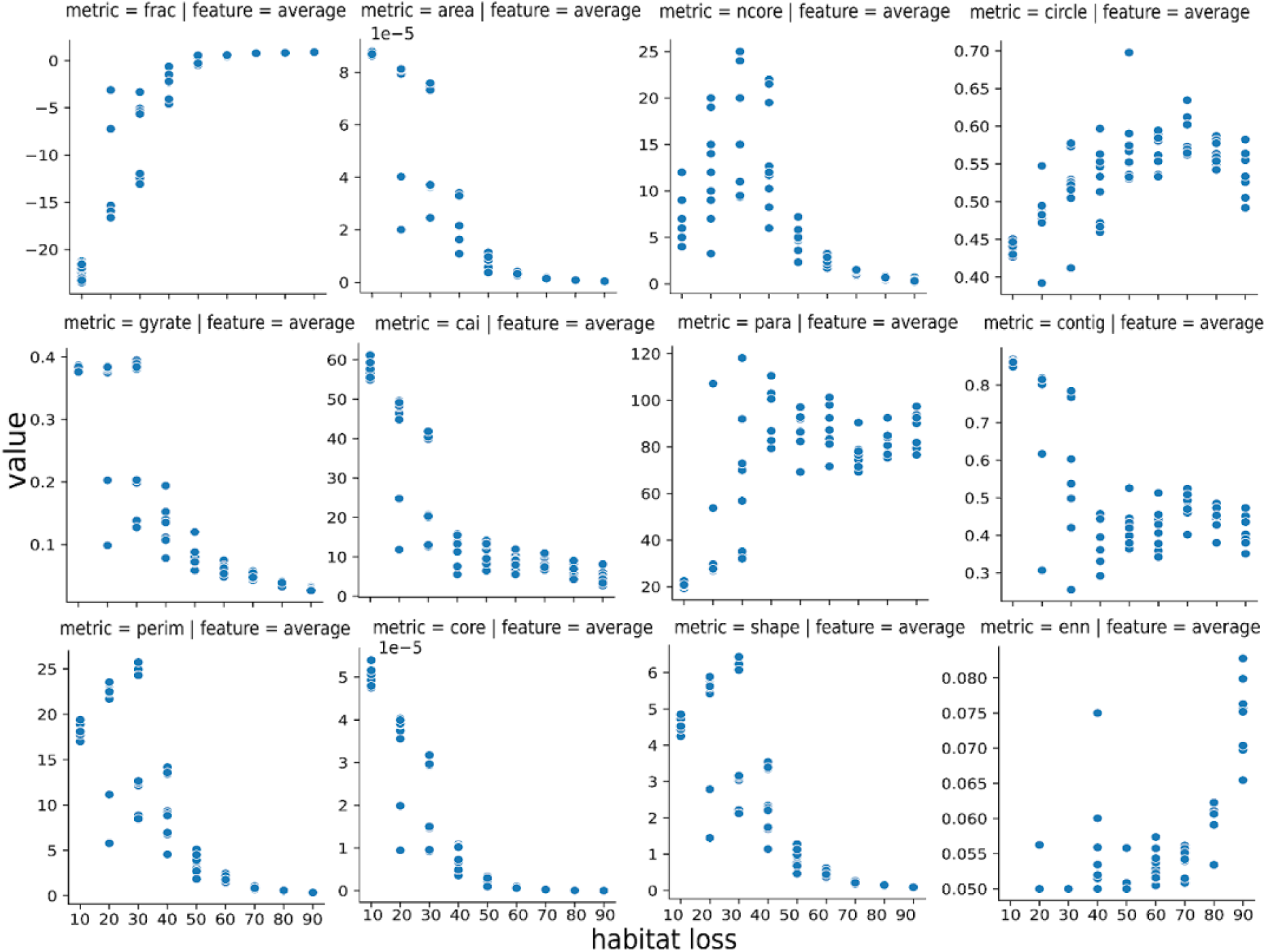
Connectivity metrics across habitat fragmentation maps (20×20) in SLiM simulations. Each plot represents a different plotted connectivity metric and the corresponding aggregation function used to report each metric. Blue dots represent different replicates for that % habitat loss. In total, 90 different simulation runs with varying habitat fragmentation maps are displayed in this figure. We considered % habitat loss ranging from 10% to 90%, at 10% increments. Metrics are selected by utilizing the landscapemetrics R package.

**Fig. S12.**
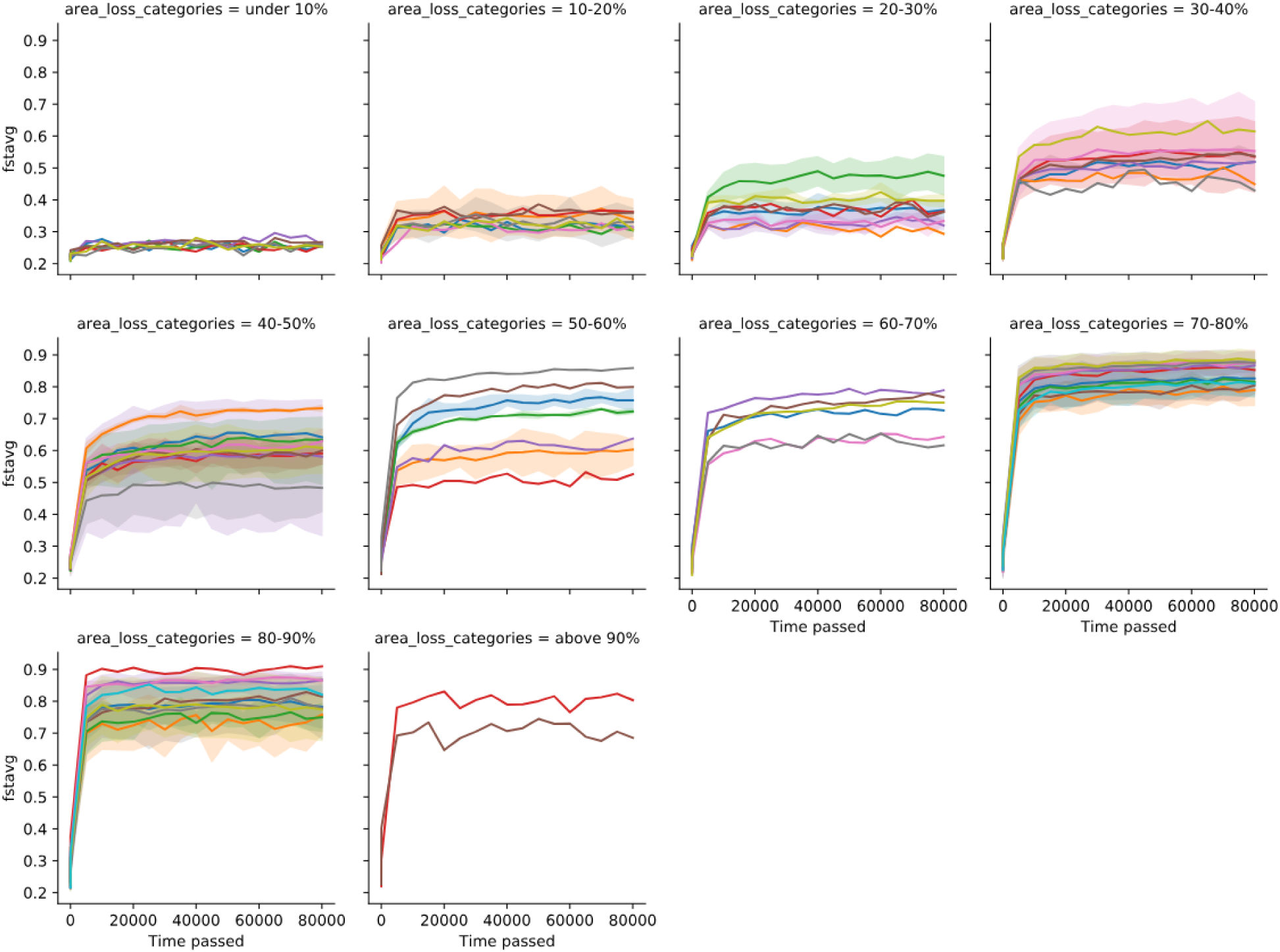
Increased F_ST_ in simulations with habitat fragmentation. We tracked individuals over 80,000 SLiM ticks after habitat fragmentation (at time=0). We tracked F_ST_ values of our populations across different area loss categories ranging from 0 to ∼90%, totalling 121 different simulation runs. Colors represent different simulation runs. Shaded area showing 95% confidence intervals for certain runs that had corresponding replicates, while others are single runs indicating just a single simulation run.

**Fig. S13.**
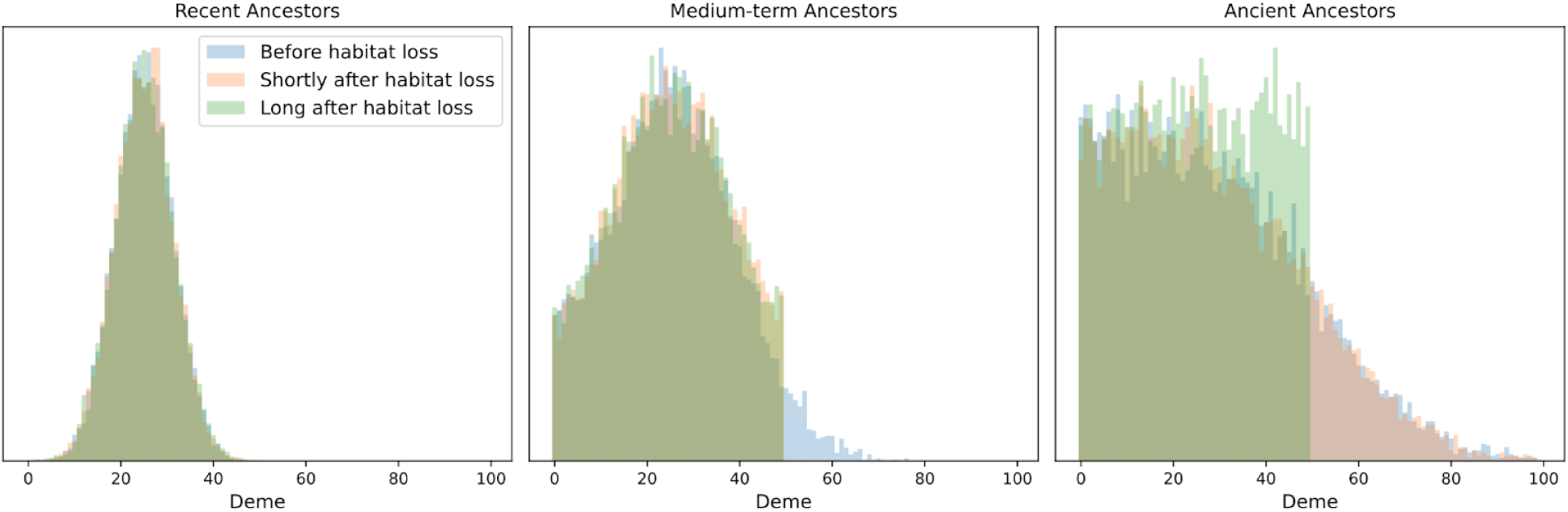
Distribution of ancestors across landscape map after habitat loss. We consider a 1D habitat of 100 demes (labeled 0, 1, … 99). All individuals are sampled in *deme 25*. At each generation, an individual has a 0.2 probability of having their ancestor come from the deme to left (if it exists) and 0.2 probability of having their ancestor come from the deme to the right (if it exists) and their ancestor comes from the same deme otherwise. “Recent ancestors” are sampled 100 generations prior to sampling time, “Medium-term ancestors” are sampled 500 generations prior to sampling time, and “Ancient ancestors” are sampled 1500 generations prior to sampling time. For the “Shortly after habitat loss” scenario, we assume that demes 50 and above became uninhabitable 500 generations ago, and for the “Long after habitat loss” scenario, we assume that demes 50 and above went extinct more than 1500 generations ago.

**Fig. S14.**
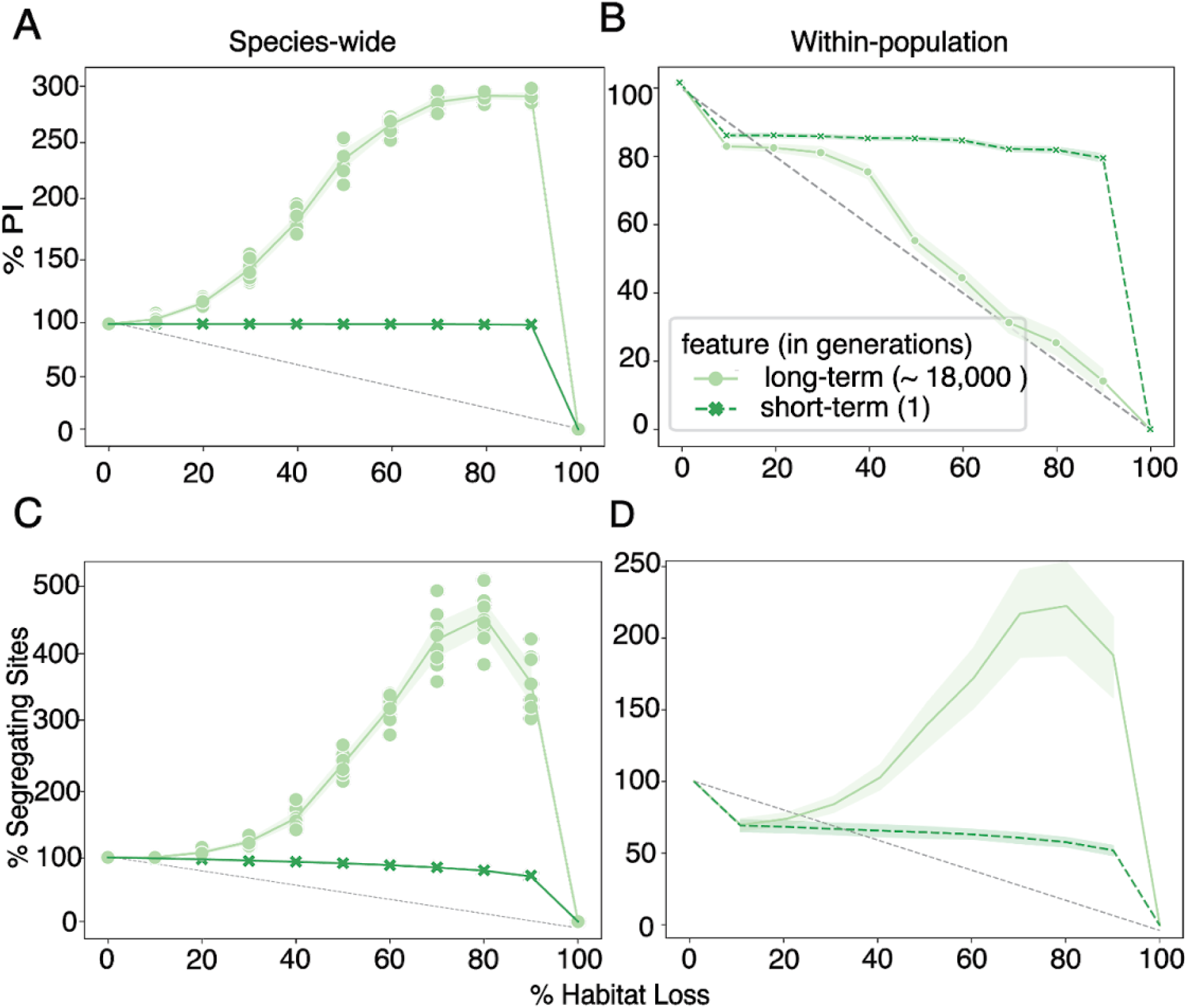
Genetic diversity metrics after habitat fragmentation. Species-wide genetic diversity, (A) π and (C) allelic richness (% segregating sites), across different percentages of habitat loss. Simulations were run for 9 replicates. Light green solid line shows average long-term genetic diversity projections using our simulation framework, with shaded area showing 95% confidence intervals. Dark green dotted line shows average short-term genetic diversity projections using our simulation framework. Each dot represents an average measure of π across a 100m-by-100m habitat map at specific percentages of habitat loss for a specific replicate, where each replicate is a different habitat fragmentation map. Light gray line indicates the y=x relationship. Within-population genetic diversity, (B) π and (D) allelic richness (% segregating sites), across different percentages of habitat loss. Light green solid line represents the average within-population long-term genetic diversity across all 5m-by-5m grids within a 100m-by-100m habitat map across all replicates, where each replicate is a different habitat fragmentation map, with shaded area showing 95% confidence intervals. Dark green dotted line represents the average within-population short-term genetic diversity across all 5m-by-5m grids within a 100m-by-100m habitat map across all replicates, where each replicate is a different habitat fragmentation map, with shaded area showing 95% confidence intervals.

**Fig. S15.**
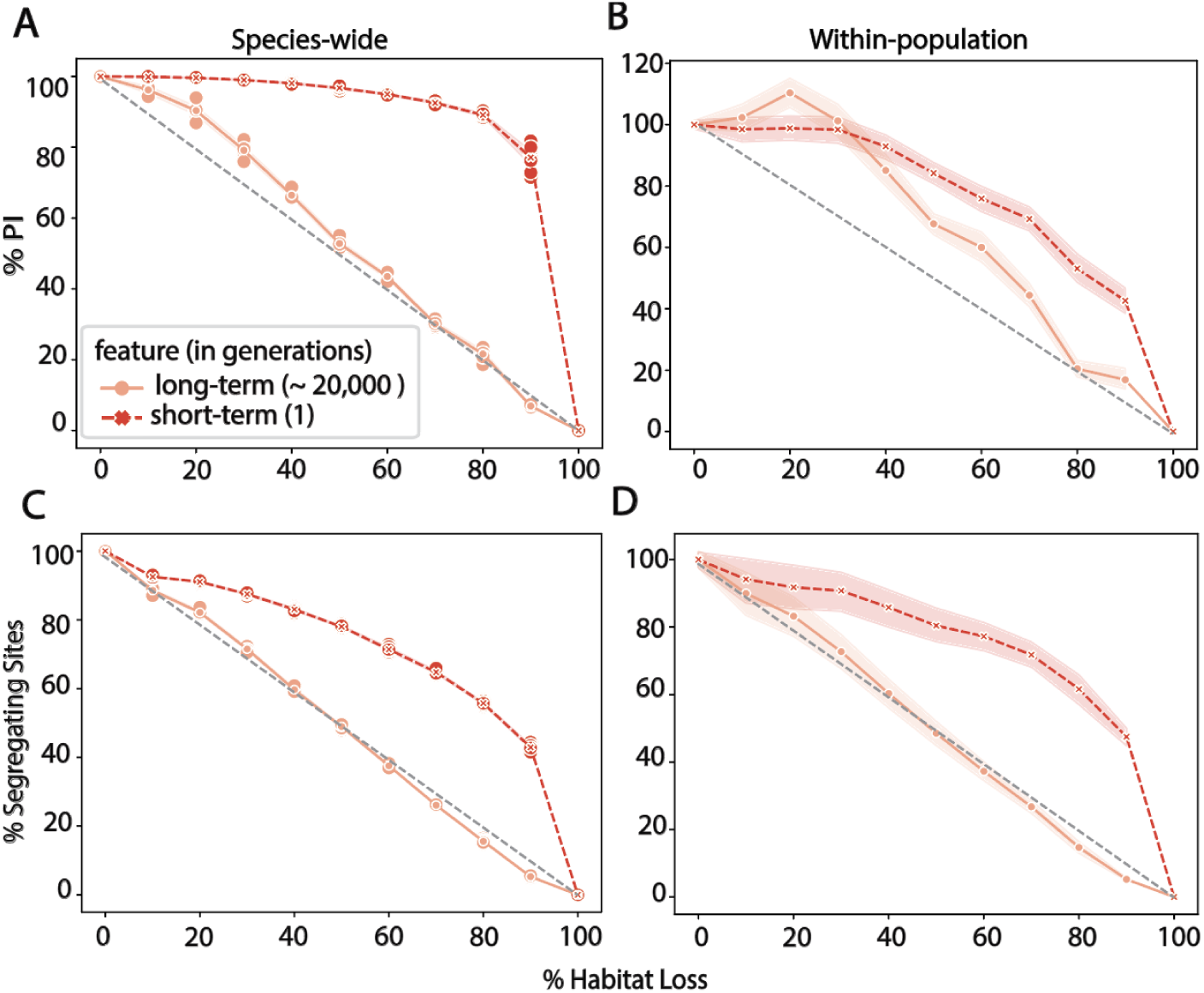
Genetic diversity metrics after habitat loss from one edge. Species-wide genetic diversity, (A) π and (C) allelic richness (% segregating sites *S*), across different percentages of habitat loss. Simulations were run for 9 replicates in 10 different area loss percentages. Light red solid line shows average long-term genetic diversity projections using our simulation framework. Dark red dotted line shows average short-term genetic diversity projections using our simulation framework. Each dot represents an average measure of π across a 100×100 units habitat map at specific percentages of habitat loss for a specific replicate. Light gray line indicates the y=x relationship. Within-population genetic diversity, (B) π and (D) allelic richness (% segregating sites), across different percentages of habitat loss. Light red solid line represents the average within-population long-term genetic diversity across all 5×5 units grids within a 100×100 units habitat map across all replicates, with shaded area showing 95% confidence intervals. Dark red dotted line represents the average within-population short-term genetic diversity across all 5×5 units grids within a 100×100 units habitat map across all replicates, with shaded area showing 95% confidence intervals.

**Fig. S16.**
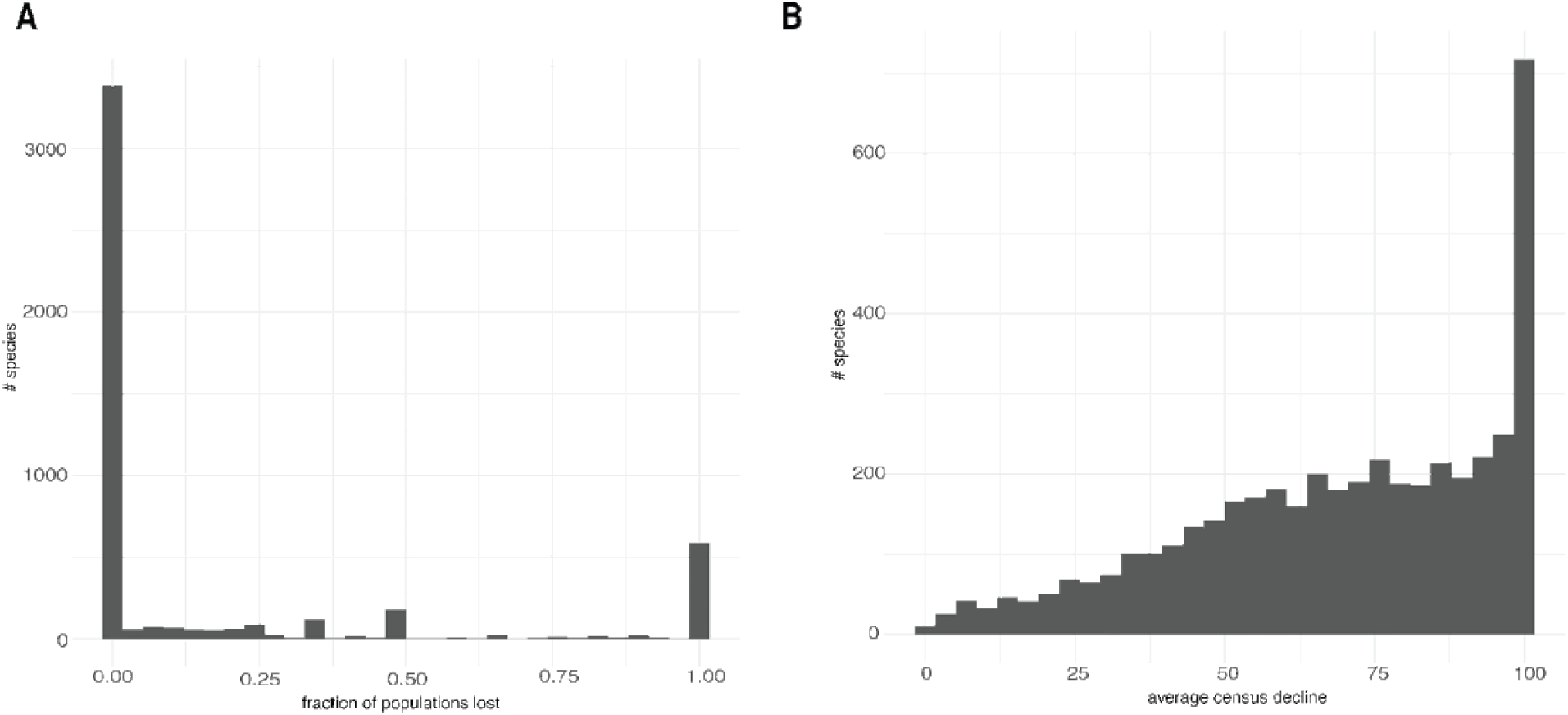
LPI raw summary data. Using data from the Living Planet Index 2024, we tracked the (A) fraction of populations lost and (B) the average census decline from 1967 to 2020 for 32895 populations across 3417 species total. Histograms show the number of species that have that fraction of population lost and average census decline respectively.

**Fig. S17.**
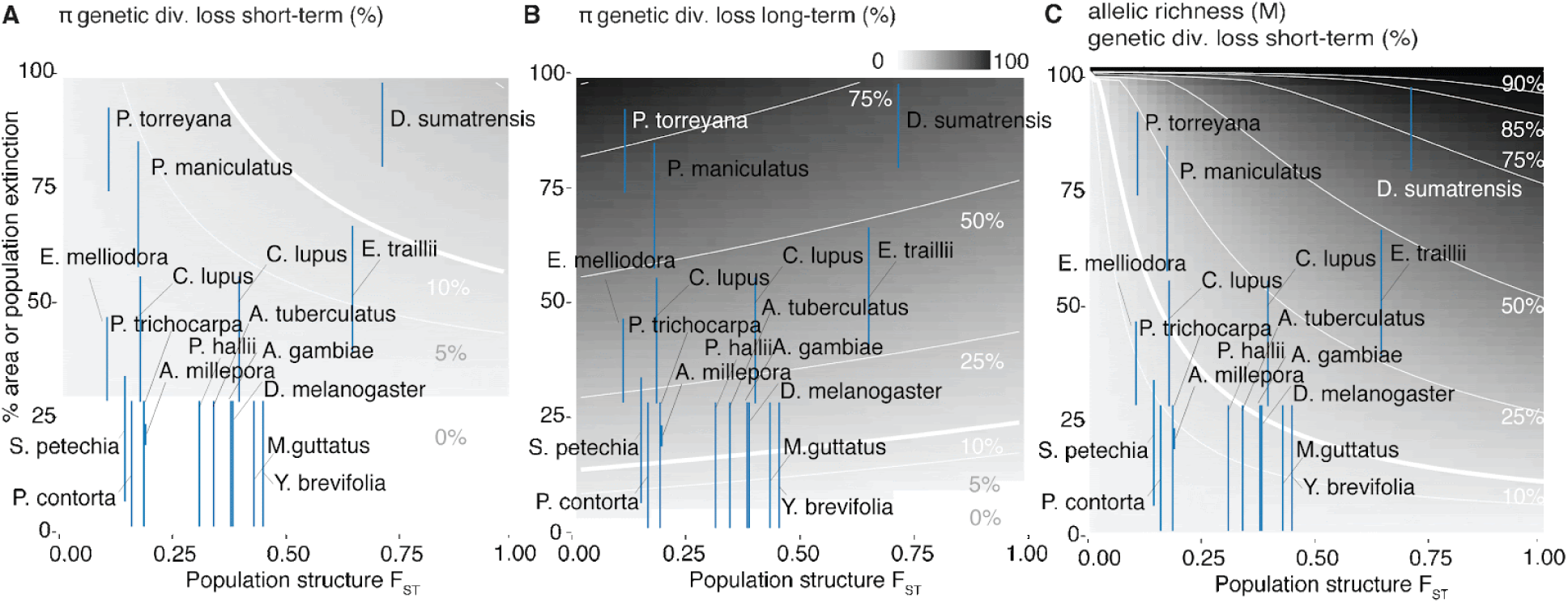
Genetic diversity loss (nucleotide diversity π and allelic richness *S*) based on MAR and GDAR. (**A**) Relationship between population structure, F_ST_ and the percentage of area or population extinction in the short-term. Estimated percentage of genetic diversity loss (π) in the short-term is represented as a gradient of white to gray, with corresponding isolines in white. Blue lines represent individual populations of species across 17 species from publicly available datasets. As a proxy for area extinction, we utilized population size changes over time for populations tracked by the Living Planet Index, when available. If that was not available, we approximated area extinctions using their Red List categories. (**B**) Relationship between population structure, F_ST_ and the percentage of area or population extinction in the long-term. Estimated percentage of genetic diversity loss (π) in the long-term is represented as a gradient of white to black, with corresponding isolines in white. Explanations of blue lines are similar to A. (**C**) Relationship between population structure, F_ST_ and the percentage of area or population extinction in the short-term. Estimated percentage of genetic diversity loss, allelic richness (M) or segregating sites (*S*) in the short-term is represented as a gradient of white to black, with corresponding isolines in white. Explanations of blue lines are similar to A.

**Fig. S18.**
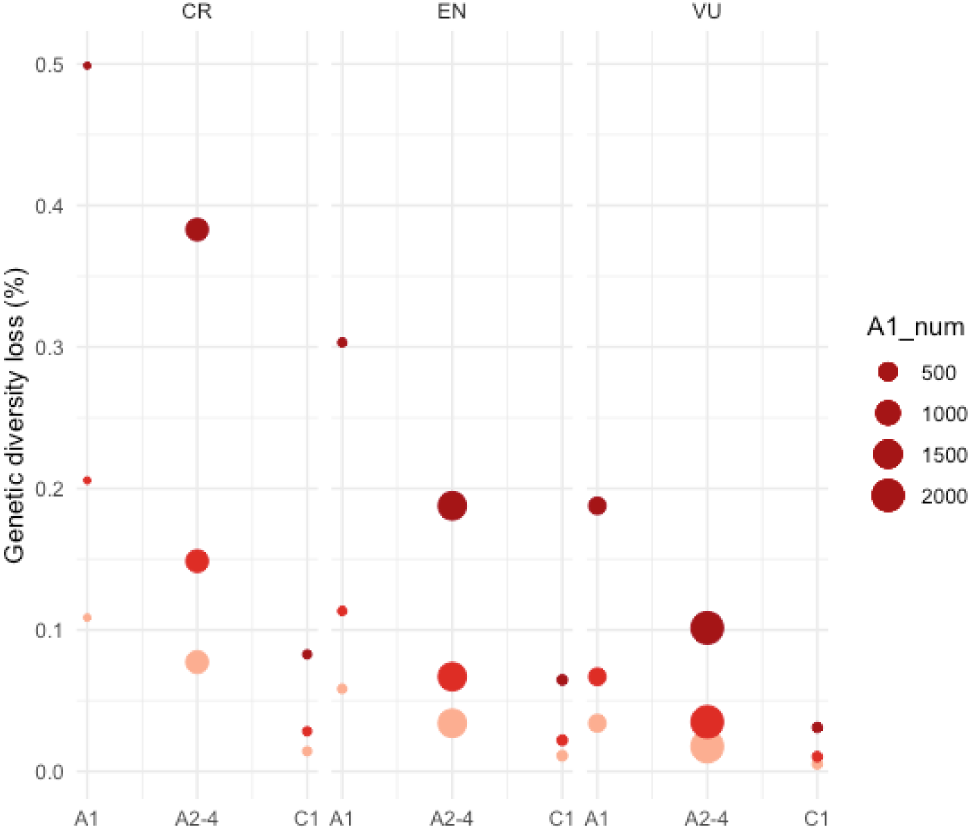
Translation of Red list genetic diversity loss (allelic richness *S*) predictions using MAR. Utilizing the IUCN Red List criterias as proxy for habitat loss, we obtained values for A1, A2-4 and C1 across three threaten statuses: Critically Endangered (CR), Endangered (EN) and Vulnerable (VU). We utilized our MAR framework to predict genetic diversity losses using three different scaling factors, Dark red z=0.3, Mid red z=0.1, Light red z=0.05.

**Fig. S19.**
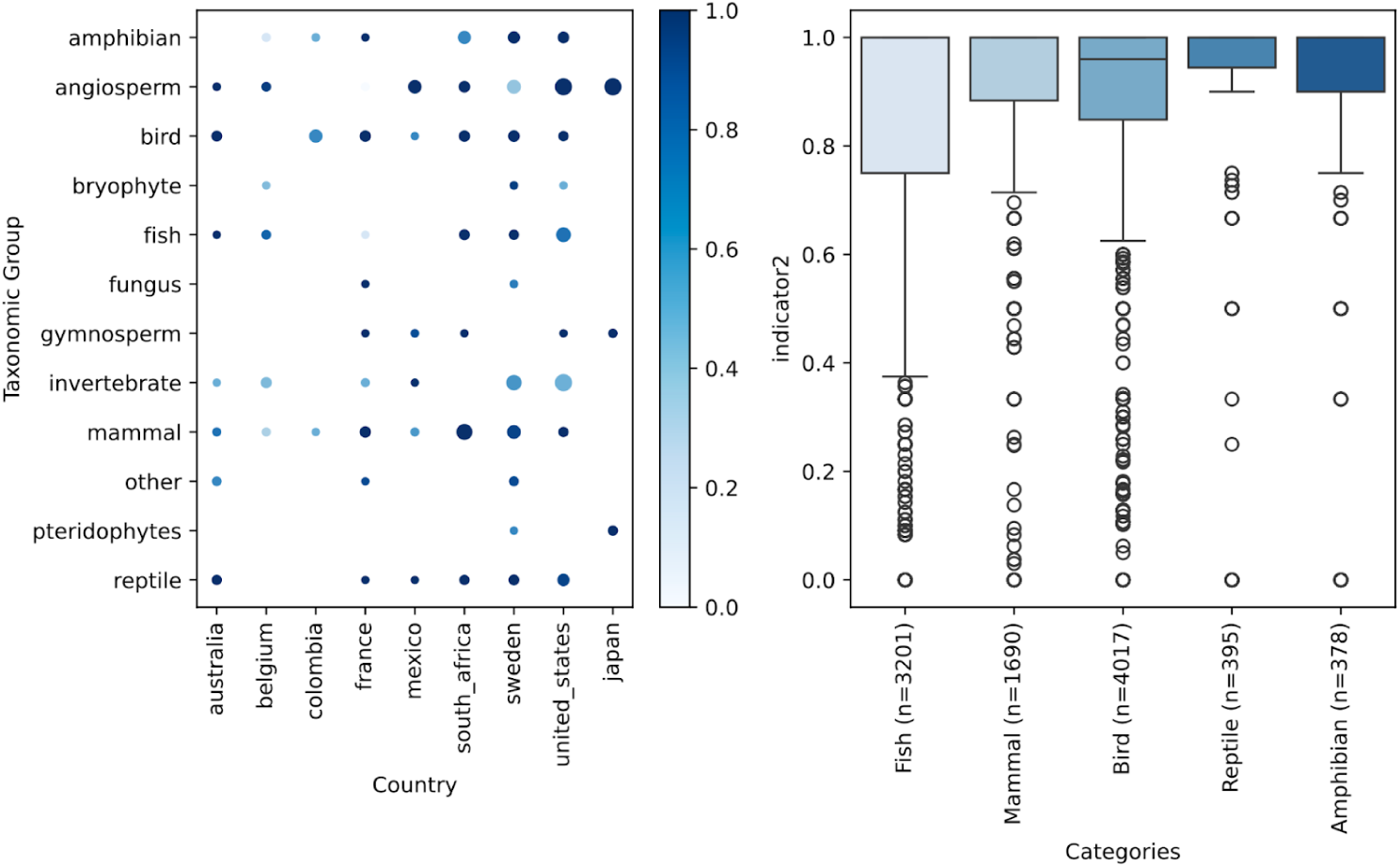
Summary of number of populations remaining from GBF indicator 2. We aggregate species collated from the Hoban et al. and LPI into 5-6 Taxonomic groups and plot the number of populations remaining (Indicator 2). (A) Data shown is from the paper Hoban et al. Size of dots indicate the number of species in that taxonomic group for that given country. Color represents the magnitude of Indicator 2. (B) Data shown is obtained from the Living Planet Index 2022 Database. Each point represents the indicator 2 value of a given species tracked in the LPI. Boxplots represent the distribution of Indicator 2 values across all species and all countries in that taxonomic group.

**Fig. S20.**
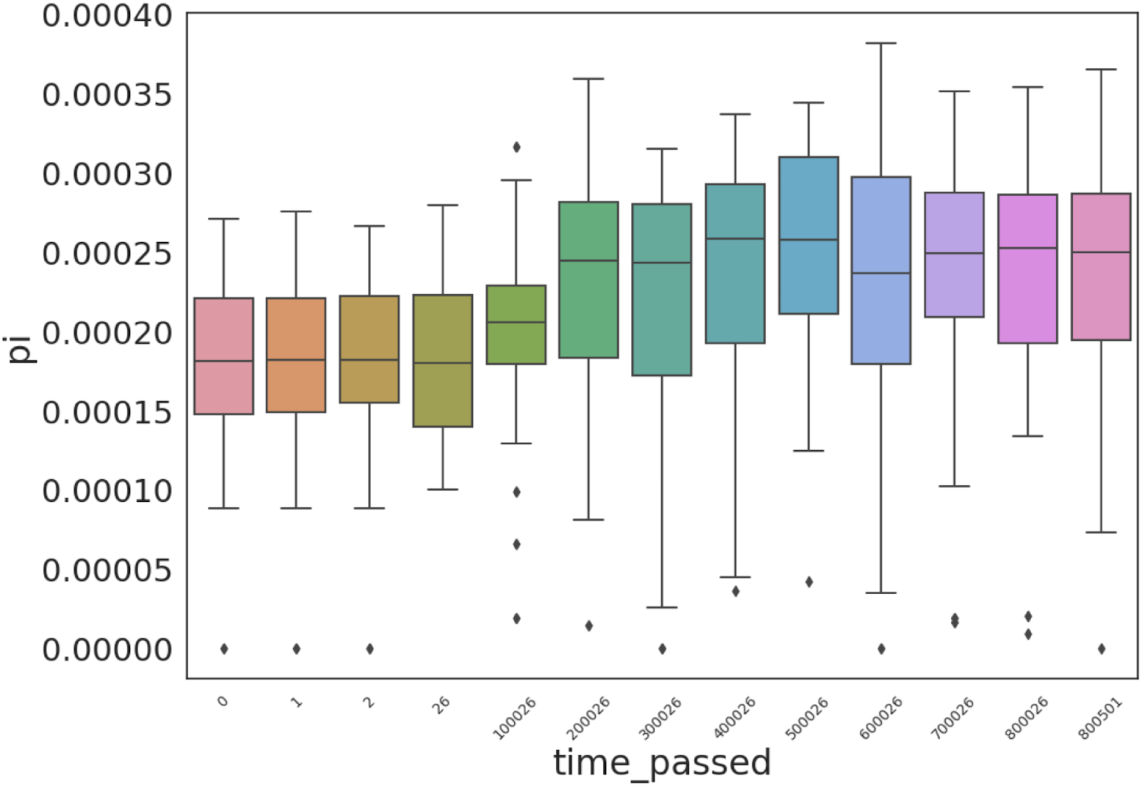
Genetic diversity (nucleotide diversity π) over burn-in time ensures stable initial conditions of area loss simulations. Population-specific π over 800,000 slim ticks. Box and whisker plots show the distribution of π over all populations in the habitat. Black dots represent outlier populations. Color gradient represents time passed. Burn-in simulations were performed for our population size of 5,000 with migration rate = 0.005 and an overall F_st_ of ∼0.3.

**Fig. S21.**
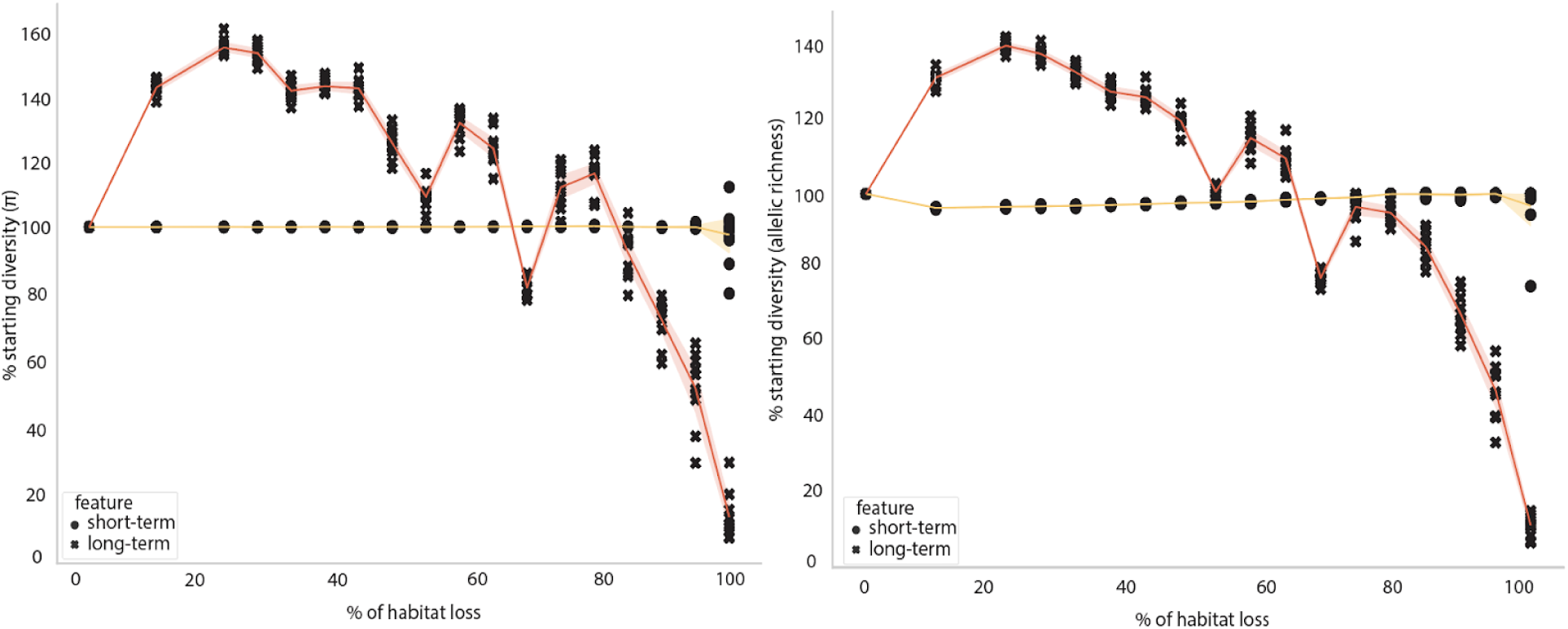
Genetic diversity (nucleotide diversity π and allelic richness *S*) across varying habitat loss with insufficient burn-in. Overlay of theoretical and simulation-based projections of short- and long-term genetic diversity loss, π, across different percentages of habitat loss from one edge. Simulations were run for 9 replicates. Red line shows average long-term genetic diversity projections. Yellow line shows average short-term genetic diversity projections. Shaded area showcases 95% confidence intervals across replicates.

**Fig. S22.**
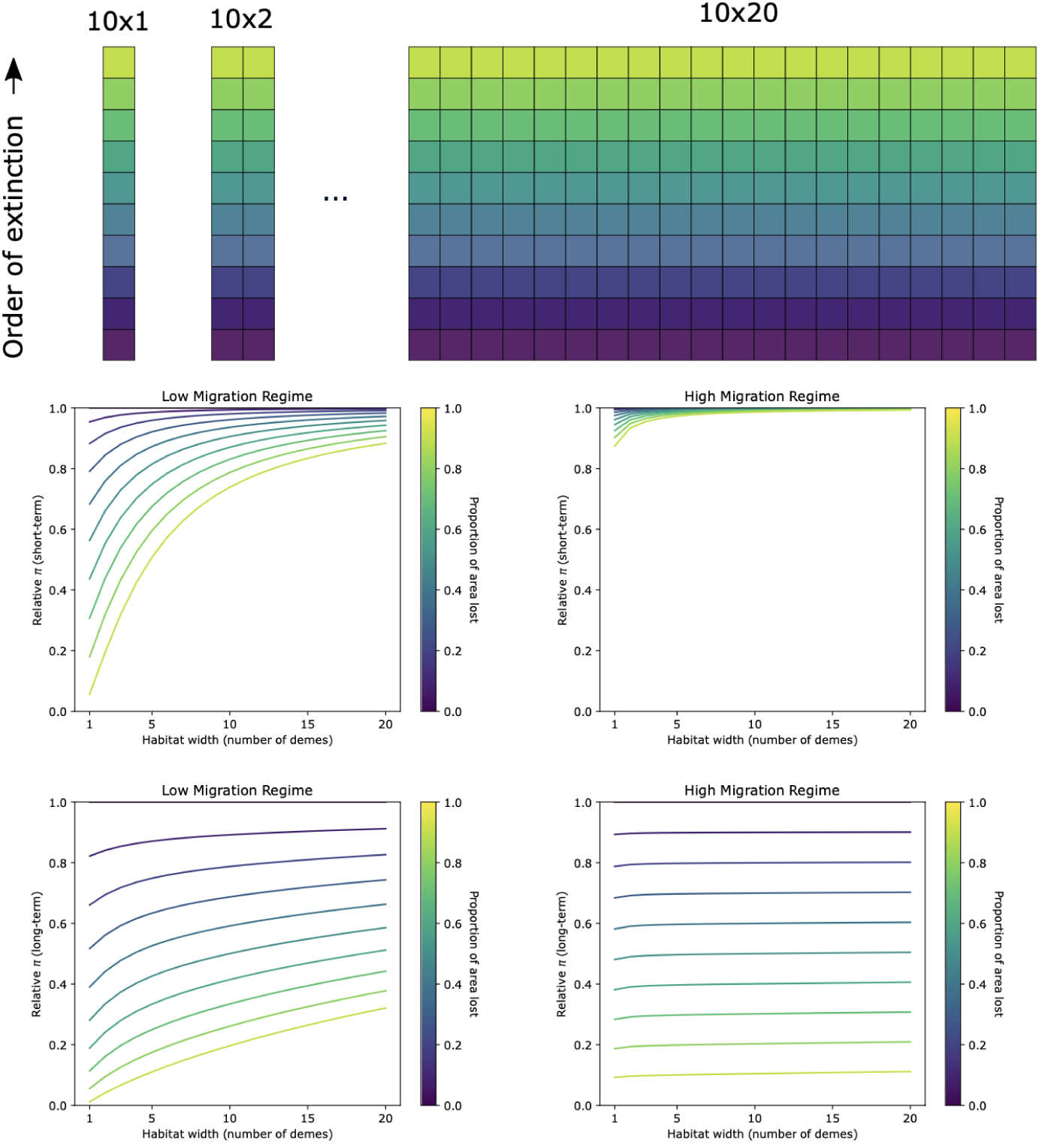
Genetic diversity trajectories across 1-D & 2-D habitats. Plots showing how genetic diversity in both short- and long-term change with geometry of habitat map and migration rate. Top row shows cartoons of habitat maps of varying sizes ranging from 10×1 to 10×20. Colors correspond to the amount of habitat loss incurred, with darker colors indicating lower losses while lighter colors indicating higher losses. Middle row shows short-term relative π across low and high migration regimes across different habitat widths. Bottom row shows long-term relative π across low and high migration regimes across different habitat widths. Each line represents a proportion of area loss across different habitat widths.

**Fig. S23.**
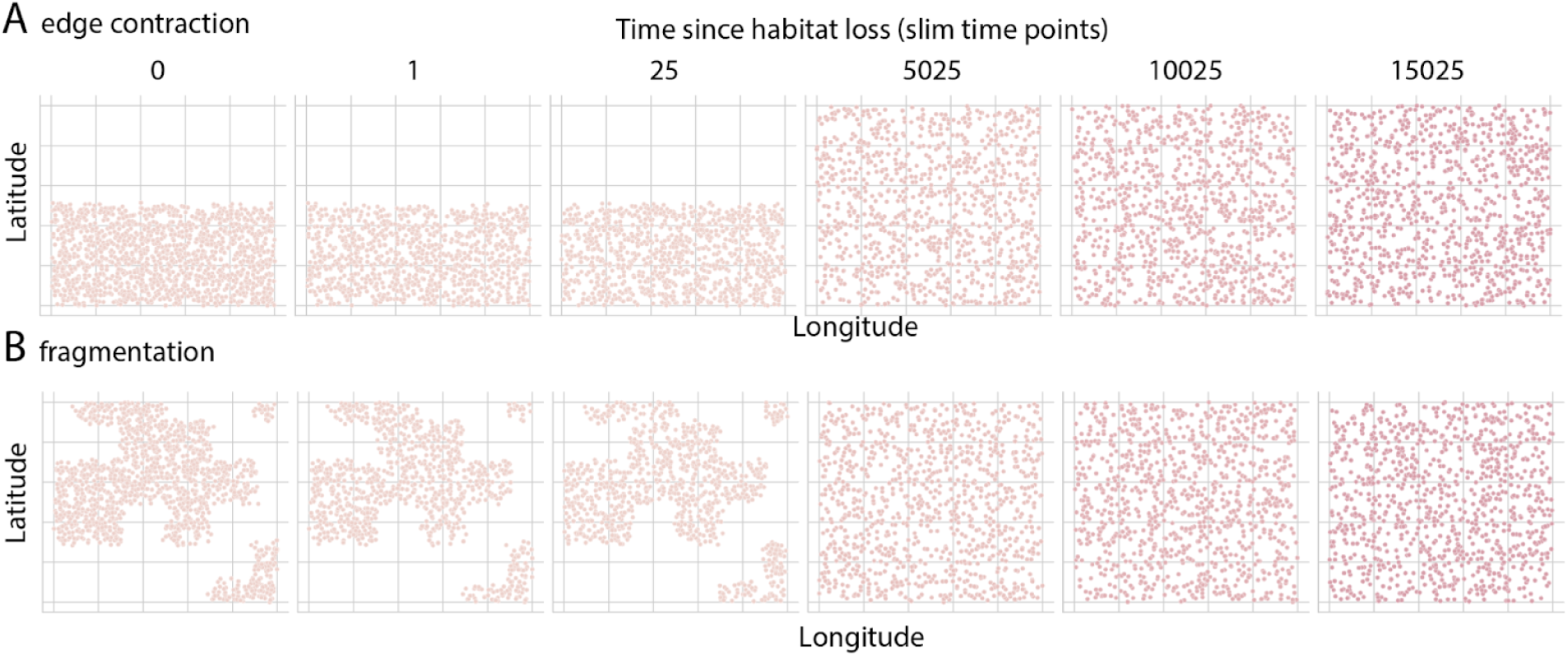
Habitat restoration maps. Cartoon showing habitat restoration within simulation space. (A) edge contraction (B) fragmentation maps. At time of habitat loss (Time=0), 50% of habitat loss is induced. Each dot represents an individual along a 2-D coordinate system (latitude, longitude). Empty boxes represent uninhabitable areas, individuals cannot disperse into these empty boxes.

**Fig. S24.**
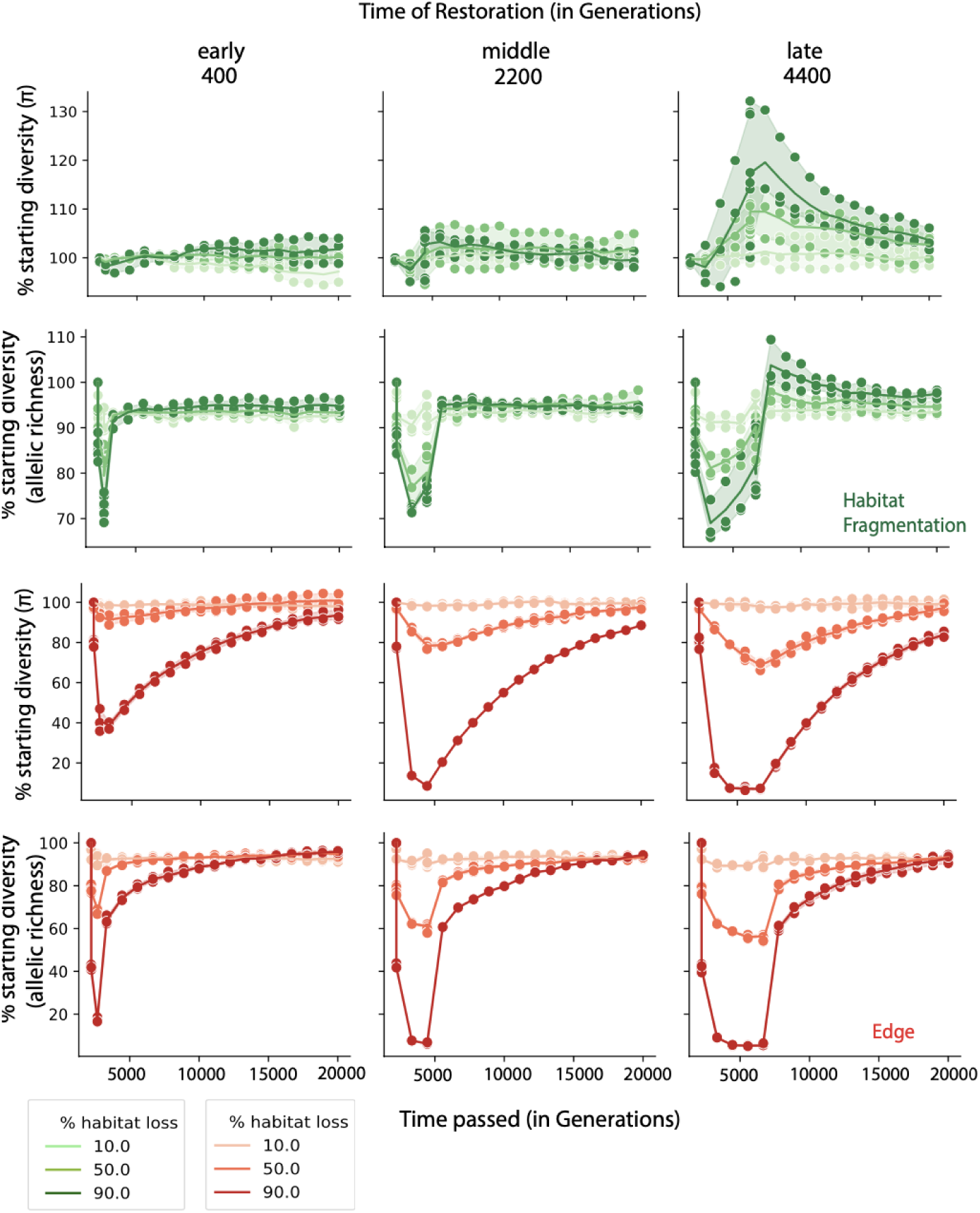
Genetic diversity metrics across habitat restoration maps. Genetic diversity metrics across habitat fragmentation (green) and edge contraction (red). Shades of green show % of habitat loss under habitat fragmentation and shades of red show % of habitat loss under edge contraction. Shown here are genetic diversity (π_species_) metrics, % π and % allelic richness across time in generations. We track genetic diversity until the next equilibrium is reached over 40000 generations. Time of restoration is illustrated and categorized into 3 categories of early (400), middle (2200) and late (4400). Each dot represents a genetic diversity measurement for % habitat loss at that specific time point. Shaded regions show the min-max across 3 replicates.

**Fig. S25.**
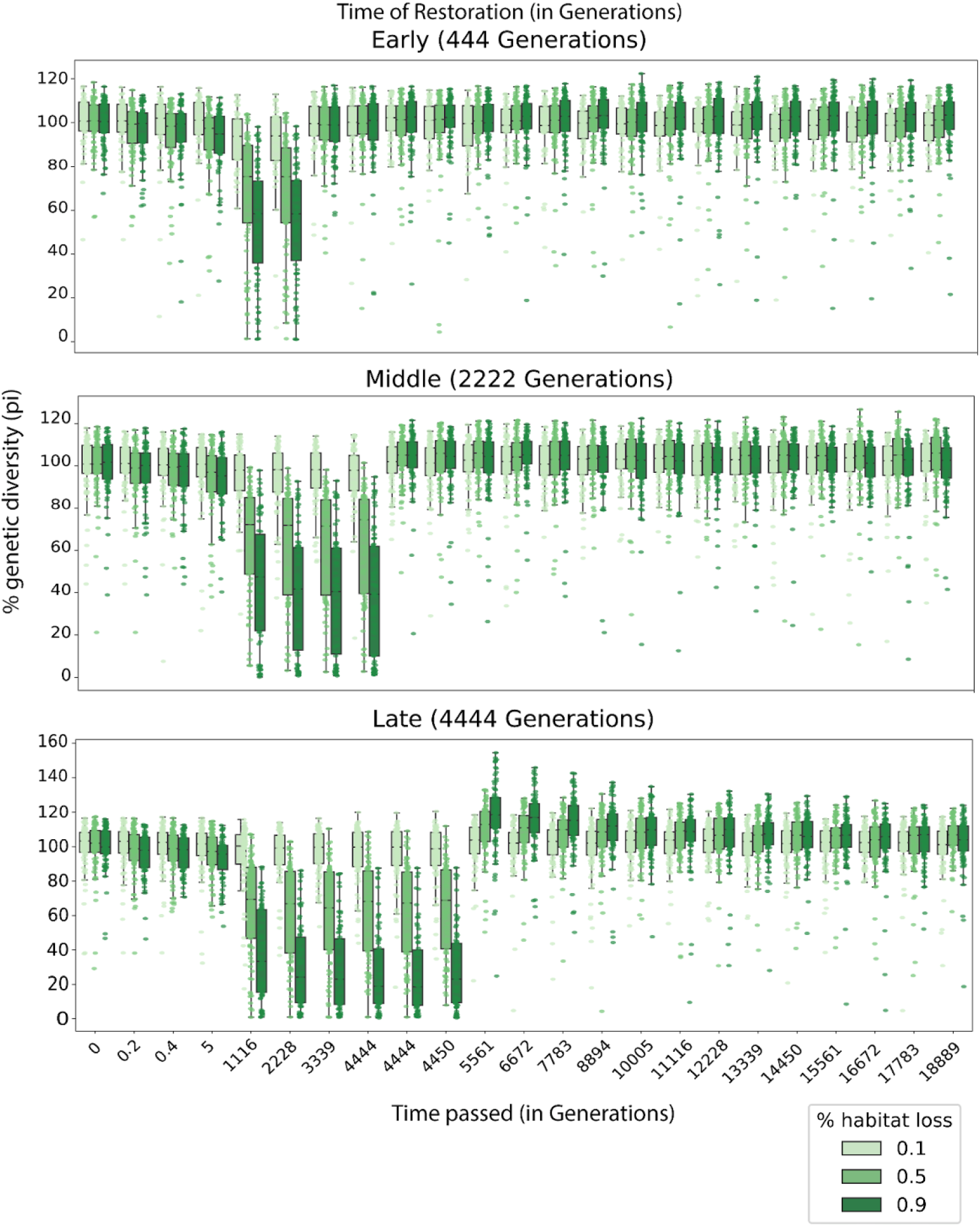
Within-population (π_local_) genetic diversity during restoration of habitat loss with fragmentation. Genetic diversity metrics across habitat fragmentation. Shades of green show % of habitat loss under habitat fragmentation. Shown here are genetic diversity (π_local_) metrics, % π across time in generations. We track genetic diversity until the next equilibrium is reached over ∼20000 generations. Time of restoration is illustrated and categorized into 3 categories of early (400), middle (2200) and late (4400). Each dot represents a measure of genetic diversity within a grid in a 10×10 simulation map. We performed these calculations across 3 replicates. Box and whisker plots show the distribution of species-wide (π_species_) over all grids in the habitat.

## Supplemental Tables

**Table S1.** SAR curve fit short-term genetic diversity (π) simulation trajectories under edge contraction. We fit 20 different functions and calculated variance explained (R^2^), Pearson r and Spearman *ρ*

| Model | $R^2$ | Pearson $r$ | Spearman $\rho$ |
| --- | --- | --- | --- |
| Asymptotic regression | 0.949 | 0.974 | 0.976 |
| Beta-P cumulative | 0.993 | 0.996 | 0.976 |
| Chapman Richards | 0.991 | 0.996 | 0.976 |
| Cumulative Weibull 3 par. | 0.992 | 0.996 | 0.976 |
| Cumulative Weibull 4 par. | 0.993 | 0.996 | 0.976 |
| Extended Power model 1 | 0.950 | 0.975 | 0.264 |
| Extended Power model 2 | 0.962 | 0.981 | 0.976 |
| Gompertz | 0.920 | 0.959 | 0.976 |
| Heleg(Logistic) | 0.993 | 0.996 | 0.976 |
| Kobayashi | 0.967 | 0.984 | 0.976 |
| Linear model | 0.422 | 0.650 | 0.976 |
| Logarithmic | 0.933 | 0.966 | 0.976 |
| Logistic(Standard) | 0.900 | 0.949 | 0.976 |
| Monod | 0.993 | 0.996 | 0.976 |
| Negative exponential | 0.949 | 0.974 | 0.976 |
| Persistence function 1 | 0.989 | 0.995 | 0.798 |
| Persistence function 2 | 0.993 | 0.996 | 0.976 |
| Power | 0.944 | 0.972 | 0.976 |
| PowerR | 0.964 | 0.982 | 0.976 |
| Rational function | 0.993 | 0.996 | 0.976 |

**Table S2.**
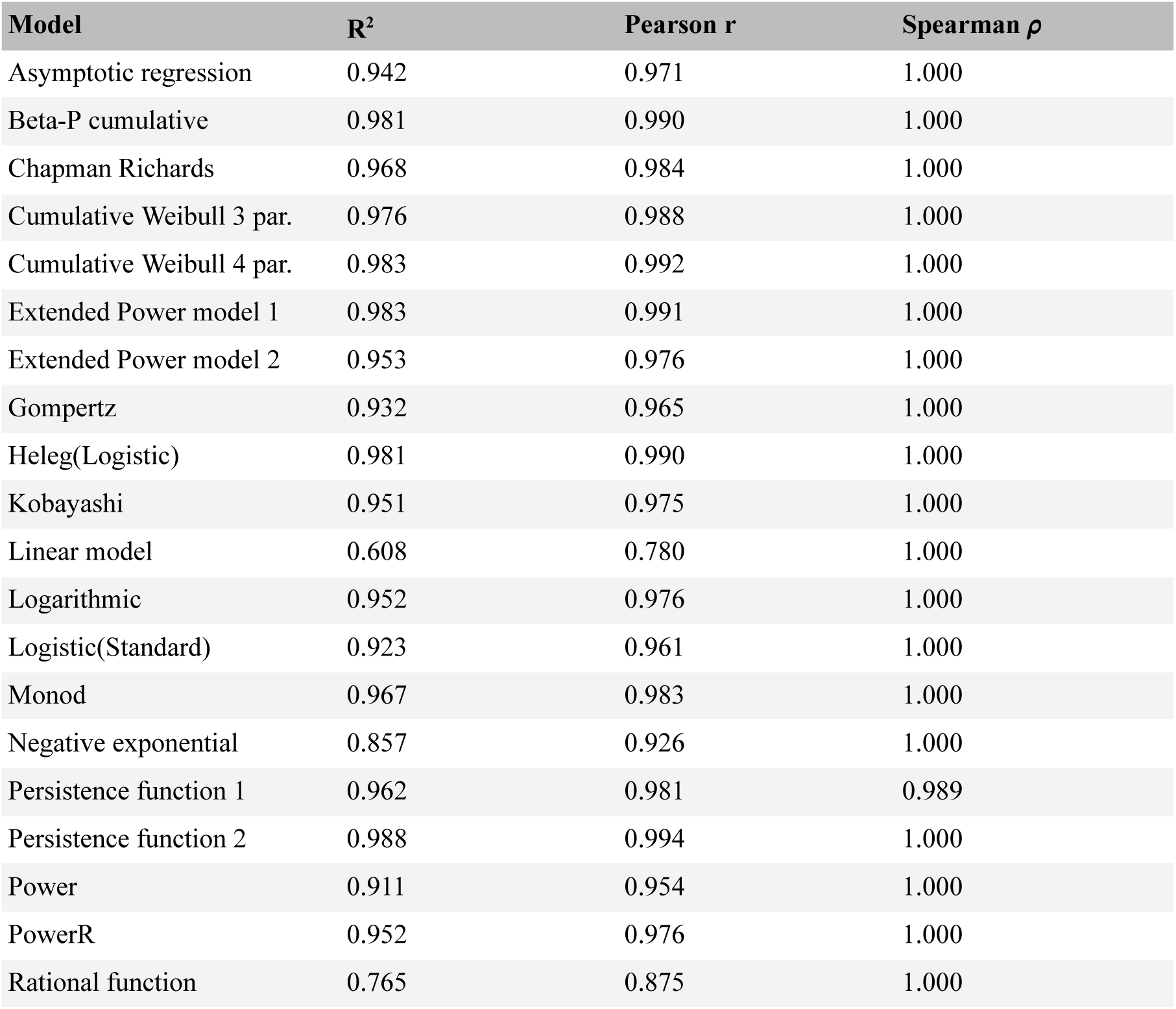
SAR curve fit to long-term genetic diversity (π) simulation trajectories under edge contraction. We fit 20 different functions and calculated variance explained (R^2^), Pearson r and Spearman *ρ*

| **Model** | **$R^2$** | **Pearson $r$** | **Spearman $\rho$** |
| --- | --- | --- | --- |
| Asymptotic regression | 0.942 | 0.971 | 1.000 |
| Beta-P cumulative | 0.981 | 0.990 | 1.000 |
| Chapman Richards | 0.968 | 0.984 | 1.000 |
| Cumulative Weibull 3 par. | 0.976 | 0.988 | 1.000 |
| Cumulative Weibull 4 par. | 0.983 | 0.992 | 1.000 |
| Extended Power model 1 | 0.983 | 0.991 | 1.000 |
| Extended Power model 2 | 0.953 | 0.976 | 1.000 |
| Gompertz | 0.932 | 0.965 | 1.000 |
| Heleg(Logistic) | 0.981 | 0.990 | 1.000 |
| Kobayashi | 0.951 | 0.975 | 1.000 |
| Linear model | 0.608 | 0.780 | 1.000 |
| Logarithmic | 0.952 | 0.976 | 1.000 |
| Logistic(Standard) | 0.923 | 0.961 | 1.000 |
| Monod | 0.967 | 0.983 | 1.000 |
| Negative exponential | 0.857 | 0.926 | 1.000 |
| Persistence function 1 | 0.962 | 0.981 | 0.989 |
| Persistence function 2 | 0.988 | 0.994 | 1.000 |
| Power | 0.911 | 0.954 | 1.000 |
| PowerR | 0.952 | 0.976 | 1.000 |
| Rational function | 0.765 | 0.875 | 1.000 |

**Table S3.** SAR curve fit to short-term genetic diversity (π) theoretical trajectories under edge contraction. We fit 20 different functions and calculated variance explained (R^2^), Pearson r and Spearman *ρ*

| **Model** | **$R^2$** | **Pearson $r$** | **Spearman $\rho$** |
| --- | --- | --- | --- |
| Asymptotic regression | 0.951 | 0.975 | 1.000 |
| Beta-P cumulative | 0.984 | 0.992 | 1.000 |
| Chapman Richards | 0.974 | 0.987 | 1.000 |
| Cumulative Weibull 3 par. | 0.980 | 0.990 | 1.000 |
| Cumulative Weibull 4 par. | 0.982 | 0.991 | 1.000 |
| Extended Power model 1 | 0.981 | 0.991 | 1.000 |
| Extended Power model 2 | 0.977 | 0.989 | 1.000 |
| Gompertz | 0.000 | 0.845 | 0.548 |
| Heleg(Logistic) | 0.982 | 0.991 | 1.000 |
| Kobayashi | 0.968 | 0.984 | 1.000 |
| Linear model | 0.782 | 0.884 | 1.000 |
| Logarithmic | 0.968 | 0.984 | 1.000 |
| Logistic(Standard) | 0.857 | 0.926 | 1.000 |
| Monod | 0.957 | 0.979 | 1.000 |
| Negative exponential | 0.796 | 0.892 | 1.000 |
| Persistence function 1 | 0.972 | 0.986 | 1.000 |
| Persistence function 2 | 0.987 | 0.994 | 1.000 |
| Power | 0.957 | 0.978 | 1.000 |
| PowerR | 0.000 | NA | NA |
| Rational function | 0.974 | 0.987 | 1.000 |

**Table S4.** SAR curve fit to long-term genetic diversity (π) theoretical trajectories under edge contraction. We fit 20 different functions and calculated variance explained (R^2^), Pearson r and Spearman *ρ*

| **Model** | **$R^2$** | **Pearson $r$** | **Spearman $\rho$** |
| --- | --- | --- | --- |
| Asymptotic regression | 0.995 | 0.997 | 1.000 |
| Beta-P cumulative | 0.994 | 0.997 | 1.000 |
| Chapman Richards | 0.000 | NA | NA |
| Cumulative Weibull 3 par. | 0.995 | 0.997 | 1.000 |
| Cumulative Weibull 4 par. | 0.994 | 0.997 | 1.000 |
| Extended Power model 1 | 0.995 | 0.998 | 1.000 |
| Extended Power model 2 | 0.995 | 0.998 | 1.000 |
| Gompertz | 0.807 | 0.898 | 0.894 |
| Heleg(Logistic) | 0.995 | 0.997 | 1.000 |
| Kobayashi | 0.994 | 0.997 | 1.000 |
| Linear model | 0.995 | 0.997 | 1.000 |
| Logarithmic | 0.883 | 0.940 | 1.000 |
| Logistic(Standard) | 0.988 | 0.994 | 1.000 |
| Monod | 0.994 | 0.997 | 1.000 |
| Negative exponential | 0.994 | 0.997 | 1.000 |
| Persistence function 1 | 0.995 | 0.997 | 1.000 |
| Persistence function 2 | 0.995 | 0.997 | 1.000 |
| Power | 0.995 | 0.997 | 1.000 |
| PowerR | 0.995 | 0.998 | 1.000 |
| Rational function | 0.995 | 0.997 | 1.000 |

**Table S5.** SAR curve fit to short-term genetic diversity (π) simulation trajectories under habitat fragmentation. We fit 20 different functions and calculated variance explained (R^2^), Pearson r and Spearman *ρ*

| **Model** | **$R^2$** | **Pearson $r$** | **Spearman $\rho$** |
| --- | --- | --- | --- |
| Asymptotic regression | $8.61 \times 10^{-4}$ | 0.029 | 0.041 |
| Beta-P cumulative | $9.24 \times 10^{-3}$ | 0.096 | 0.105 |
| Chapman Richards | $2.58 \times 10^{-3}$ | 0.051 | -0.041 |
| Cumulative Weibull 3 par. | $6.48 \times 10^{-5}$ | -0.008 | -0.041 |
| Cumulative Weibull 4 par. | $2.59 \times 10^{-5}$ | -0.005 | -0.041 |
| Extended Power model 1 | $4.73 \times 10^{-3}$ | 0.069 | 0.055 |
| Extended Power model 2 | $1.01 \times 10^{-5}$ | 0.003 | -0.041 |
| Gompertz | $9.24 \times 10^{-3}$ | 0.096 | 0.105 |
| Heleg(Logistic) | 0.00 | -0.008 | -0.041 |
| Kobayashi | $7.18 \times 10^{-5}$ | -0.008 | -0.041 |
| Linear model | $8.58 \times 10^{-4}$ | 0.029 | 0.041 |
| Logarithmic | $7.18 \times 10^{-5}$ | 0.008 | 0.041 |
| Logistic(Standard) | $3.47 \times 10^{-3}$ | 0.059 | 0.041 |
| Monod | $1.40 \times 10^{-4}$ | 0.012 | -0.041 |
| Negative exponential | $2.88 \times 10^{-3}$ | 0.054 | -0.041 |
| Persistence function 1 | $5.82 \times 10^{-3}$ | 0.076 | 0.072 |
| Persistence function 2 | $1.40 \times 10^{-4}$ | 0.012 | -0.041 |
| Power | $7.17 \times 10^{-5}$ | 0.008 | 0.041 |
| PowerR | 0.00 | NA | NA |
| Rational function | $1.06 \times 10^{-4}$ | 0.010 | -0.041 |

**Table S6.** SAR curve fit to long-term genetic diversity (π) simulation trajectories under habitat fragmentation. We fit 20 different functions and calculated variance explained (R^2^), Pearson r and Spearman *ρ*

| **Model** | **$R^2$** | **Pearson $r$** | **Spearman $\rho$** |
| --- | --- | --- | --- |
| Asymptotic regression | 0.493 | 0.702 | 0.620 |
| Beta-P cumulative | 0.516 | 0.719 | 0.620 |
| Chapman Richards | 0.000 | NA | NA |
| Cumulative Weibull 3 par. | 0.000 | -0.229 | -0.620 |
| Cumulative Weibull 4 par. | 0.000 | -0.540 | -0.620 |
| Extended Power model 1 | 0.522 | 0.723 | 0.685 |
| Extended Power model 2 | 0.263 | 0.513 | 0.620 |
| Gompertz | 0.496 | 0.704 | 0.620 |
| Heleg(Logistic) | 0.000 | NA | NA |
| Kobayashi | 0.376 | -0.613 | -0.620 |
| Linear model | 0.481 | 0.693 | 0.620 |
| Logarithmic | 0.376 | 0.613 | 0.620 |
| Logistic(Standard) | 0.000 | NA | NA |
| Monod | 0.000 | -0.470 | -0.620 |
| Negative exponential | 0.000 | NA | NA |
| Persistence function 1 | 0.526 | 0.725 | 0.685 |
| Persistence function 2 | 0.199 | 0.446 | 0.620 |
| Power | 0.345 | 0.587 | 0.620 |
| PowerR | 0.391 | 0.625 | 0.620 |
| Rational function | 0.444 | 0.666 | 0.620 |

**Table S7.** SAR curve fit to short-term genetic diversity (π) theoretical trajectories under habitat fragmentation. We fit 20 different functions and calculated variance explained (R^2^), Pearson r and Spearman *ρ*

| **Model** | **$R^2$** | **Pearson $r$** | **Spearman $\rho$** |
| --- | --- | --- | --- |
| Asymptotic regression | 0.496 | NA | NA |
| Beta-P cumulative | 0.496 | NA | NA |
| Chapman Richards | 0.000 | NA | NA |
| Cumulative Weibull 3 par. | 0.496 | NA | NA |
| Cumulative Weibull 4 par. | 0.496 | NA | NA |
| Extended Power model 1 | 0.496 | NA | NA |
| Extended Power model 2 | 0.496 | NA | NA |
| Gompertz | 0.498 | NA | NA |
| Heleg(Logistic) | 0.496 | NA | NA |
| Kobayashi | 0.496 | NA | NA |
| Linear model | 0.496 | NA | NA |
| Logarithmic | 0.495 | NA | NA |
| Logistic(Standard) | 0.496 | NA | NA |
| Monod | 0.496 | NA | NA |
| Negative exponential | 0.496 | NA | NA |
| Persistence function 1 | 0.496 | NA | NA |
| Persistence function 2 | 0.496 | NA | NA |
| Power | 0.496 | NA | NA |
| PowerR | 0.496 | NA | NA |
| Rational function | 0.496 | NA | NA |

**Table S8.** SAR curve fit to long-term genetic diversity (π) theoretical trajectories under habitat fragmentation. We fit 20 different functions and calculated variance explained (R^2^), Pearson r and Spearman *ρ*

| **Model** | **$R^2$** | **Pearson $r$** | **Spearman $\rho$** |
| --- | --- | --- | --- |
| Asymptotic regression | 0.149 | 0.386 | 0.325 |
| Beta-P cumulative | 0.001 | 0.027 | -0.232 |
| Chapman Richards | 0.002 | 0.046 | -0.066 |
| Cumulative Weibull 3 par. | 0.000 | -0.194 | -0.325 |
| Cumulative Weibull 4 par. | 0.000 | -0.313 | -0.325 |
| Extended Power model 1 | 0.167 | 0.408 | 0.347 |
| Extended Power model 2 | 0.063 | 0.251 | 0.325 |
| Gompertz | 0.000 | NA | NA |
| Heleg(Logistic) | 0.000 | NA | NA |
| Kobayashi | 0.098 | -0.313 | -0.325 |
| Linear model | 0.139 | 0.372 | 0.325 |
| Logarithmic | 0.098 | 0.313 | 0.325 |
| Logistic(Standard) | 0.002 | 0.046 | -0.059 |
| Monod | 0.000 | -0.216 | -0.325 |
| Negative exponential | 0.000 | 0.020 | -0.302 |
| Persistence function 1 | 0.169 | 0.411 | 0.350 |
| Persistence function 2 | 0.041 | 0.204 | 0.325 |
| Power | 0.091 | 0.301 | 0.325 |
| PowerR | 0.000 | NA | NA |
| Rational function | 0.128 | 0.358 | 0.325 |

**Table S9.** SAR curve fit to within-population short-term genetic diversity (π) simulation trajectories under habitat fragmentation. We fit 20 different functions and calculated variance explained (R^2^), Pearson r and Spearman *ρ*

| **Model** | **$R^2$** | **Pearson $r$** | **Spearman $\rho$** |
| --- | --- | --- | --- |
| Asymptotic regression | 0.004 | 0.066 | 0.062 |
| Beta-P cumulative | 0.009 | 0.096 | 0.062 |
| Chapman Richards | 0.009 | 0.096 | 0.062 |
| Cumulative Weibull 3 par. | 0.009 | 0.096 | 0.062 |
| Cumulative Weibull 4 par. | 0.007 | 0.081 | 0.062 |
| Extended Power model 1 | 0.009 | 0.096 | 0.062 |
| Extended Power model 2 | 0.008 | 0.089 | 0.062 |
| Gompertz | 0.000 | NA | NA |
| Heleg(Logistic) | 0.009 | 0.096 | 0.062 |
| Kobayashi | 0.007 | 0.081 | 0.062 |
| Linear model | 0.004 | 0.066 | 0.062 |
| Logarithmic | 0.007 | 0.081 | 0.062 |
| Logistic(Standard) | 0.004 | 0.066 | 0.062 |
| Monod | 0.009 | 0.093 | 0.062 |
| Negative exponential | 0.007 | 0.082 | 0.062 |
| Persistence function 1 | 0.009 | 0.096 | 0.068 |
| Persistence function 2 | 0.009 | 0.094 | 0.062 |
| Power | 0.007 | 0.081 | 0.062 |
| PowerR | 0.009 | 0.096 | 0.062 |
| Rational function | 0.009 | 0.093 | 0.062 |

**Table S10.** SAR curve fit to within-population long-term genetic diversity (π) simulation trajectories under habitat fragmentation. We fit 20 different functions and calculated variance explained (R^2^), Pearson r and Spearman *ρ*

| **Model** | **$R^2$** | **Pearson $r$** | **Spearman $\rho$** |
| --- | --- | --- | --- |
| Asymptotic regression | 0.398 | 0.631 | 0.629 |
| Beta-P cumulative | 0.419 | 0.647 | 0.629 |
| Chapman Richards | 0.418 | 0.647 | 0.629 |
| Cumulative Weibull 3 par. | 0.419 | 0.647 | 0.629 |
| Cumulative Weibull 4 par. | 0.414 | 0.644 | 0.629 |
| Extended Power model 1 | 0.418 | 0.647 | 0.629 |
| Extended Power model 2 | 0.389 | 0.623 | 0.629 |
| Gompertz | 0.334 | 0.578 | 0.587 |
| Heleg(Logistic) | 0.419 | 0.647 | 0.629 |
| Kobayashi | 0.386 | 0.621 | 0.629 |
| Linear model | 0.386 | 0.621 | 0.629 |
| Logarithmic | 0.393 | 0.627 | 0.629 |
| Logistic(Standard) | 0.419 | 0.647 | 0.629 |
| Monod | 0.386 | 0.621 | 0.629 |
| Negative exponential | 0.386 | 0.621 | 0.629 |
| Persistence function 1 | 0.001 | -0.029 | -0.629 |
| Persistence function 2 | 0.395 | 0.628 | 0.629 |
| Power | 0.373 | 0.611 | 0.629 |
| PowerR | 0.382 | 0.618 | 0.629 |
| Rational function | 0.397 | 0.630 | 0.629 |

**Table S11.** SAR curve fit to within-population short-term allelic richness (S) simulation predictions under habitat fragmentation on replicates with high connectivity. We fit 20 different functions and calculated variance explained (R^2^), Pearson r and Spearman *ρ*

| **Model** | **$R^2$** | **Pearson $r$** | **Spearman $\rho$** |
| --- | --- | --- | --- |
| Asymptotic regression | 0.938 | 0.968 | 0.949 |
| Beta-P cumulative | 0.936 | 0.967 | 0.949 |
| Chapman Richards | 0.937 | 0.968 | 0.949 |
| Cumulative Weibull 3 par. | 0.936 | 0.967 | 0.949 |
| Cumulative Weibull 4 par. | 0.938 | 0.968 | 0.949 |
| Extended Power model 1 | 0.936 | 0.968 | 0.949 |
| Extended Power model 2 | 0.936 | 0.967 | 0.949 |
| Gompertz | 0.000 | NA | NA |
| Heleg(Logistic) | 0.936 | 0.967 | 0.949 |
| Kobayashi | 0.934 | 0.966 | 0.949 |
| Linear model | 0.884 | 0.940 | 0.949 |
| Logarithmic | 0.934 | 0.967 | 0.949 |
| Logistic(Standard) | 0.936 | 0.967 | 0.949 |
| Monod | 0.928 | 0.963 | 0.949 |
| Negative exponential | 0.883 | 0.940 | 0.949 |
| Persistence function 1 | 0.937 | 0.968 | 0.949 |
| Persistence function 2 | 0.934 | 0.967 | 0.949 |
| Power | 0.930 | 0.964 | 0.949 |
| PowerR | 0.000 | NA | NA |
| Rational function | 0.937 | 0.968 | 0.949 |

**Table S12.**
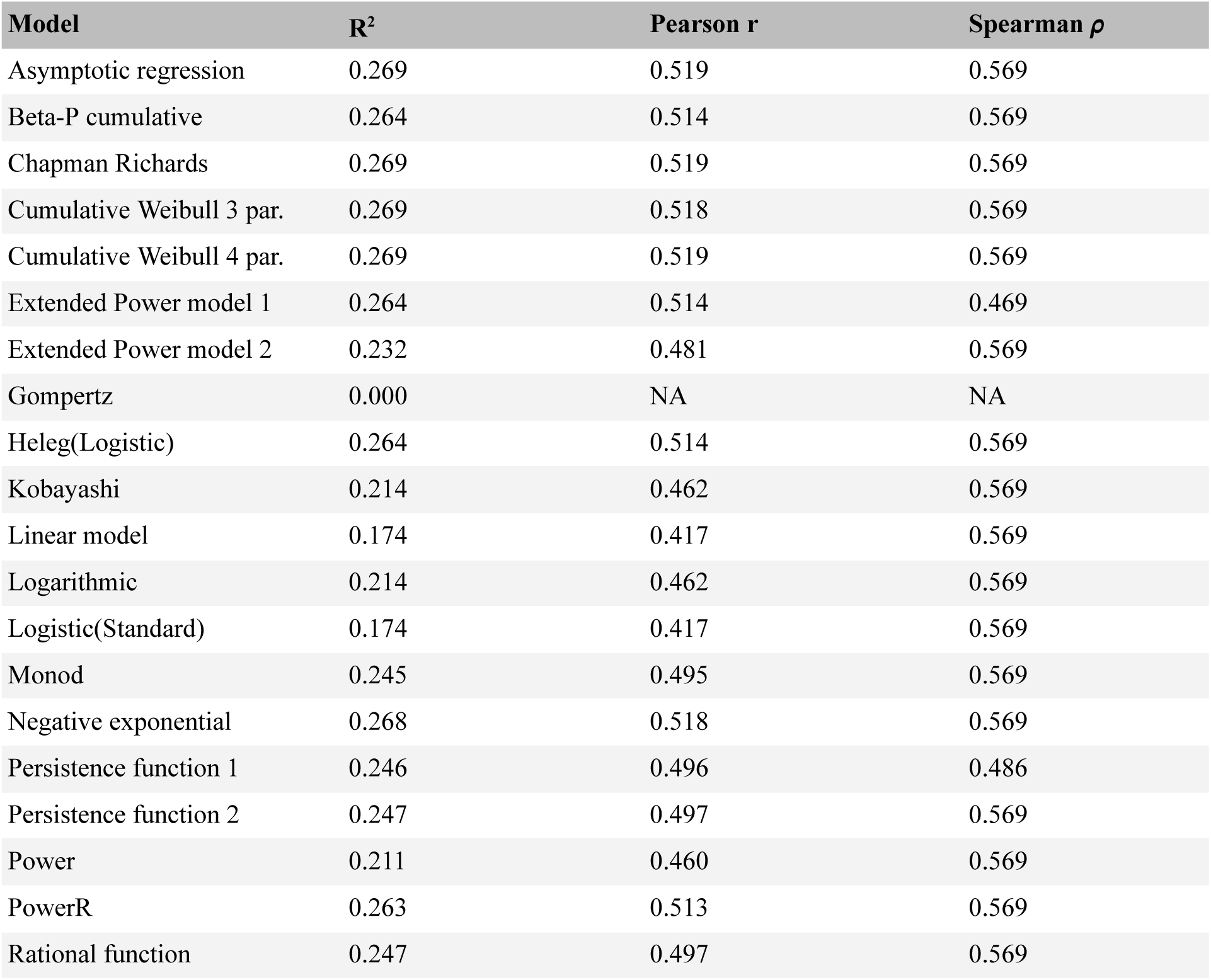
SAR curve fit to within-population long-term allelic richness (S) simulation predictions under habitat fragmentation on replicates with high connectivity. We fit 20 different functions and calculated variance explained (R^2^), Pearson r and Spearman *ρ*

**Table S13.** SAR curve fit to within-population short-term genetic diversity (π) simulation predictions under habitat fragmentation on replicates with high connectivity. We fit 20 different functions and calculated variance explained (R^2^), Pearson r and Spearman *ρ*

| **Model** | **$R^2$** | **Pearson $r$** | **Spearman $\rho$** |
| --- | --- | --- | --- |
| Asymptotic regression | 0.931 | 0.965 | 0.749 |
| Beta-P cumulative | 0.932 | 0.966 | 0.749 |
| Chapman Richards | 0.881 | 0.939 | 0.749 |
| Cumulative Weibull 3 par. | 0.932 | 0.966 | 0.749 |
| Cumulative Weibull 4 par. | 0.932 | 0.966 | 0.749 |
| Extended Power model 1 | 0.932 | 0.966 | 0.749 |
| Extended Power model 2 | 0.906 | 0.952 | 0.749 |
| Gompertz | 0.000 | NA | NA |
| Heleg(Logistic) | 0.932 | 0.966 | 0.749 |
| Kobayashi | 0.782 | 0.884 | 0.749 |
| Linear model | 0.458 | 0.677 | 0.749 |
| Logarithmic | 0.782 | 0.884 | 0.749 |
| Logistic(Standard) | 0.881 | 0.939 | 0.749 |
| Monod | 0.932 | 0.965 | 0.749 |
| Negative exponential | 0.881 | 0.939 | 0.749 |
| Persistence function 1 | 0.848 | 0.921 | 0.749 |
| Persistence function 2 | 0.932 | 0.965 | 0.749 |
| Power | 0.776 | 0.881 | 0.749 |
| PowerR | 0.000 | NA | NA |
| Rational function | 0.931 | 0.965 | 0.749 |

**Table S14.**
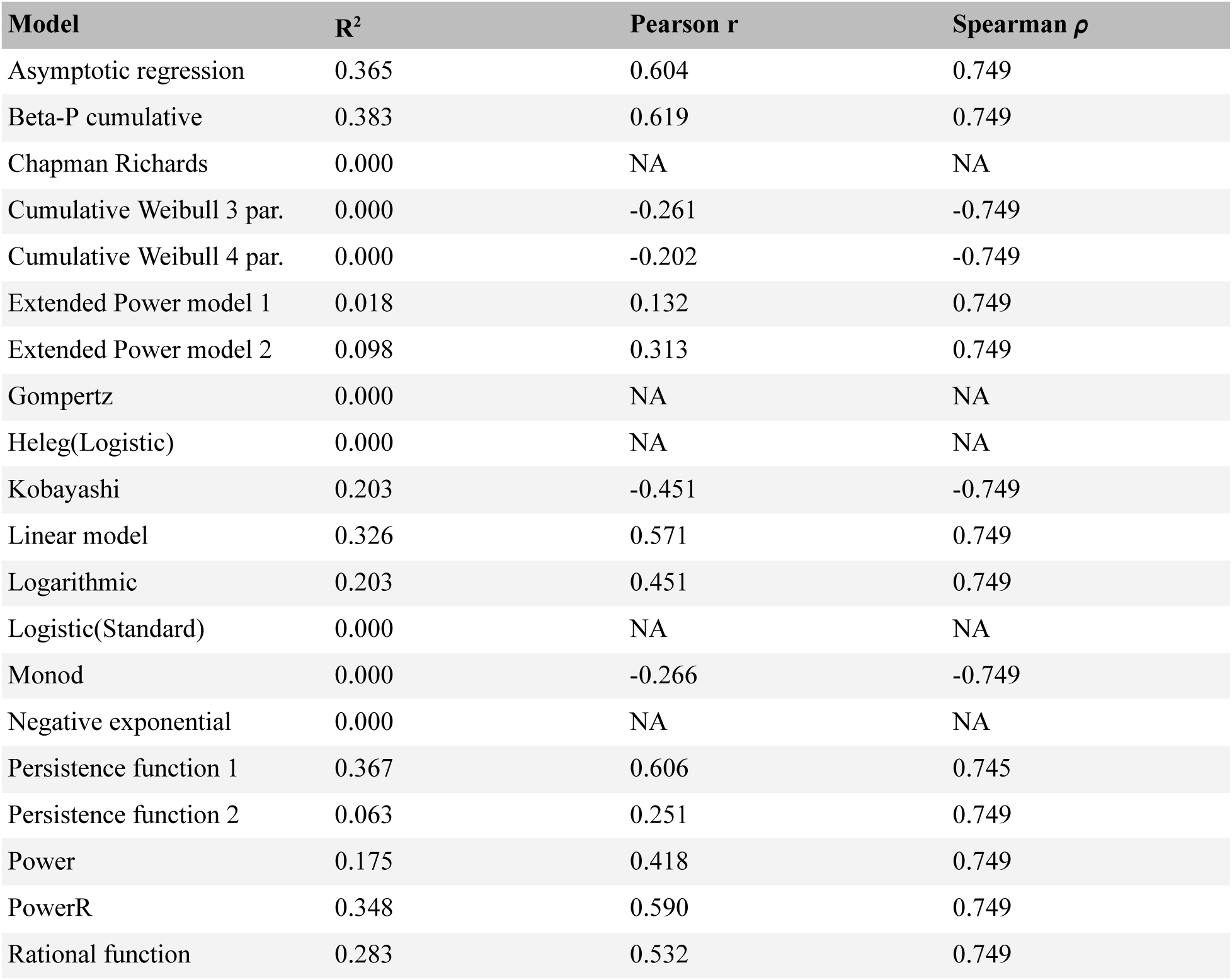
SAR curve fit to within-population long-term genetic diversity (π) simulation predictions under habitat fragmentation on replicates with high connectivity. We fit 20 different functions and calculated variance explained (R^2^), Pearson r and Spearman *ρ*

**Table S15.**
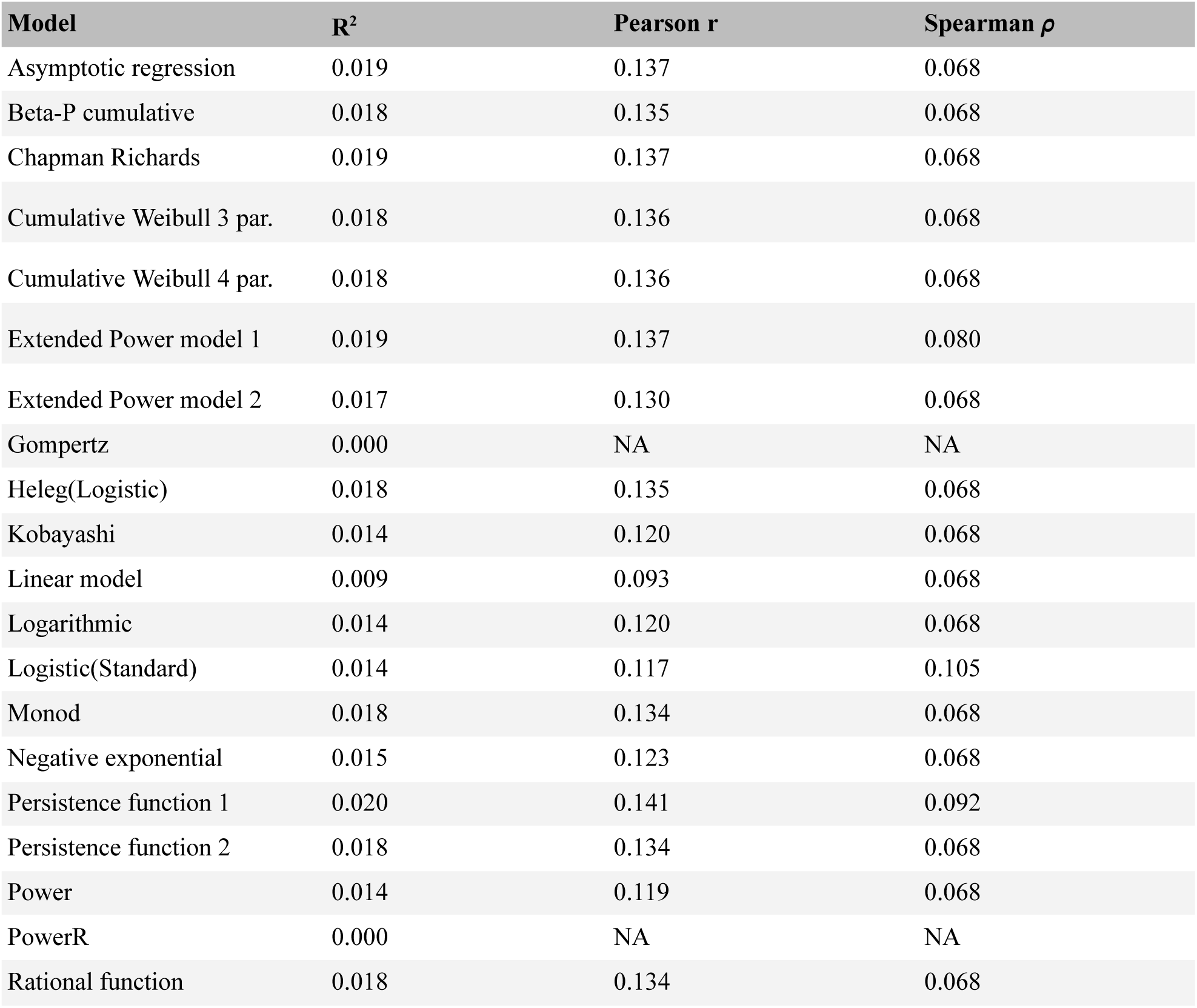
SAR curve fit to within-population short-term genetic diversity (π) simulation predictions in larger 20×20 habitat maps under habitat fragmentation. We fit 20 different functions and calculated variance explained (R^2^), Pearson r and Spearman *ρ*

**Table S16.** SAR curve fit to within-population long-term genetic diversity (π) simulation predictions in larger 20 x 20 habitat maps under habitat fragmentation. We fit 20 different functions and calculated variance explained (R^2^), Pearson r and Spearman *ρ*

| **Model** | **$R^2$** | **Pearson $r$** | **Spearman $\rho$** |
| --- | --- | --- | --- |
| Asymptotic regression | 0.273 | 0.523 | 0.600 |
| Beta-P cumulative | 0.277 | 0.526 | 0.600 |
| Chapman Richards | 0.276 | 0.525 | 0.600 |
| Cumulative Weibull 3 par. | 0.277 | 0.526 | 0.600 |
| Cumulative Weibull 4 par. | 0.278 | 0.527 | 0.600 |
| Extended Power model 1 | 0.274 | 0.523 | 0.600 |
| Extended Power model 2 | 0.273 | 0.523 | 0.600 |
| Gompertz | 0.245 | 0.495 | 0.584 |
| Heleg(Logistic) | 0.275 | 0.524 | 0.600 |
| Kobayashi | 0.271 | 0.521 | 0.600 |
| Linear model | 0.265 | 0.514 | 0.600 |
| Logarithmic | 0.262 | 0.512 | 0.600 |
| Logistic(Standard) | 0.281 | 0.530 | 0.600 |
| Monod | 0.272 | 0.521 | 0.600 |
| Negative exponential | 0.272 | 0.522 | 0.600 |
| Persistence function 1 | 0.278 | 0.527 | 0.600 |
| Persistence function 2 | 0.272 | 0.521 | 0.600 |
| Power | 0.269 | 0.518 | 0.600 |
| PowerR | 0.271 | 0.520 | 0.600 |
| Rational function | 0.273 | 0.522 | 0.600 |

**Table S17.**
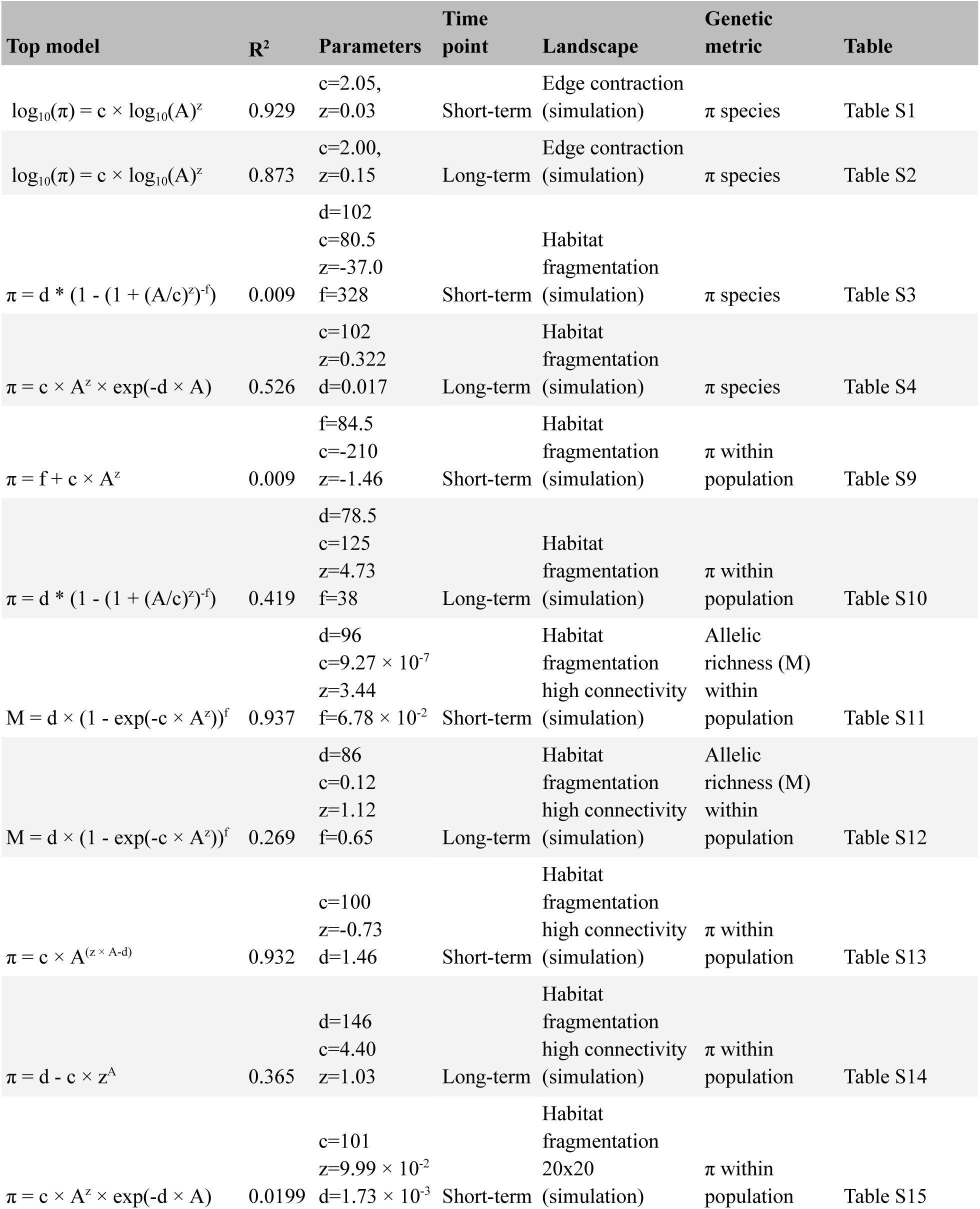
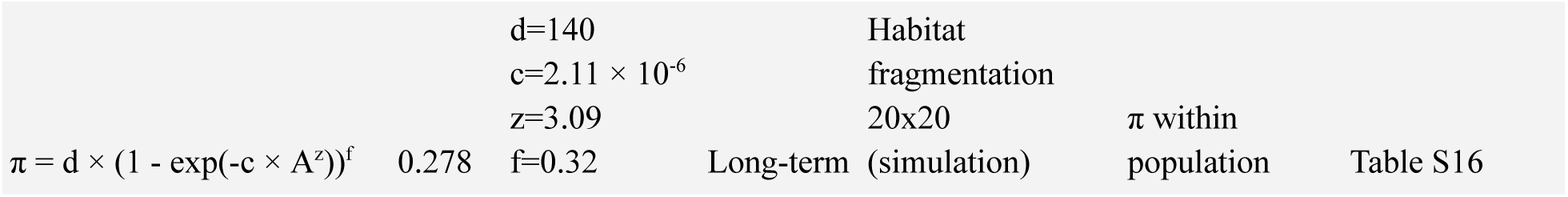
Genetic diversity and area relationship summaries for different landscapes and metrics. Fitted values in a log-log power law function between habitable area and genetic diversity (π)

**Table S18.**
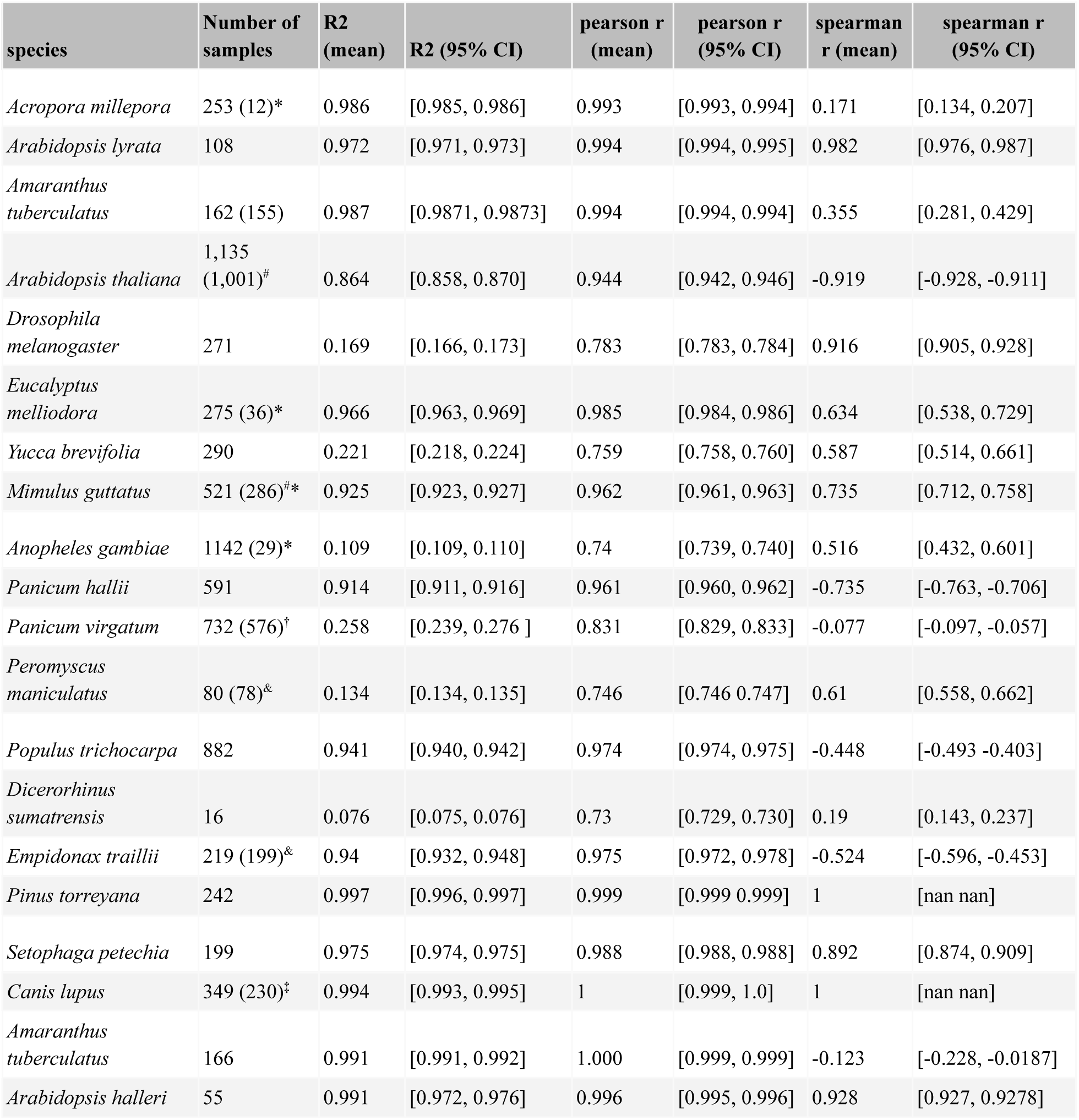
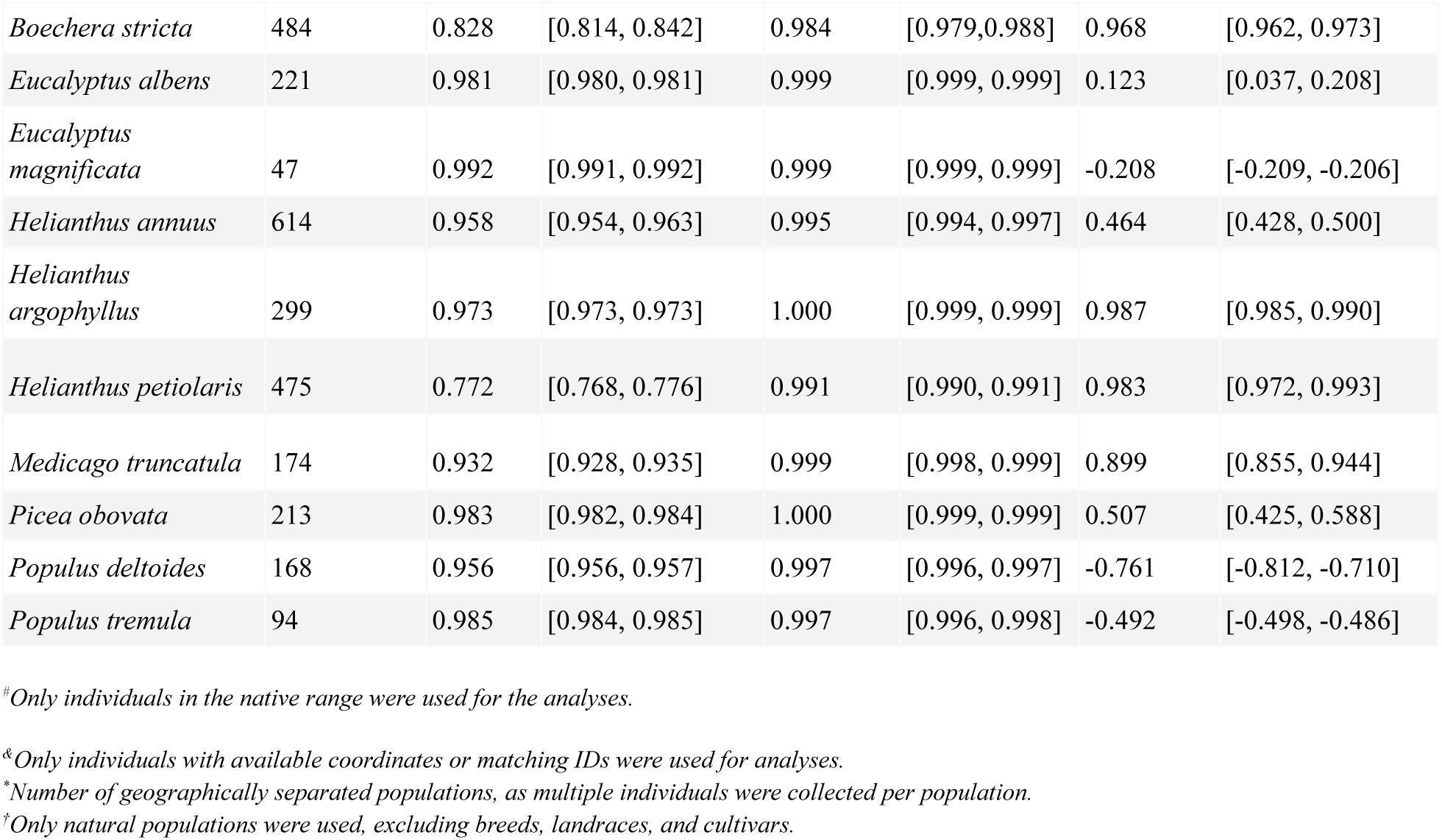
Genetic diversity and habitable area fit with 29 empirical species. Fitted values in a log-log power law function between area and genetic diversity (π) across short-term empirical simulations in 15 species. We fit the power law functions for short-term **(Fig S2)** and calculate variance explained (R^2^), Pearson r and Spearman rho.

**Table S19.** F_ST_ values across diverse species. F_ST_ values calculated using admixture R package (15) to estimate the most likely number of populations (K) tracked. *z_GDAR_* values using SAR R package (16) following the calculation of *z_MAR_* values in (6).

| Species | Publication | $F_{ST}$ avg | $F_{ST}$ max | $z_{GDAR}$ | $z_{MAR}$ |
| --- | --- | --- | --- | --- | --- |
| *Acropora millepora* | [10.1126/science.aba4674](https://doi.org/10.1126/science.aba4674) | 0.105 | 0.105 | 0.006 | 0.246 |
| *Amaranthus tuberculatus* | [10.1073/pnas.190087011](https://doi.org/10.1073/pnas.190087011) | 0.273 | 0.497 | -0.002 | 0.109 |
| *Anopheles gambiae* | [10.1038/nature24995](https://doi.org/10.1038/nature24995) | 0.321 | 0.484 | 0.044 | 0.214 |
| *Arabidopsis halleri* | [10.1371/journal.pgen.1005361](https://doi.org/10.1371/journal.pgen.1005361) | 0.669 | 0.935 | 0.160 | 0.299 |
| *Arabidopsis lyrata* | 10.1371/journal.pgen.1009477 | 0.137 | 0.137 | 0.018 | 0.236 |
| *Arabidopsis thaliana* | 10.1016/j.cell.2016.05.063 | 0.303 | 0.552 | 0.002 | 0.324 |
| *Boechera stricta* | [10.1186/s13059-019-1729-9](https://doi.org/10.1186/s13059-019-1729-9) | 0.229 | 0.384 | 0.035 | 0.054 |
| *Canis lupus* | [10.1111/mec.13364](https://doi.org/10.1111/mec.13364) | 0.092 | 0.092 | 0.064 | 0.256 |
| *Dicerorhinus sumatrensis* | 10.1038/s41467-021-22386-8 | 0.69 | 0.87 | -0.005 | 0.412 |
| *Drosophila melanogaster* | [10.1093/molbev/msab259](https://doi.org/10.1093/molbev/msab259) | 0.316 | 0.505 | 0.034 | 0.437 |
| *Empidonax traillii* | [10.1111/gcb.15639](https://doi.org/10.1111/gcb.15639) | 0.616 | 0.974 | 0.034 | 0.214 |
| *Eucalyptus albens* | [10.1111/mec.15287](https://doi.org/10.1111/mec.15287) | 0.039 | 0.039 | -0.006 | 0.037 |
| *Eucalyptus magnificata* | [10.1111/mec.15287](https://doi.org/10.1111/mec.15287) | 0.087 | 0.087 | -0.040 | NA |
| *Eucalyptus melliodora* | [10.7554/eLife.31835](https://doi.org/10.7554/eLife.31835) | 0.01 | 0.01 | 0.009 | 0.466 |
| *Helianthus annuus* | [10.1038/s41586-020-2467-6](https://doi.org/10.1038/s41586-020-2467-6) | 0.182 | 0.239 | 0.013 | 0.081 |
| *Helianthus argophyllus* | [10.1038/s41586-020-2467-6](https://doi.org/10.1038/s41586-020-2467-6) | 0.042 | 0.069 | 0.009 | 0.095 |
| *Helianthus petiolaris* | [10.1038/s41586-020-2467-6](https://doi.org/10.1038/s41586-020-2467-6) | 0.302 | 0.302 | 0.001 | 0.099 |
| *Medicago truncatula* | [10.1534/genetics.113.159319](https://doi.org/10.1534/genetics.113.159319) | 0.331 | 0.391 | 0.023 | 0.134 |
| *Mimulus guttatus* | [10.1038/s42003-021-01795-x](https://doi.org/10.1038/s42003-021-01795-x) | 0.372 | 0.613 | 0.018 | 0.274 |
| *Panicum hallii* | 10.1038/s41586-020-03127-1 | 0.238 | 0.369 | 0.455 | 0.824 |
| *Panicum virgatum* | [10.1111/mec.13467](https://doi.org/10.1111/mec.13467) | 0.53 | 0.82 | 0.009 | 0.232 |
| *Peromyscus maniculatus* | [10.1111/evo.13150](https://doi.org/10.1111/evo.13150) | 0.087 | 0.087 | 0.037 | 0.488 |
| *Picea obovata* | [10.1101/2023.01.31.526517](https://doi.org/10.1101/2023.01.31.526517) | 0.118 | 0.152 | 0.020 | 0.055 |
| *Pinus torreyana* | [10.1093/aobpla/plab058](https://doi.org/10.1093/aobpla/plab058) | 0.014 | 0.014 | 0.027 | 0.142 |
| *Populus deltoides* | [10.1002/ece3.3466](https://doi.org/10.1002/ece3.3466) | 0.097 | 0.097 | 0.006 | 0.072 |
| *Populus tremula* | [10.1186/s13059-018-1444-y](https://doi.org/10.1186/s13059-018-1444-y) | 0.058 | 0.058 | -0.005 | 0.087 |
| *Populus trichocarpa* | [10.13139/OLCF/1411410](https://doi.org/10.13139/OLCF/1411410) | 0.101 | 0.179 | 0.006 | 0.275 |
| *Setophaga petechia* | [10.1126/science.aan4380](https://doi.org/10.1126/science.aan4380) | 0.054 | 0.054 | 0.011 | 0.178 |
| *Yucca brevifolia* | [10.3732/ajb.1600069](https://doi.org/10.3732/ajb.1600069) | 0.395 | 0.716 | 0.018 | ×0.128 |

Abbreviations. ×Values excluded from global averages used for conservation applications due to uncertain estimates, suboptimal genomic data type.

**Table S20.**
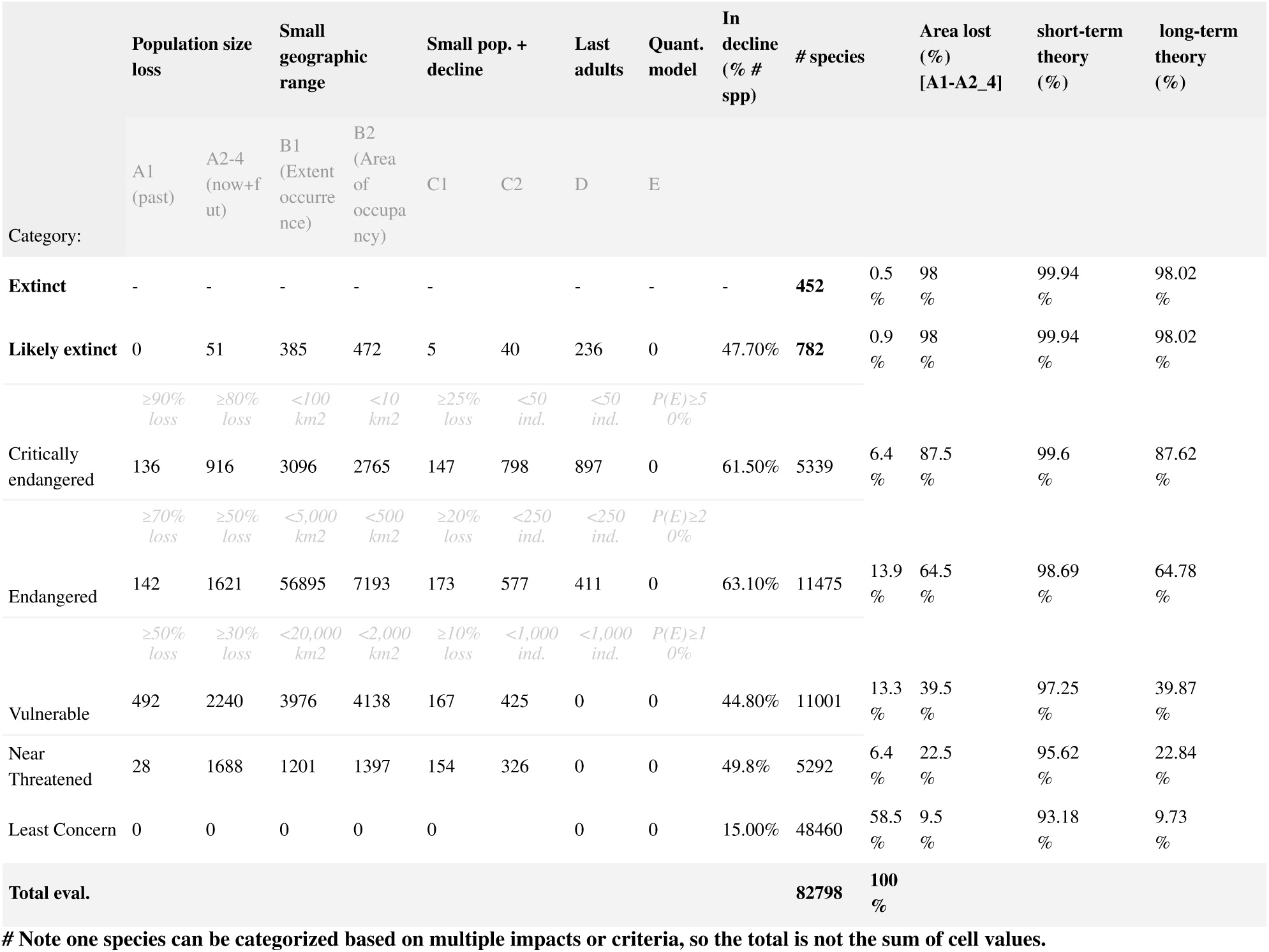
IUCN Red List area and population criteria for 80 thousand species. Each species was parsed for the indicator used to be classified in a given category. Summary of Red List database (www.iucnredlist.org). Counts of each category as well as criteria used in their classification are summarized (for details see extended guidelines). Area loss is obtained by using the Red List criteria for each category. Estimates of short and long-term genetic diversity loss were calculated using our theoretical and simulation-based framework.

## Notes

### Competing Interest Statement

The authors have declared no competing interest.

### Summary of Updates

Increased number of species analyzed. Included medium term projections. Fixed typos.

